# TimeFlow: a density-driven pseudotime method for flow cytometry data analysis

**DOI:** 10.1101/2025.02.16.638508

**Authors:** Margarita Liarou, Thomas Matthes, Stéphane Marchand-Maillet

## Abstract

Pseudotime methods order cells undergoing differentiation from the least to most differentiated. We developed TimeFlow, a new method for computing pseudotime in multi-dimensional flow cytometry datasets. TimeFlow tracks the differentiation path of each cell on a graph by following smooth changes in the cell population density. To compute the probability density function of the cells, it uses a normalizing flow model. We profiled bone marrow samples from three healthy patients using a 20-color antibody panel for flow cytometry and prepared datasets that ranged from 5,000 to 600,000 cells and included monocytes, neutrophils, erythrocytes and B-cells at various maturation stages. TimeFlow computed fine-grained pseudotime for all the datasets, and the cell orderings were consistent with prior knowledge of human hematopoiesis. Experiments showed its potential in generalizing across patients and unseen cell states. We compared our method to 11 other pseudotime methods using in-house and public datasets and found very good performance for both linear and branching trajectories. TimeFlow’s pseudotemporal orderings are useful for modelling the dynamics of cell surface proteins along linear trajectories. The biologically meaningful results in branching trajectories suggest the possibility of future applications with automated cell lineage detection. Code is available at https://github.com/MargaritaLiarou1/TimeFlow and bone marrow data will be accessible upon acceptance.

## 1 Introduction

Gene or protein expression changes as cells differentiate from a stem cell population into various mature cell types. Single-cell RNA sequencing (RNA-seq) [1] measures the expression levels of RNA molecules at a single cell level (expression profile) but destroys the cell upon its measurement. Consequently, RNA-seq cannot repeatedly measure the same cell to track changes in its expression in real time. Trajectory Inference (TI) methods assume that a cellular ”snapshot” taken at a specific timepoint captures single cells at all different states of the asynchronous differentiation process [2, 3]. TI methods compute the cell pseudotime, a unit free measure for the progression of a cell relative to the onset of differentiation [2]. Pseudotime values are used to order cells from the least to most mature along their differentiation pathway (cell lineage). This ordering is useful for modelling the evolution of a gene or protein along a cell lineage as a function of pseudotime, and further, studying the cell commitment towards different lineages [4].

More than 70 methods have been developed for the analysis of sparse single cell RNA-seq datasets with up to 20,000 genes and 10^2^ to 10^4^ cells [5]. Using tools from graph theory, principal manifold learning and stochastic processes [6], TI methods find linear trajectories with no branches [7, 8, 9], tree-shaped trajectories with multiple branches [10, 11, 12, 13], and cyclic trajectories or trajectories which may include disconnected segments [14, 15, 16, 17]. Adapting TI methods to flow cytometry datasets, which measure the expression of 10-50 cell surface proteins and usually contain 10^4^ to 10^6^, remains less explored [18, 19, 20]. The first TI method for cytometry, Wanderlust [21] resolves non-branching trajectories by representing cells as nodes in a set of k-Nearest Neighbour Graphs (k-NNGs) and orders them based on their shortest path Euclidean distance from the root node. Wishbone [22] extends Wanderlust to single-branched trajectories, while the work in [23] adapts it to tree-shaped trajectories but requires the user to manually distinguish mature and immature cell populations. CytoTree [24] organizes cells into clusters and connects their centres with a Minimum Spanning Tree (MST) to infer a branching trajectory backbone. We note that other important computational methods for cytometry such as SPADE [25], FlowSOM [26], FLOW-MAP [27] and TrackSOM [28] group cells along the branches of an MST, but do not explicitly target the single-cell pseudotime problem, and thus return no cell orderings. Aggregating differentiating cells into discrete clusters [7, 8, 9, 29, 30, 31], or k-NN graph communities [14, 15, 32]–two common practices in TI–contradicts the continuum of cell transitions present in nature. Lim et al. [33] showed in their recent work that clustering reduces the ”trajectory property” of the data and segments the differentiation pathways. Moreover, most available TI methods scale poorly with the number of cells [5]. We expect the scalability problem to be pronounced in flow cytometry datasets, which usually contain a significantly larger number of cells than RNA-seq datasets. For instance, CytoTree [24], recommends downsampling datasets that exceed 100,000 cells to one-tenth of the original size. To target the above challenges, we developed TimeFlow, a new pseudotime computation method for flow cytometry data with a single static snapshot. Our method tracks individual cell paths on a graph and differs from existing graph-based TI methods by using density-gradient graph weights.

## 2 Methods and Materials

### 2.1 Problem definition and TimeFlow

During its differentiation, a cell traverses a continuum of cell states, forming a path that starts from a stem cell population and ends in a mature, fully differentiated one. Our goal is to associate each cell with a pseudotime value in [0, 1] representing the cell’s progress along its path. The dataset is denoted as X = {x_i_}, where each cell state x_i_ ∈ X, the D-dimensional space of protein expression markers. The expression level for the m-th marker of the i-th cell is denoted as x_im_, with i ∈ [1*, …,* N] and m ∈ [1*, …,* D]. The dataset is a mixture of cell states, each captured at a different point in its evolution from the root—a stem cell identified by the user based on prior biological knowledge, e.g., expression of the surface marker CD34 in hematopoietic stem and progenitor cells (HSPCs)—to mature cell. We assume that we can reconstruct this evolution by finding a chain of cells for which

a. d (x_i_, x_j_) ≤ *ɛ*, for a small value of *ɛ*: the Euclidean distance between two consecutive cells in the chain is small.
b. |f_X_ (x_j_) − f_X_ (x_i_)|, the absolute value of the difference in probability density f_X_ between those cells is minimal.

These two criteria encourage cell transitions within isotropic local neighbourhoods that lead to smooth changes in the local density. Criterion (a) is common to several other TI methods as they assume that two cells close in differentiation share similar expression profiles [2], [23]. Criterion (b) is used here for the first time in a pseudotime computation method. We argue that both criteria follow naturally when the problem is recast in the language of the Optimal Transport (OT) theory [34, 35, 36, 37] (see details in Supplementary Section S1). We note that smooth changes in the cell density have also been reported (but not explicitly exploited as an algorithmic step) in other works [2, 21, 38]. Our method, TimeFlow, satisfies criteria (a) and (b) by taking the following steps:

1. Density estimation: estimate the probability density of cell states, f_X_, as a function of the marker values. The choice of the density estimation method is discussed in Sections 2.2 and 3.8.
2. Graph construction: construct a k-Nearest Neighbour (k-NN) graph to preserve the geometric locality. Each node corresponds to a cell state and edges connect states to their k nearest neighbours based on their Euclidean pairwise distances.
3. Graph weighting scheme: assign each edge (x_i_, x_j_) on the graph the absolute value of the difference of the probability density at the endpoints of the edge (density gradient), |f_X_ (x_j_) − f_X_ (x_i_)|.
4. Single-source shortest path tree: compute the shortest path from the root cell to every other cell on the weighted graph.
5. Pseudotime computation: sum the Euclidean distances between the nodes that comprise the shortest path of a cell. The pseudotime of a cell is this sum, after scaling the complete pseudotime distribution to [0, 1].

By (3), TimeFlow searches for transitions that are orthogonal to the density gradient over the k-NN graph. Thus, the pseudotime t_i_ of a cell x_i_ is inferred by measuring the markers’ evolution along the minimum density gradient path from the source to x_i_. Figure 1 is a schematic overview of TimeFlow.

**Figure 1:**
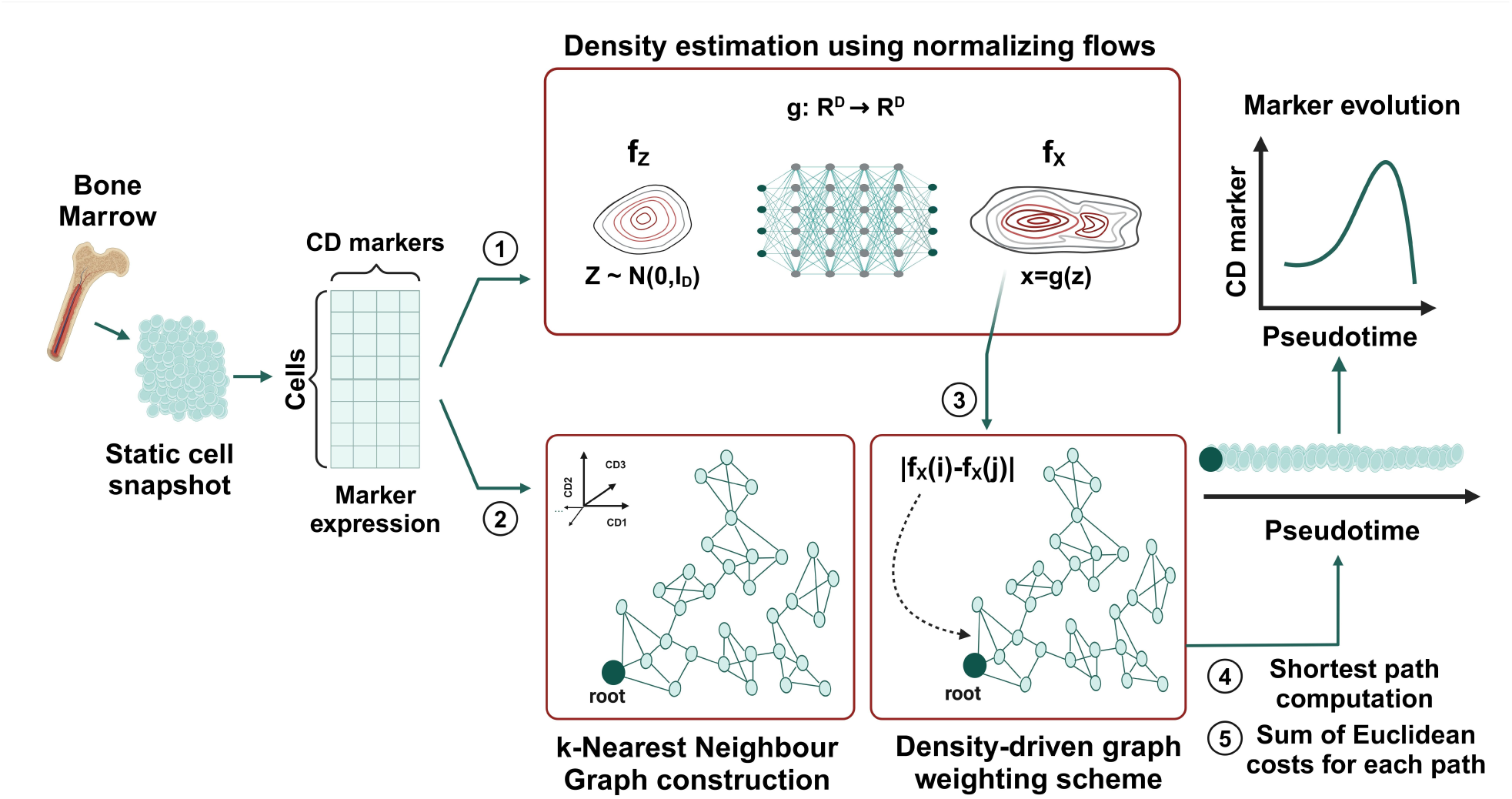
Schematic overview of TimeFlow. TimeFlow takes as input a root cell and a marker expression matrix, where each row corresponds to a cell and each column to a CD marker. It follows the next steps. Step 1: computes probability density of cell states, f_X_, using a normalizing flow model. Step 2: constructs a k-NN graph based on the cells’ pairwise Euclidean distances. Step 3: weights each edge (x_i_, x_j_) on the graph by density-gradient weights, |f_X_ (x_j_) − f_X_ (x_i_)|. Step 4: computes the shortest path of from the root to every other cell on the weighted graph. Step 5: calculates the pseudotime value of a cell by summing the Euclidean distances between the nodes along its path. It reorders the cells by increasing pseudotime value. If the trajectory is linear, a GAM model may be fitted to model the evolution of a marker as along pseudotime. The illustration was created using elements from BioRender.com.

### 2.2 TimeFlow uses the Real NVP transform for density estimation

TimeFlow includes an interchangeable density fitting module to estimate the cell density f_X_ of large cytometry datasets in 10 *<* D ≤ 50 dimensions. Kernel Density Estimation (KDE) [39, 40] has shown poor performance above three dimensions [41], and the kernel bandwidth selection is expensive for large datasets. An alternative for density estimation is normalizing flows [42, 43, 44], which unlike other generative models such as GANs [45] or VAEs [46], not only allow for the generation of new data samples, but also for exact evaluation of f_X_ (step (1) of TimeFlow). Normalizing flows model f_X_ through a base distribution f_Z_ of a random variable Z in R^D^ whose density is known (e.g., multivariate standard Gaussian *N* (z|0, I_D_). They require an invertible and differentiable function g_θ_ : R*^D^* → R*^D^* (bijector), with differentiable inverse g_θ_*^−^*^1^ : *R^D^* → *R^D^*, such that x = g_θ_(z) and z = g_θ_*^−^*^1^(x). Based on the change of variables formula, the target and base distributions are related as

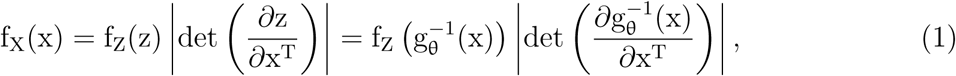

where 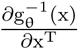 is the Jacobian matrix of g_θ_ ^−1^ evaluated at x. Function g_θ_ is usually implemented as a composition of L invertible functions g = g_L_ ◦· · · ◦g_1_ through a deep neural network architecture. TimeFlow uses a general-purpose, coupling-based flow model, the Real Non-Volume Preserving transform (Real NVP) [47], which has shown great performance for both inference and data generation tasks in different domains [48, 49, 50]. In the flow cytometry context, Real NVP has already proved its potential in [51], as part of a new single-cell annotation method. Real NVP stacks multiple linear transformations for high expressivity and reduces the cost of evaluating the Jacobian determinant from O (D^3^) to O (D) by exploiting the properties of triangular matrices. A more detailed description on the Real NVP transform is provided in Supplementary Section S2. For independent and identically distributed (i.i.d.) data, the model parameters θ are learnt by minimizing the average negative log likelihood function as shown in Supplementary Section S2 (eq.5). Details on the technical implementation of TimeFlow are given in Supplementary Section S3.

### 2.3 Evaluation of pseudotemporal orderings

Given the absence of precise ground truth, we compared the performance of TimeFlow to other methods using the gating labels (maturation stages), which associate each cell to its maturation stage as a weak ground truth. We calculated the Kendall’s Tau and Spearman’s Rank correlation coefficients between the inferred pseudotime of a cell and its maturation stage, represented as an integer that encodes its sequential order [10, 52] (e.g., immature cell: 1, intermediate cell: 2, . . .). Higher ordinal correlation indicates better alignment between pseudotime and the known stage ordering of the cells. We also computed the Cell Accuracy metric, which we defined as the proportion of cells correctly assigned to a pseudotime bin that corresponds to their maturation stage. For this, we sorted the cells by pseudotime and divided the pseudotime axis into as many bins as maturation stages. Assuming Q different stages with q in [1*, . . .,* Q], 1 corresponding to the least mature stage and Q to the most mature, the bins’ lower and upper ranges along the sorted cells are given as [1, n_1_), [n_1_, n_1_ + n_2_)*, . . .,* [n_1_ + n_2_ + *. . .* + n_Q_*_−_*_1_, n_1_ + n_2_ + *. . .* + n_Q_], where n_Q_ is the known total number of cells in stage q. Furthermore, we defined the Cell Stage Consistency (CSC) metric, which uses the Kendall’s Tau correlation to account for the constraints in the bin order. For the CSC, we labeled each bin with an integer that corresponds to the prevailing cell stage in the bin and created a new discrete ordering. The CSC score is equal to the Kendall’s Tau correlation between this discrete ordering and the weak ground truth. CSC scores range from -1 to 1, and values equal to 1 indicate that a method preserves the stage ordering and captures the general upward, downward, or stable trend of a marker along pseudotime. In addition, following the example of Palantir [12], we computed the Pearson Correlation Coefficient (PCC) between pseudotime and lineage commitment markers. These markers are often referred to as canonical markers in cytometry and are expected to monotonically increase within a specific cell lineage. High, positive values of PCC are indicative of biologically meaningful orderings. We selected the canonical markers CD14 and CD16, for monocytic and neutrophilic differentiation, respectively, as well as CD45, CD45RA, and CD19 for B-cells [53].

With a sufficiently large number of sampled cells (as is the case with our cytometry datasets) and given the continuous nature of hematopoiesis, we expect a well-performing method to produce a smooth and continuous trajectory. Resolving cell transitions at a fine scale is particularly useful for understanding intermediate states and marker evolution in less-studied differentiation processes. The pseudotime resolution affects the most critical downstream task of trajectory inference: modelling of the CD markers as a function of the inferred pseudotime. While a continuous pseudotime distribution allows for modelling the gradual variation in marker expression, a discrete pseudotime distribution may result in abrupt shifts and discontinuities in marker evolution, missing transitional states and complicating biological interpretation. Examples of coarsely resolved pseudotime distributions are provided in Section 3.2.2. To quantify the pseudotime resolution of a method across a trajectory, we computed the Shannon entropy of its pseudotime distribution as a measure of spread. We first sorted the cells of a cell lineage by pseudotime and then calculated the pseudotime histogram by dividing the cells into 100 equal-sized bins for sufficient resolution. We normalized the histogram counts into a probability distribution to compute its entropy and quantified its uncertainty using the formula H = − Σ_i_ p_i_ · log (p_i_), where p_i_, is the probability of the i-th bin. Higher entropy suggests that cells are spread across the bins and lower entropy implies that cells are concentrated within fewer bins. In the latter case, the distribution resembles more a sharp Dirac delta distribution, rather than a smooth pseudotime distribution, suitable for highly resolved cell transitions. Nevertheless, we stress that higher pseudotime resolution does not imply more accurate cell orderings. The pseudotime entropy serves only as a complementary way to assess whether a method yields fine- or coarse-grained orderings and should not be interpreted in terms of pseudotime accuracy, as also discussed in the Results section 3.2.2.

### 2.4 Datasets

#### 2.4.1 In house human bone marrow datasets

Our main application was the study of hematopoiesis. Hematopoiesis refers to the process in human bone marrow (BM), which leads to the production of all blood and immune cell types of the human body [54, 55, 56]. This is a well-studied process that serves as a good reference to understand the potential of a pseudotime estimation method. We focused our analysis on the generation of mature B-lymphocytes (B-cells), erythrocytes (Ery), monocytes (Mono) and neutrophils (Neu). We obtained BM samples from three patients (P1, P2, P3) and used a panel of 20 CD surface markers to define the different cell populations by flow cytometry (Supplementary Section S4.1). The patients had no hematological disease, and their samples were obtained during a pre-surgery control for hip replacement. For each BM sample, we followed the recently published gating strategy [57] and identified B-cell, monocytic, neutrophilic and erythrocytic populations from the least to most differentiated stages. We then arbitrarily subdivided each of these four populations into immature, intermediate, and mature stages and saved the gated populations as separate CSV files. Hence, we obtained four linear trajectories per patient to test TimeFlow on each trajectory independently of the others. In addition, we merged the four populations of each patient into new separate CSV files, creating the datasets, P1-BM, P2-BM, and P3-BM, which contained a branching trajectory of four differentiation pathways each. The datasets ranged from 5,000 to 600,000 cells. Cell counts for each population are summarized in Supplementary Table S1.

#### 2.4.2 Public mass cytometry datasets

We obtained other relevant flow and mass cytometry public datasets from the literature.

##### Human bone marrow mass cytometry datasets

We downloaded from the R package “HD-CytoData” [58], the Levine-13 and Levine-32 mass cytometry datasets, first introduced in [59] and [60], and used to benchmark clustering methods in [61]. We believe these datasets are suitable for pseudotime analysis because they describe a branching trajectory in BM and contain populations such as B-cells and monocytes at various maturation stages. Levine-13 measures the expression of 13 CD markers on 167,044 cells and Levine-32 integrates 265,627 cells from the BM of two healthy patients and measures their expression for 32 CD markers. This allowed us to test how TimeFlow handles data with a larger number of dimensions compared to conventional flow cytometry. More details on these datasets are given in Supplementary Sections S4.2, S4.3 and Supplementary Tables S2-S3.

##### Mouse thymus mass cytometry dataset

We chose the mass cytometry dataset from Wishbone [25], which characterizes T cell development in the mouse thymus and contains 25,000 cells measured for 13 CD markers. This process includes one branching decision where lymphoid progenitors separate into CD8^+^ cytotoxic (CD8^+^ SP) and CD4^+^ helper T cells (CD4^+^ SP). More details are given in Supplementary Section S4.4 and Supplementary Table S4.

We describe all the pre-processing steps applied on each dataset in Supplementary Section S5.

## 3 Results

To understand the potential of our method we used all the previously described in-house and public datasets. After computing the cell pseudotime for linear trajectories, we reordered the cells and fitted a Generalized Additive Model (GAM) [62] with a cubic spline. In this way, we modelled the evolution of markers in linear trajectories as a nonlinear function of pseudotime, without using cell labels. We did not follow the same process for branching trajectories because TimeFlow does not yet automatically identify cell lineages.

### 3.1 TimeFlow on hematopoietic linear and branching trajectories with flow cytometry data

We started by testing TimeFlow on linear trajectories using the P1-Mono, P1-Neu, P1-Ery and P1-B cells datasets. Figure 2A-D presents examples of marker dynamics along linear trajectories. In the P1-Mono dataset, cells at early pseudotime correspond to immature monocytes and exhibit minimal CD14 expression, while cells at intermediate pseudotime show a gradual increase in CD14 expression, peaking in mature cells. Similarly, in the P1-Neu dataset, we observed increasing expression in CD16 along pseudotime, as expected (Section 2.3). In the P1-Ery dataset, we observed an initial increase of CD36 in cells at early pseudotime, followed by a progressive decrease in cells at later pseudotime. This is a characteristic nonlinear pattern during erythropoiesis [63]. In line with prior biological knowledge of B cell maturation, CD45 expression increased monotonically as cells progressed towards the mature B-cell stage. Figure 2E-H presents stacked bar plots showing the distribution of cells (sorted by pseudotime) across the known maturation stages. In spite of the observed overlap in some intermediate cell states, TimeFlow reproduced the expected cell stage transitions and expression patterns. Supplementary Figure S1 illustrates some of these patterns (CD34, CD133, CD64, CD36, CD15, CD45RA, and CD19) for linear trajectories.

**Figure 2:**
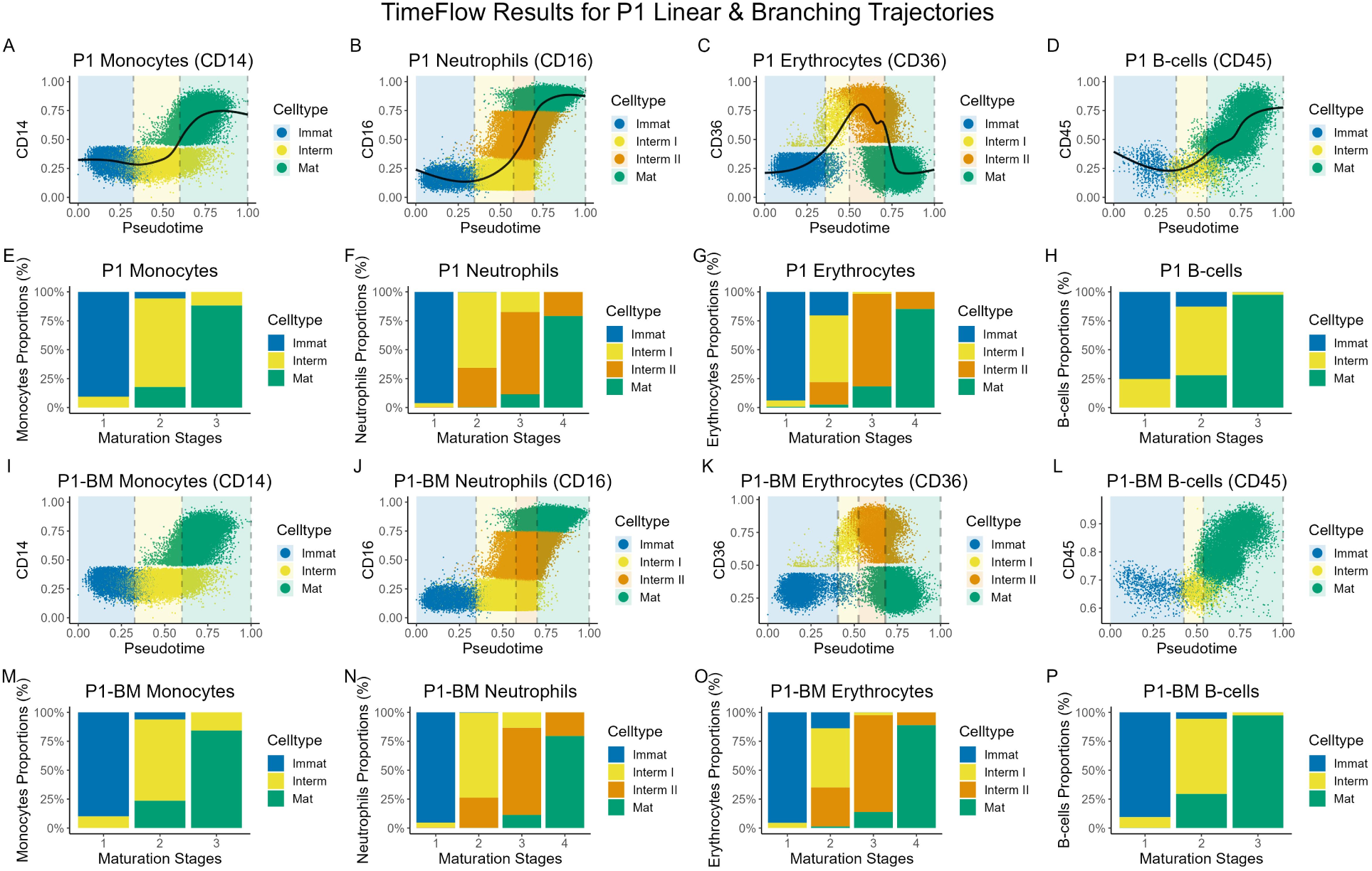
TimeFlow on linear (A-H) and branching (I-P) trajectories from P1 datasets. Each scatterplot dot represents a cell, coloured by its maturation stage. The marker expression was scaled in [0,1]. For linear trajectories, a GAM model (solid black curve) was fitted without use of labels. The vertical dashed grey lines indicate the boundaries between the cell stages. For each cell stage, the cut-off is defined at the pseudotime value corresponding to the last cell in that stage’s ordered sequence (cumulative cell counts, see Section 2.3). Boundaries might overlap if a method does not separate between distinct stages. Regions correspond to different cell stages and are colored such that the transitions between them are highlighted. Stacked bar plots show the cell distribution (%) in each of their known maturation stages, following pseudotime ordering. The stacked barplots cut-offs are the same with the cut-offs on the pseudotime axis. Gaps in the marker expression (y-axis) between consecutive cell populations are a consequence of using rectangular gates during dataset preparation.

We used the combined P1-BM dataset to test whether TimeFlow produces biologically plausible orderings in the presence of multiple cell lineages. As indicated by Figure 2I-L, most cells were ordered in accordance with their known maturation stage. Stacked bar plots in Figure 2M-P resembled the outcomes from the linear trajectories, with minor improvements in P1-BM Neu and minor deteriorations in P1-BM Ery. Supplementary Figures S2-S5 present the scatterplot of each marker against the inferred pseudotime for each cell lineage within the P1-BM dataset. The meaningful pseudotemporal orderings in the context of diverse cell types suggest the possibility of a lineage detection strategy that would build upon these orderings (see Section 4.1).

### 3.2 Comparison of TimeFlow to other methods

We compared the performance of TimeFlow to 11 other established TI methods. We used ElPiGraph, PAGA, Palantir and TinGa, which are designed for complex trajectories, in all in-house and public branching datasets. We evaluated Comp1, DPT and ElPiLinear on the P1/2/3-Mono/Neu/Ery/Bcells linear trajectories, but limited the comparisons between Wanderlust, SCORPIUS, scShaper and CytoTree to the smaller linear trajectories of P1/2/3-B-cells/Mono/Ery, because not all these methods returned results for all three large neutrophilic datasets within a reasonable time (18h on a modern computer). Supplementary Section S6 briefly describes the above methods, and Supplementary Table S5 provides a detailed breakdown of the datasets, cell lineages, maturation stages, and methods applied in this study. We ran the experiments using the default parameters of each method, with a fixed root cell per trajectory, and 20 random seeds to account for potential stochasticity. For each evaluation metric, method, and trajectory, we averaged the scores across the 20 random seeds to calculate the mean score of each method for a specific trajectory. We found the variance across seeds to be negligible for all methods, except for sc-Shaper, which was more susceptible to seed selection for a few datasets, potentially due to its ensemble k-means clustering approach. Finally, we conducted one-sided paired t-tests with a significance level of 0.05 to test whether TimeFlow had a greater mean score than another method for each evaluation metric.

Supplementary Tables S6, S7, S8 provide the analytic scores per method and dataset for the Kendall’s Tau, Spearman’ s Rank correlation, Cell Accuracy and CSC, as well as the mean scores across all cell lineages, both with and without aggregation at the dataset level. The first row of Figure 3 presents the scores for each method applied to both inhouse and public datasets with branching trajectories. TimeFlow yielded the highest mean scores for the Kendall’s Tau, Spearman’s Rank correlations, and CSC. It followed closely Palantir and PAGA in Cell Accuracy, and ElPiGraph in PCC, and consistently performed statistically significantly better than TinGa for all metrics. Furthermore, the distributions of its scores were less variable compared to other methods and did not exhibit low-score outliers. PAGA generally produced highly accurate mean scores. However, it performed poorly for the neutrophilic and B-cell lineages in P3-BM, failing to capture the correct order of stage transitions (Supplementary Table S6). Regarding the PCC scores, we observed that ElPiGraph had the highest mean PCC score, despite its lower ranking in terms of CSC. This may be explained by its strong performance in the BM neutrophilic datasets, for which it yielded the highest PCC scores for CD16 among all methods. Palantir resulted in accurate orderings for most of the trajectories, except for P1-BM Neu, P2-BM Ery, Levine-13 Monocytes and Levine-32 B-cells, where it missed the expected order of stage transitions (Supplementary Table S5). As shown in the second row of Figure 3, TimeFlow consistently produced meaningful orderings for all small datasets, had the highest mean scores for all metrics and unlike other methods, did not suffer from large variability or outliers. Results for large datasets with linear trajectories in the third row of Figure 3 indicate that TimeFlow, ElPiLinear and DPT yielded highly accurate results, although DPT scored lower for PCC in neutrophilic specific markers, presumably due to coarse-grained pseudotime estimates and multiple duplicate assignments.

**Figure 3:**
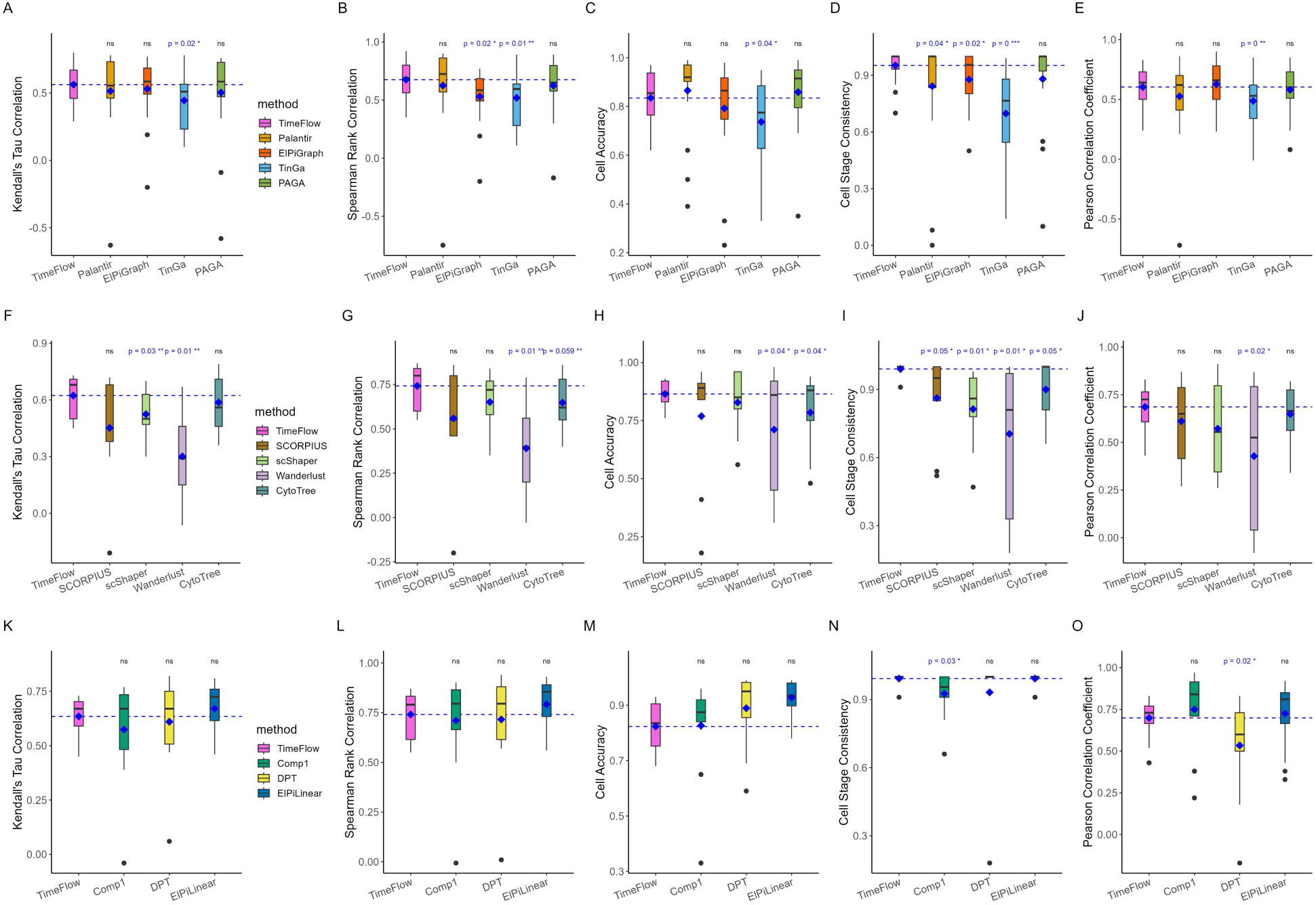
Methods comparisons across different datasets and results for the evaluation metrics of Kendall’s Tau and Spearman’s Rank correlation, Cell Accuracy, Cell Stage Consistency and Pearson’s correlation coefficient between pseudotime and lineage-specific CD markers. Boxplots show the median, quartiles, minimum, and maximum values. The blue diamond shape in the box indicates the mean score of a method. The horizontal dashed blue line represents the mean of TimeFlow for comparison. Results in the first row correspond to all in-house and public datasets with branching trajectories (P1/2/3-BM, Levine-13, Levine-32, Wishbone). The results in the second row correspond to small datasets with linear trajectories (P1/2/3-Mono/Ery/B-cells), and the results in the third row correspond to both small and large datasets with linear trajectories (P1/2/3-Mono/Neu/Ery/B-cells). P-values correspond to paired one-sided t-tests with a significance level of 0.05 (*signifies p-value *<* 0.05, **: p-value *<* 0.01, ***: p-value *<* 0.001, ns: not significant). More details on the datasets and methods used in the evaluation are given in Supplementary Table S5 and analytic scores for the metrics in the Supplementary Tables S6, S7, S8.

### 3.3 Interpretation of cell orderings for different trajectories

We discuss the results for the branching trajectory P1-BM. All methods correctly positioned the B-cells along their maturation stages, except for TinGa, which assigned early pseudotime values to disproportionally many mature B-cells, making indistinguishable the stage transitions (Figure 4(1-10). We observed that ElPiGraph concentrated most intermediate B-cells within a narrow time window (yellow shaded region in Figure 4(2), making barely visible this transitional stage. However, as illustrated in Figure 4(11-20), both ElPiGraph and TinGa accurately arranged the neutrophilic populations, with the percentages of mature neutrophils increasing progressively throughout the process. PAGA also positioned neutrophils in agreement with their stages, but multiple cells received the same pseudotime value, causing the transition from intermediate I to II and mature stages to be less evident (Figure 4(13)). Palantir resulted in substantial overlap between the two intermediate stages of neutrophils (Figure 4(14,19)). Figure 4(21-30) shows that both Palantir and PAGA yielded accurate orderings for the erythroid lineage, while ElPiGraph and TinGa failed to distribute sufficiently more mature erythrocytes at later pseudotime values than intermediate II erythrocytes. Despite some overlap between intermediate I and II erythrocytes, TimeFlow ordered most cells from the two intermediate stages in accordance with their known maturation stage. Many more comparisons among the different methods for branching trajectories are provided in Supplementary Figures S6-S11 and discussed in Supplementary Section S6. We remark that TimeFlow also captured reliably the expected marker patterns across linear trajectories from both small and large datasets (Supplementary Figures S12-S23). More results in linear trajectories are discussed in Supplementary Section S7 and analytic scores are presented in Supplementary Tables S7-S8.

**Figure 4:**
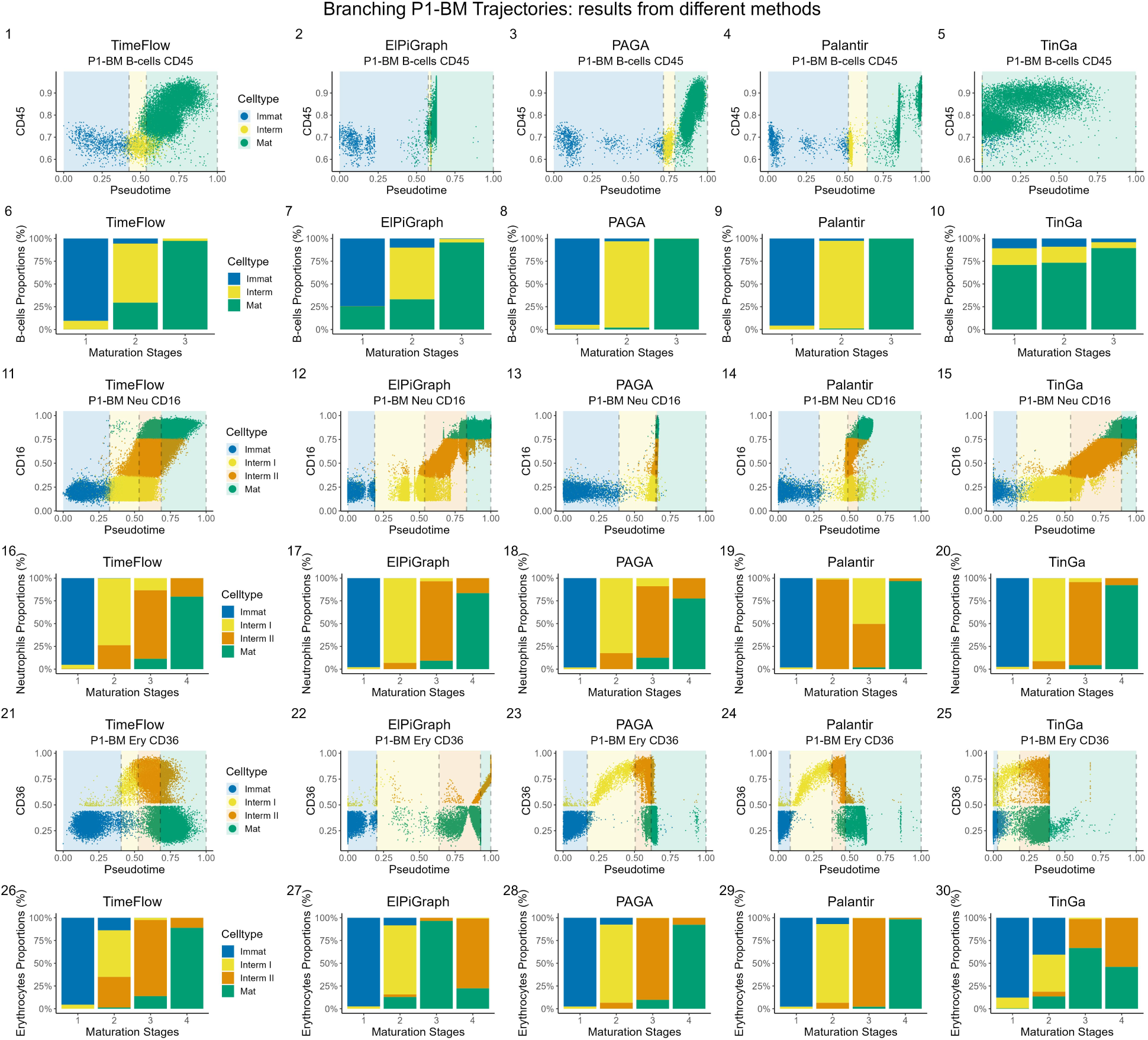
Side-by-side comparisons of TimeFlow, ElPiGraph, PAGA, Palantir, TinGa on the P1-BM dataset with a branching trajectory. Each scatterplot dot represents a cell, coloured by its maturation stage. The marker expression was scaled in [0,1]. The vertical dashed grey lines indicate the boundaries between the cell stages. For each cell stage, the cut-off is defined at the pseudotime value corresponding to the last cell in that stage’s ordered sequence (cumulative cell counts, see Section 2.3). Boundaries might overlap if a method does not separate between distinct stages. Regions correspond to different cell stages and are colored such that the transitions between them and their corresponding marker evolution are highlighted. Stacked bar plots show the cell distribution (%) in each of their known maturation stages, following pseudotime ordering. The stacked barplots cut-offs are the same with the cut-offs on the pseudotime axis. Gaps in the marker expression (y-axis) between consecutive cell populations are a consequence of using rectangular gates during dataset preparation. (1-10) Methods comparisons on the P1-BM B-cells dataset. (11-20) Methods comparisons on the P1-BM Neu dataset. (21-30) Methods comparisons on the P1-BM Ery dataset.

With regards to public branching datasets, all methods struggled to distinguish between the two early hematopoietic populations of Levine-13 B-cells, HSCs and MPPs (Supplementary Figure S24). TimeFlow pseudotime increased on average as cells progressed along the B-cell lineage (Supplementary Figure S24 A, F). Despite spreading immature B-cells along the entire pseudotime axis and assigning many mature CD38mid and low B-cells with early values, both PAGA and Palantir resulted in high accuracy and correlation scores (Supplementary Figure S24 C, D, H, I, Supplementary Table S5). Supplementary Figure S25 presents results on the Levine-13 monocytic lineage and interpretations are given in Supplementary Section S8. Regarding Levine-32 B-cells, PAGA outperformed the other methods and correctly assigned mature B-cells greater pseudotime values than Pre-B-cells (Supplementary Figure S26 C, H). It also distinguished better cell transitions between HSCs and HSPCs. PAGA was followed closely by TimeFlow and TinGa, for which the count of mature B-cells increased progressively along pseudotime. Palantir assigned several HSPCs high pseudotime values and incorrectly positioned Pre-B-cells at lower pseudotime values (Supplementary Figure S26 D, I). Supplementary Figure S27 presents results on Levine-32 monocytic lineage and interpretations are given in Supplementary Section S8. All methods respected the known cell stage ordering for the Wishbone dataset, where most lymphoid progenitors were correctly followed by the CD4+ SP and CD8+ SP populations (Supplementary Figure S28). However, several methods mapped the exact same pseudotime value to numerous cells, particularly near the beginning of this single-branched trajectory (Supplementary Section S8). Supplementary Figure S29 shows the runtime of TimeFlow and other methods that scaled for the largest datasets used in this study. The results are discussed in Supplementary Section S8. Overall, TimeFlow tended to preserve the known stage ordering for each branching trajectory and the marker patterns for each linear trajectory.

### 3.4 Entropy of pseudotime distributions for different methods

Pseudotime entropy serves as a measure of the cell spread and a proxy to evaluate the pseudotime resolution of the methods. Figure 5 presents a collection of pseudotime distributions and marker evolution plots for methods that scored 1 in terms of CSC and performed highly for all other metrics. Unlike TimeFlow, which spread the cells across the pseudotime axis, several other methods produced peaked, localized pseudotime distributions, with cells concentrated within a narrow range of pseudotime values. As verified by Figure 5 and Supplementary Table S9, which provides the entropy scores, TimeFlow consistently resulted in scores above 3.5, averaging 3.98 for linear trajectories and 3.93 for branching trajectories. Average scores of other methods for branching trajectories ranged from 2.47 to 2.7. The impact of coarse-grained pseudotime is evident in the marker evolution plots. For instance, in Figures 5A, F, TimeFlow, with an entropy score of 3.85, resolved in fine-scale the pseudotime of P1-BM Neu, showing the progression of the cells from one stage to another and the continuous changes in the CD16 expression. In Figure 5G, PAGA, which had an entropy of 1.27 for P1-BM Neu, indicated a sharp increase in CD16 around pseudotime 0.65 and assigned 47.55% of the cells to the range between 0.64 and 0.65, which corresponds to the boundaries of intermediate stages I and II. Assigning nearly half of the neutrophils the same pseudotime value limits the information available on stage transitions and fails to align with the expected dynamic changes in the CD16 expression during neutrophilic differentiation [70]. Similarly, in the P2-BM Neu trajectory, 64.48% of the cells were mapped to a narrow pseudotime range between 0.51 and 0.53 by PAGA, which had an entropy score of 1.17 (Figure 5D, I). For the same trajectory, Palantir did not scatter the cells across intermediate pseudotime values (Figure 5J) and yielded an entropy score of 1.52. ElPiGraph produced discrete groups of B-cells along pseudotime, making difficult to distinguish the transition from intermediate to mature B-cells (Figure 5K, R). For linear trajectories, DPT had an entropy score of 0.62 and accumulated nearly half of the P1-Neu cells (48.01%) within the pseudotime range of 0.991 and 0.994, where the cut-offs of the intermediate I and intermediate II stages were located (Figure 5N). Consequently, it did not capture the gradient CD16 expression pattern (Figure 5T). We assume that some of the above outcomes might be related to the inherent algorithmic steps of some methods such as graph node embeddings, clustering or community detection for cell aggregation, followed by projection of multiple individual cells onto these nodes/clusters. TimeFlow orders the cells without aggregating them into groups, but instead it reconstructs a density-driven path for each cell, avoiding low entropy pseudotime distributions. However, we stress that high entropy in pseudotime should not be interpreted as a measure of accurate cell ordering. For example, Wanderlust, which also computes the shortest path for each cell to the root, exhibited high entropy scores (Supplementary Table S9) but failed to respect the monocytic maturation stages in P1/2/3-Mono as verified in Supplementary Figure S12-14.

**Figure 5:**
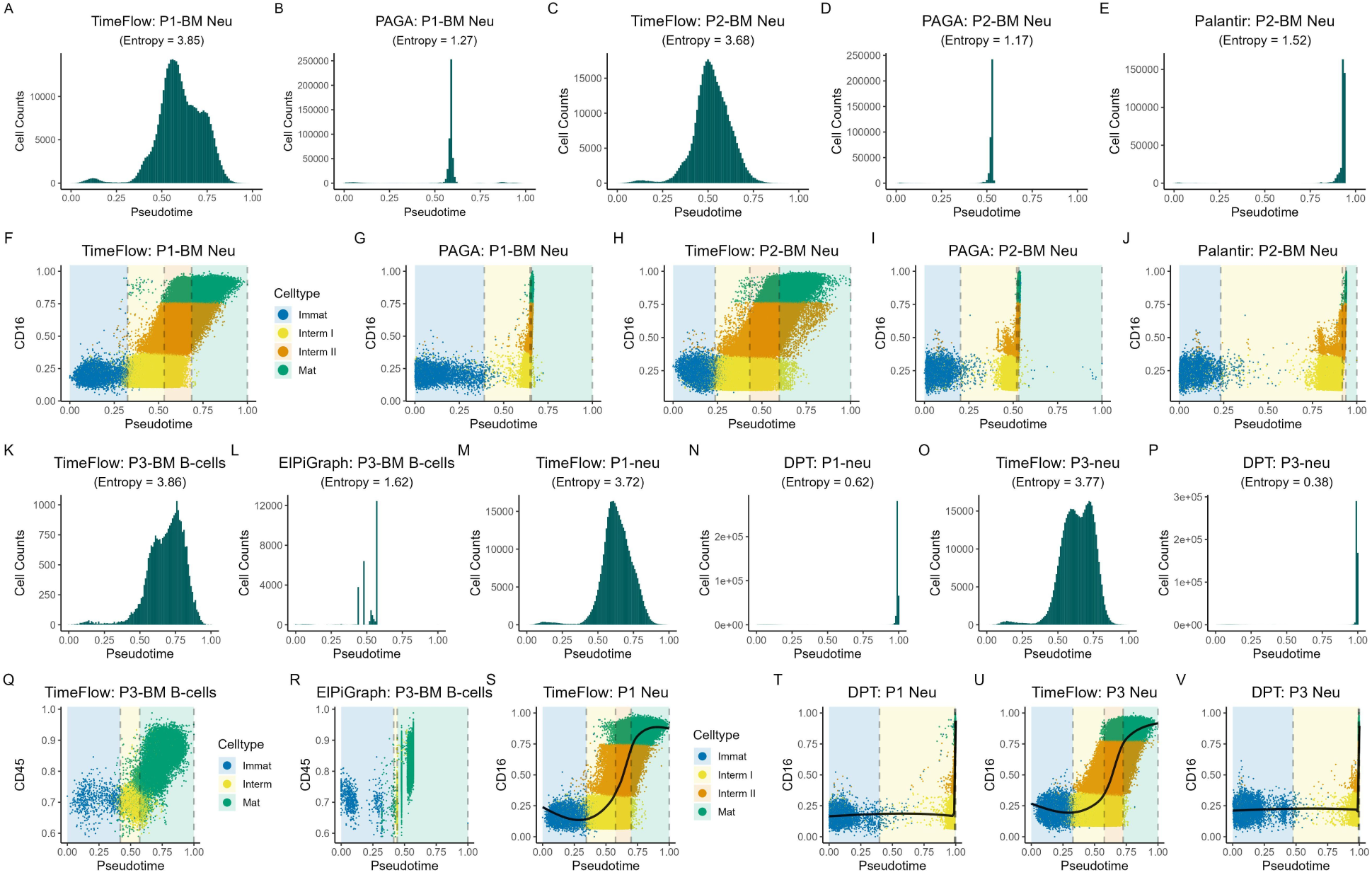
Side-by-side comparisons of pseudotime histograms along with their entropy scores and scatterplots between lineage-specific markers and pseudotime. Each scatterplot dot represents a cell, coloured by its maturation stage. The marker expression was scaled in [0,1]. The vertical dashed grey lines indicate the boundaries between the cell stages. For each cell stage, the cut-off is defined at the pseudotime value corresponding to the last cell in that stage’s ordered sequence (cumulative cell counts, see Section 2.3). Boundaries might overlap if a method does not separate between distinct stages. Regions correspond to different cell stages and are colored such that the transitions between them are highlighted. (A-J) Comparisons for P1/2-BM Neu between TimeFlow, PAGA and Palantir (K-V) Comparisons for P3-BM B-cells between TimeFlow and ElPiGraph and and P1/3-Neu between TimeFlow and DPT.

In summary, 1) TimeFlow orderings were in accordance with known cell stage transitions in branching trajectories and reflected the dynamics of well-studied markers in linear trajectories. The cell transitions were finely resolved, as indicated by its high pseudotime entropy scores 2) several methods originally designed for RNA-seq data adapted well to flow cytometry data, achieving high scores in terms of cell ordering accuracy. However, they often concentrated multiple cells within narrow pseudotime ranges without resolving the cell transitions, as implied by their low pseudotime entropy scores 3) computing the pseudotime with more than one method is essential to obtain more reliable results and compare the pseudotime distributions.

### 3.5 Meaningful pseudotime computation with TimeFlow across patients and unseen cell stages

We overlaid the expression patterns of several lineage-specific markers for the linear trajectories of P1/2/3. We used linear interpolation to align the marker expression along a common pseudotime axis of 1000 equally spaced points. This approach allowed us to account for the different dataset sizes and make comparable the fitted GAM cubic splines across the patients. Figure 6A-H shows clear agreement between the overlaid curves of the markers. Despite some expected variability between the three patients, TimeFlow resulted in consistent expression patterns, demonstrating its potential in describing a healthy differentiation process.

**Figure 6:**
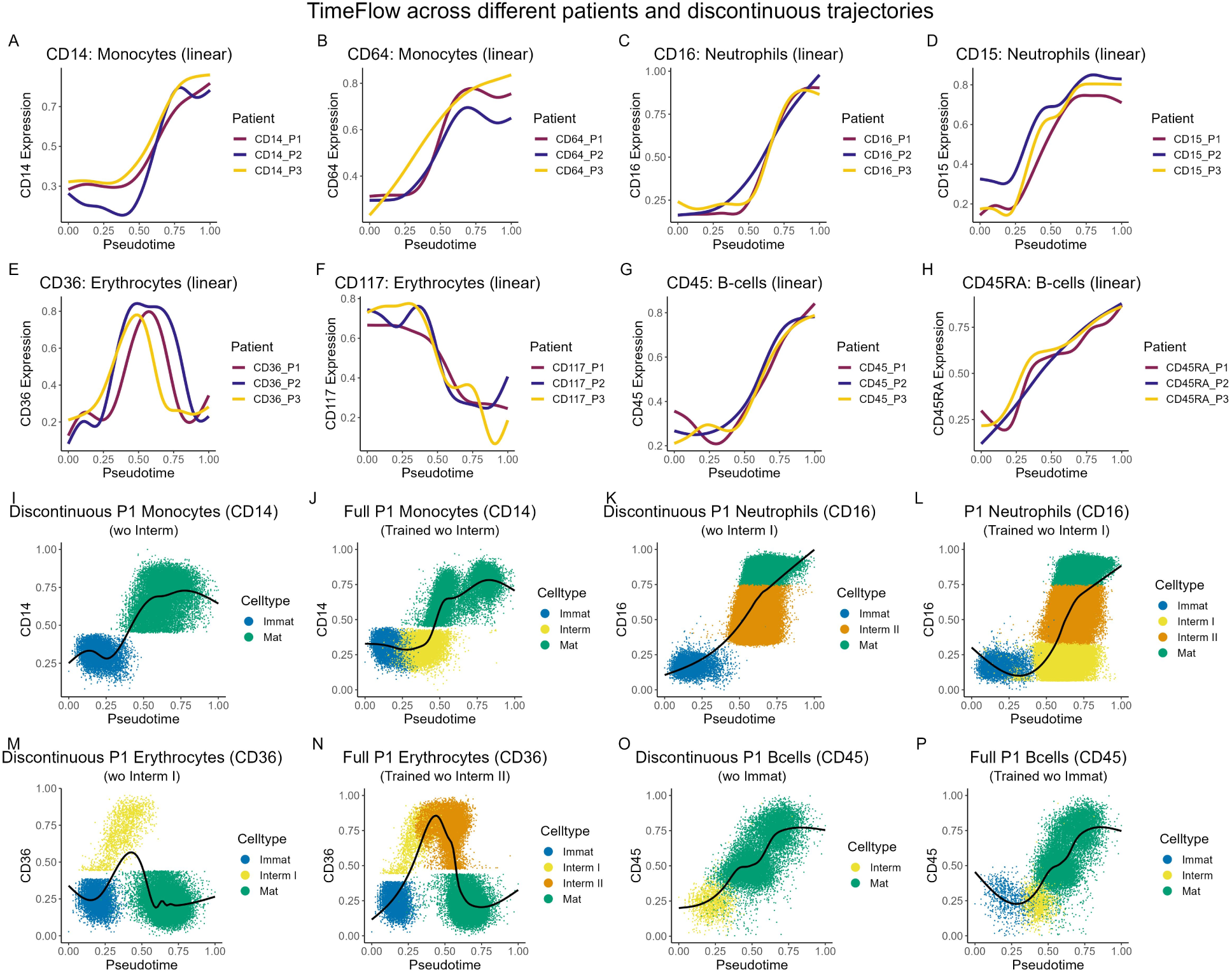
TimeFlow across linear trajectories from different patients and linear discontinuous trajectories. The marker expression was scaled in [0,1]. GAM models (solid black curves) were fitted without use of labels. Each scatterplot dot represents a cell, coloured by its maturation stage. (A-H) Overlaid marker dynamics of characteristic markers for P1/2/3-Mono/Neu/Ery/B-cells. (I-P) Marker dynamics across discontinuous and full linear trajectories from P1. Each scatterplot dot represents a cell, coloured by its maturation stage. The density module was fitted on the cells of each discontinuous trajectory and was used to estimate the pseudotime of the full trajectory, including the missing cell populations.

### 3.6 Accurate pseudotemporal orderings with TimeFlow by transferring weights across patients

Our method uses a generative model (Real NVP) for density fitting and learns the weights of its neural networks during the training process. We examined whether TimeFlow could reuse the weights learnt from one sample not only for pseudotime estimation of that specific sample, but also across independent samples that describe the same biological process, saving time and computation. We first fitted the density model to the branching P1-BM dataset and then used the inferred weights to estimate the cell pseudotime in the P2- and P3-BM datasets. We repeated this process, changing the patient for which the density model was fitted each time (e.g., transferring P2-BM weights to P1-BM and P3-BM). Supplementary Table S10 presents the CSC scores obtained from different experimental setups for each lineage in the BM datasets. TimeFlow resulted in the expected cell stage orderings for all lineages within the P1-BM dataset, achieving CSC scores from 0.91 to 1 when weights of P2- and P3-BM were used. Stage transitions also remained accurate when weights of P1-BM and P3-BM were transferred to P2-BM. All CSC scores were equal to 1, when P1-BM and P2-BM weights were applied to P3-BM, except for the P3-BM Ery (CSC=0.66) when P1-BM weights were used. We followed the same approach for linear trajectories. Despite swapping model weights across patients, the mean CSC scores remained above 0.95 for linear trajectories, too (Supplementary Table S11). The above results confirmed the generalization potential of TimeFlow across samples that describe the dynamics of the same differentiation process and the possibility to extend the density structure of the snapshot across patients.

### 3.7 Accurate pseudotemporal orderings with TimeFlow for unseen cell states

We challenged TimeFlow with pseudotime estimation for cell stages that are absent during the density fitting. We modified the linear trajectories, discarding cells from a particular cell stage, and thus created datasets that contained discontinuous trajectories. We fitted the density module to the remaining cells of each trajectory and used the learnt weights to evaluate the probability density function at each cell in the original corresponding datasets. Then, we re-estimated the pseudotime in each trajectory using the whole available data. Supplementary Tables S12-S15 present the CSC scores for the orderings of each population in the full, continuous linear trajectories, specifying which cell stage was absent during density fitting. We observed that TimeFlow preserved the expected cell stage orderings for the monocytes, neutrophils and erythrocytes of all three patients, independently of which cell stage was absent during density fitting. We only found lower scores (CSC=0.81) for three out of nine cases across the B-cell populations. For instance, as shown in Figure 6I-P, our method arranged accurately the new cells within the discontinuous trajectories without violating the stage ordering. The potential of TimeFlow to successfully integrate previously unseen cells in existing discontinuous trajectories, without refitting the density model, is a promising property for analysing pathological hematopoiesis in bone marrow, where discontinuous trajectories with gaps have been observed [64].

### 3.8 Sensitivity to density estimation method and robustness to other parameters

We examined whether TimeFlow is sensitive to the choice of different density estimation methods. We experimented with two different normalizing flow models, the Masked Affine Autoregressive Transform (MAF) [71] and the Masked Piecewise Rational Quadratic Autoregressive Transform (AR-RQS) [72], as well as Mellon [43], a new probabilistic density estimation method for single cell data (e.g., RNA, ATAC modalities). MAF transforms linearly the input data, while AR-RQS applies a non-linear rational quadratic spline transform. Unlike Real NVP, the transformation of each input dimension depends only on the preceding dimensions and is parametrized by an autoregressive neural network. Mellon computes a continuous, differentiable probability density function using a Gaussian process and linking the cell density estimation to the nearest-neighbour distance distributions. All the above methods allow for exact density evaluation which is required in TimeFlow step 1 and is easily interchangeable due to the modular nature of TimeFlow. We obtained new density estimates at single cell level and re-computed the cell pseudotime for both linear and branching trajectories ^1^. As illustrated in Figure 7A-D, F-H, each density-variant of TimeFlow ordered successively the cells from immature to intermediate and mature, capturing the gradual gain in the expression of CD14 and CD45 along the linear monocytic and B-cell trajectories, respectively. Paired t-tests with a significance level of 0.05 did not show statistically significant differences between the density-variants (Figure 7J, K), suggesting that what matters for our method are not the density estimates per se, but rather the density gradient, which is used to trace the cell evolution paths. We also conducted paired t-tests to compare the results of linear and branching trajectories using Real-NVP, MAF and AR-RQS for the evaluation metrics of Kendall’s Tau, Spearman’s Rank correlation, and the Cell Accuracy, but found no significant difference between these methods across the different trajectories.

**Figure 7:**
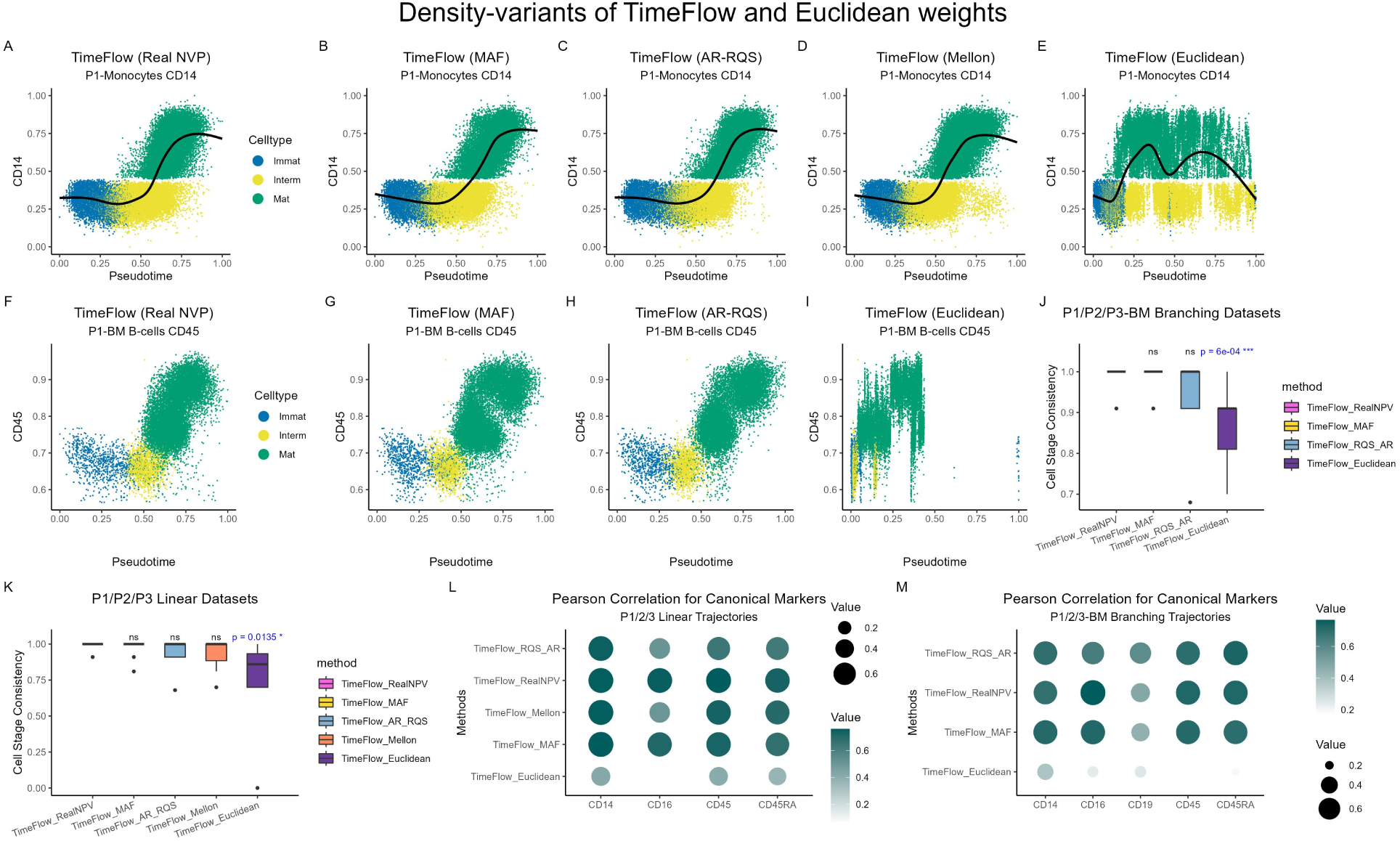
TimeFlow with three different density modules or Euclidean weights. Each scatterplot dot represents a cell, coloured by its maturation stage. The marker expression was scaled in [0,1]. For linear trajectories, a GAM model (solid black curve) was fitted without use of labels. Gaps in the marker expression (y-axis) between consecutive cell populations are a consequence of using rectangular gates during dataset preparation. (A-E) Marker expression of CD14 in P1-Mono along pseudotime using different density estimation methods for the density-gradient weights and the investigation of Euclidean weights. (F-I) Marker expression of CD45 in P1-BM B-cells along pseudotime. (J-K) Boxplots with Cell Stage Consistency scores for the TimeFlow density-variants and the Euclidean weighting across all datasets with branching or linear trajectories. They show the median, quartiles, minimum, and maximum values. P-values correspond to paired one-sided t-tests with a significance level of 0.05 (*signifies p-value *<* 0.05, **: p-value *<* 0.01, ***: p-value *<* 0.001, ns: not significant). (L-M) Average PCC scores between lineage specific markers and pseudotime inferred by TimeFlow density-variants or using Euclidean weights. Higher correlation scores are indicated by larger dot size and darker colour.

Although this is not backed by the criteria that TimeFlow satisfies (Section 2.1), we investigated replacing the density-gradient weights of the edges with the absolute Euclidean distance of their endpoints. This weighting resulted in obscure transitions between intermediate and mature cells and loss of the CD14 increase in the linear monocytic trajectory (Figure 7E). Furthermore, it distorted the orderings in the P1-BM B-cells trajectory (Figure 7I). Paired t-tests showed that TimeFlow-Real NVP performed significantly better than its Euclidean counterpart on both linear and branching trajectories in terms of CSC (Figure 7J, K). In terms of PCC, high correlations between lineage-specific markers and pseudotime are preserved for each density-variant but barely captured when Euclidean weights are used (Figure 7L, M). The negative impact of the Euclidean weights on the pseudotemporal orderings we observed aligns with previous studies, which have extensively discussed the problem of short-circuits on the graph [21, 22, 10, 65]. Computing shortest paths on a graph constructed in high-dimensional space and weighted by Euclidean distances increases the risk of spurious edges between cells that are distant in development but close in the ambient space (curse of dimensionality problem). Wanderlust addressed this problem using an ensemble of k-NN graphs, while DPT computed Euclidean distances in the diffusion-mapped space. Here, we showed empirically that the density-gradient weights protect TimeFlow from this short-circuits problem.

Finally, we examined the robustness of TimeFlow to different configurations of its hyper-parameters for the density and pseudotime modules using all the in-house datasets of the three patients. We tested different numbers of coupling layers, hidden neurons and batch sizes in the Real NVP architecture for branching and linear trajectories (Supplementary Tables S16-S17) and varied the number of k neighbours-the only parameter of the pseudotime module-from 3 to 20. We summarized the results of each Real NVP configuration in Supplementary Tables S18-S19 by averaging the scores over all patients. TimeFlow was robust to the Real NVP hyperparameters, and the mean CSC scores remained above 0.95 for both linear and branching trajectories. Following the same strategy, we summarized results for values of k in Supplementary Tables S20-S21. We consistently obtained CSC scores above 0.97 for monocytes, neutrophils and erythrocytes. B-Cell trajectories started deteriorating for values of k greater than 15 (CSC=0.88). We also analysed the distributions of scores obtained for all trajectories at each specific value of k with regards to Kendall’s Tau and Spearman’s Rank correlation, and Cell Accuracy. For each evaluation metric, we conducted paired t-tests to compare the score distributions for every pair of k values, resulting in ^18^ = 153 pairwise comparisons. We applied Bonferroni correction to control for false positives and obtained no p-values above the corrected threshold, indicating that our method is robust to the number of k.

## 4 Discussion

We developed TimeFlow to compute cell pseudotime in flow cytometry datasets with linear or branching trajectories. Our method is grounded in a model for cell evolution based on OT. In practice, it limits the cell evolution to its k-NN neighbourhood in the marker space and minimizes changes in the density space. TimeFlow finely and smoothly ordered monocytes, neutrophils, erythrocytes and B-cells within their specific differentiation pathway, and proved useful in modelling the marker dynamics along all linear trajectories. It showed flexibility in the choice of density estimation method and potential in generalizing across patients and previously unseen cell states without requiring re-training the density model.

### 4.1 Extensions for TimeFlow

Further algorithmic improvements can extend the use of TimeFlow. First, our method does not estimate the number of different cell lineages and does not assign cells with a lineage label. However, by construction, it can explore the structure of the snapshot and serve as the basis for a pseudotime-driven lineage detection strategy, which is necessary to fully understand branching trajectories and describe the markers’ dynamics within each lineage. In the future, the pseudotemporal orderings might guide the search of fully differentiated cell states, as well as lineage pathways. Second, TimeFlow expects a user-defined root cell, based on the expression of well-known early differentiation markers, as suggested in other works [21, 22]. Despite the careful selection of CD markers in cytometry experiments, if none of the included markers is representative of early differentiation, defining the root of the trajectory becomes complicated. An unsupervised strategy to identify stem cell populations would address the problem of defining the root of the differentiation process and enable TimeFlow to infer the dynamics of less characterized biological processes [66]. Third, by construction, TimeFlow is not suitable for disconnected trajectories, which evolve from different stem populations [15, 67]. To account for such trajectories, TimeFlow should adapt to disconnected graphs. The above algorithmic extensions would make TimeFlow fully independent from external user guidance in dissecting differentiation pathways.

### 4.2 Future directions with TimeFlow

Density estimation with generative modelling allowed TimeFlow to generalize pseudotime estimation across different samples from the same biological process by reusing the model parameters that were learnt from a specific patient, and without training the model from scratch. Additionally, TimeFlow integrated new cell populations under differentiation into existing trajectories, without violating the expected cell stage transitions. Our findings raise the question on the use of TimeFlow for further downstream tasks with biological interest. An example is the construction of a reference model for normal hematopoiesis by integration of bone marrow samples from different healthy patients. This direction aligns with recent advances in immunology, which focus on cross-dataset integration and construction of reference maps or single-cell atlases [68, 69, 70] to study immunological changes between normal and disrupted processes in the bone marrow. More generally, we expect TimeFlow to fit well in the flow cytometry community. Pseudotime analysis with TimeFlow offers hematologists and immunologists a different perspective compared to traditional manual gating when analysing BM subsets from flow cytometry. It is impractical to visually inspect the cells’ positions in ^D^ pairs of 2D marker scatterplots during manual gating. This practice also increases the risk of missing patterns that explain heterogeneity between differentiating cell populations, which have been previously unexplored. In contrast, TimeFlow accounts simultaneously for all the input dimensions, bringing a complementary data-driven approach to the study of cell differentiation dynamics in large datasets. It models consistently the marker dynamics across independent BM samples and supports the inference of reference patterns in healthy haematopoiesis. Another future direction of clinical interest is to compare these healthy patterns with the differentiation patterns from patients with hematological diseases such as myelodysplastic syndromes (MDS) and acute myeloid leukemia (AML). Comparing the cell abundance (cell proportions) along the pseudotime axis in healthy and pathological bone marrow samples might also support research in diagnosis of MDS or AML with flow cytometry. These comparisons will provide more evidence on the role of protein markers during dysregulated hematopoiesis and improve differential expression analysis.

## 5 Code Availability

TimeFlow is implemented with Python and PyTorch [71]. The code is available at the GitHub account https://github.com/MargaritaLiarou1/TimeFlow under the Creative Commons Attribution 4.0 International Public License. Interested users will also find tutorials to use TimeFlow and reproduce the results of this work.

## 6 Data Availability

Our new pre-processed hematopoietic flow cytometry datasets will be accessible upon acceptance. The public mass cytometry dataset for T cell development in mouse thymus is available at the GitHub page of Wishbone [24] https://github.com/ManuSetty/wishbone/ tree/master/data. The public mass cytometry datasets Levine-13 [65] and Levine-32 [66] are available for download from the R package HD-CytoData [64].

## 8 Institutional Review Board Statement

The study was conducted in accordance with the Declaration of Helsinki and approved by the Institutional Ethics Committee of the University Hospital Geneva (Ethics Committee number 2020-00174 (date: 3 March 2020) and 2020-00176 (date: 28 April 2020)).

## 9 Informed Consent Statement

Informed consent was obtained from all subjects involved in the study.

## 11 Acknowledgements

We thank the team of biologists from the Flow Diagnostics laboratory of the Clinical Pathology Service, particularly Julie Ducreux, as well as the technicians, in particular Cindy Lenvers, for their technical assistance and support.

## Footnotes

1 We tested all linear and branching P1/2/3 trajectories with the TimeFlow-AQR/MAF variants and only linear P1/2/3 trajectories with the TimeFlow-Mellon variant.

