## Supplementary Material for "TimeFlow: a density-driven pseudotime method for flow cytometry data analysis"

#### Contents

**1 Section S1 Connection to Optimal Transport** **3**

**2 Section S2 Background on the Real NVP transform** **4**

**3 Section S3 Technical implementation of TimeFlow** **6**

**4 Section S4 Datasets** **7**

**5 Section S5 Data pre-processing** **8**

**6 Section S6 Trajectory Inference methods used for comparisons** **8**

**7 Section S7 More results on branching and linear trajectories from in-house datasets 11**

**8 Section S8 More results on branching trajectories from public datasets 14**

**9 Section S9 Supplementary Tables 15**

### 1 Section S1 Connection to Optimal Transport

In this section, we detail how Optimal Transport (OT) theory [1, 2, 3, 4] supports the two criteria (a, b) related to changes in cell expression and cell density, which are presented in Section 2.1 of the main text and serve as the basis for TimeFlow.

While some Trajectory Inference (TI) methods study in vivo developmental processes using a single static snapshot taken at a specific time point, other methods target in vitro processes or time-course experiments, where multiple, independent cell snapshots are taken at a few different physical time points ($t_0 = 0 < \ldots < t_i < \ldots < t_K = T$). Methods falling in the latter category, usually model the developmental process as a time-varying probability distribution (i.e., stochastic process) to account for the available temporal labels [5, 6, 7, 8]. We borrow the notation of the main text, where a cell state is referred as $x_i \in X \subseteq R^D$, and X is the D-dimensional marker space. In addition, we denote the distribution of cells receiving a pseudotime value t by $f_t : X \to R^+, t \in [0,\ T]$. A reasonable question is how the stem cell distribution $\mu = f_0$ at $t = 0$ evolves into a mature cell distribution $\nu = f_T$ at $t = T$. OT-based methods for TI have addressed this question by computing transport plans or probabilistic mappings between cells at consecutive time points that move the source distribution to the target distribution (static OT) or by learning a time-continuous model that connects the marginal cell distributions over the time course (dynamic OT) [9]. A transport plan $\gamma$ moves the probability mass between two distributions $\mu$ and $\nu$ with some arbitrarily chosen cost. Assuming the Euclidean setting prevails in the markers' space, the cost function of moving a unit mass from the source $x \in X$ to target $y \in X$ is usually set to be the squared Euclidean distance $d(x, y) = ||x - y||^2$. Based on the Kantorovich formulation of OT [1], for any two probability measures $\mu$, $\nu$ on X, the Kantorovich or Wasserstein distance of order p between $\mu$ and $\nu$, is defined by the formula

$$W_p(\mu, \nu) := \inf_{\gamma \in \Pi(\mu,\nu)} \left[ \int_{X \times X} d(x, y)^p \ d\gamma(x, y) \right]^{\frac{1}{p}} \tag{1}$$

where $p \in [1, \infty)$ and $\Pi(\mu, \nu)$ the set of all couplings $\gamma$ on $X \times X$ with marginals are $\mu, \nu$ [1]. For consistency with the Euclidean setup, we consider $p = 1$. The optimal transport plan $\gamma^* \in \Pi(f_0, f_T)$ is the plan which achieves the minimum of the equation 1. By definition [1, 3, 10] for the given distributions $f_0$ and $f_T$, the constant speed geodesic (here shortest path) is given by the pushforward measure $f_t := (\pi_t)_{\#} \gamma^*$, where $\pi_t(\mu, \nu) = \left(1 - \frac{t}{T}\right) \mu + \frac{t}{T}\nu$ and $\gamma^*$ the optimal transport plan with $f_0$ and $f_T$ marginals. In this context, the geodesic term refers to a curve in the Wasserstein space of the probability measures that connects distributions $f_0$ and $f_T$ in the most cost-efficient way. Intuitively, this means that the temporal sequence of distributions $f_t(t \in [0, T])$ over X smoothly deforms (evolves) $f_0$ onto $f_T$, while minimizing the cost function d. Thus, for two cells close in differentiation with representations $x_i$ and $x_j$ at t and $t + \delta t$, respectively, as mentioned in the main text, we expect:

a. $d(x_i, x_j) \leq \epsilon_1$, for a small value of $\epsilon_1$: the representations of the cell at t and $t + \delta t$ to have similar marker expression,

b. $|f_{t+\delta t}(x_j) - f_t(x_i)| \leq \epsilon_2$, for a small value of $\epsilon_2$: the pointwise changes between the marginal temporal distributions of the cells to be smooth and translated by the markers' evolution.

Unlike other OT-based TI methods, which can access both $f_0$ and $f_T$, in our single static snapshot setup, there are no temporal labels that would allow explicit estimation of transport maps. Instead, we only assume access to the stem cell distribution $f_0$ based on some prior biological knowledge (e.g., $CD34^{++}$ haematopoietic stem and progenitor cells (HSPCs)). The implementation of TimeFlow, presented in the main text Section 2.2, is a greedy solution that reconstructs cell paths by adapting the above two criteria in the single static snapshot setup. This snapshot is a sample of the t-marginal distribution $f = \sum_{t=0}^{T} f_t$ into the marker space X. Assuming a localized mass split for $f_t$, the density at $x_i$ is dominated by the value of $f_{t_i}(x_i)$. We characterize the geodesic of distribution f as many pointwise cell geodesics under the form of shortest paths in the snapshot, computed by the Dijkstra [11] algorithm. We limit the evolution of a cell to its k-Nearest Neighbours (k-NN) in the marker space (criterion a) and compute the shortest paths by minimizing the changes in the density space (criterion b). Dijkstra is by nature a greedy algorithm that provides a locally optimal solution to the shortest path problem. In the context of TimeFlow, the shortest paths show the most likely and least cost cell evolution routes in the snapshot. This is a greedy solution on a discrete graph structure that approximates the smooth variations of the cell density in the continuous space and agrees with the OT approach, where the objective is to minimize the total cost of transporting cell mass between consecutive experimental time points.

### 2 Section S2 Background on the Real NVP transform

TimeFlow uses the Real Non-Volume Preserving transformation (Real NVP) [12] to model the probability density function of the cells. Here, we follow the notation of Section 2.2 in the main text and detail the Real NVP transformation. Transforming a multivariate Gaussian base distribution with an affine transformation $x = g(z) = Az + b$, where A an invertible square matrix and $z = g^{-1}(x) = A^{-1}(x - b)$ the inverse transformation, results in another Gaussian target distribution. Real NVP exploits the invertibility of an affine transformation, while also allowing for non-Gaussian target distributions by stacking multiple affine coupling layers. Each of these layers transforms a subset of the input dimensions with a scaling and a translation operation, whose parameters are learnt by neural networks. Real NVP splits the vector z into two disjoint parts $z = [z_{1:d}, z_{d+1:D}]$ of dimensionality d and D − d, respectively, and then maps z into a vector x, partitioned as $x = [x_{1:d}, x_{d+1:D}]$. The first part of the output vector, $x_{1:d}$, remains unchanged, while the second part $x_{d+1:D}$ receives an affine transformation whose coefficients are given by nonlinear functions of $z_{1:d}$. The mapping is given as

$$\begin{aligned} x_{1:d} &= z_{1:d} \\ x_{d+1:D} &= \exp\left(s\left(z_{1:d}, \theta_s\right)\right) \odot z_{d+1:D} + t\left(z_{1:d,} \theta_t\right), \end{aligned} \tag{2}$$

where $\odot$ the elementwise Hadamard product, $s(\cdot)$ and $t(\cdot) : R^d \to R^{D-d}$, the scaling and translation operations of the transformation. These operations are implemented by neural networks with learnable parameters $\theta = [\theta_s, \theta_t]$. The inverse mapping is computed as

$$\begin{aligned} z_{1:d} &= x_{1:d} \\ z_{d+1:D} &= \exp\left(-s\left(x_{1:d}, \theta_s\right)\right) \odot \left(x_{d+1:D} - t\left(x_{1:d,}, \theta_t\right)\right). \end{aligned} \tag{3}$$

If the observed data are independent and identically distributed, then the training objective of flow-based models is the average negative log likelihood function, given as

$$\begin{aligned} &\operatorname{argmin}_\theta E_x\left[-\log f_x(x)\right] = \frac{1}{N}\sum_{n=1}^{N} \log f_x\left(x_n \mid \theta\right) \\ &= \frac{1}{N}\sum_{n=1}^{N}\left\{\log f_Z\left(\, g_\theta^{-1}\left(x_n\right)\right) + \log\left|\det\left(\frac{\partial g_\theta^{-1}\left(x_n\right)}{\partial x^T}\right)\right|\right\}. \end{aligned} \tag{4}$$

The density module in TimeFlow assumes a multivariate standard Gaussian distribution $N(z|0, I_D)$ for the Z variable and implements K Real NVP coupling layers, forming a deep neural network architecture that ensures an expressive transformation. The training objective function in 4 becomes

$$\frac{1}{N}\sum_{n=1}^{N}\left\{\frac{D}{2}\log(2\pi) + \frac{1}{2}\left\|\, g_k^{-1}\left(x_n\right)\right\|^2 + \sum_{k=1}^{K}\log\left|\det\left(\frac{\partial\, g_k^{-1}\left(x_n\right)}{\partial x^T}\right)\right|\right\}. \tag{5}$$

As mentioned in the main text, evaluating the density function at each data point $x_i$ requires the Jacobian matrix of the inverse transformation. For one Real NVP transform, this Jacobian matrix is computed as

$$J = \begin{bmatrix} I_d & 0 \\ \frac{\partial z_{d+1:D}}{\partial x_{1:d}} & \operatorname{diag}\left(\exp\left[-s\left(x_{1:d}\right)\right]\right) \end{bmatrix}. \tag{6}$$

This is a lower triangular matrix whose determinant simplifies to the product of the diagonal elements

$$\det \mathrm{J} = \prod_{\mathrm{j}=1}^{\mathrm{d}} \exp\left(-\mathrm{s}\left(\mathrm{x}_{1:\mathrm{d}}\right)\right)_{\mathrm{j}} = \exp\left[\sum_{\mathrm{j}=1}^{\mathrm{d}} -\mathrm{s}\left(\mathrm{x}_{1:\mathrm{d}}\right)_{\mathrm{j}}\right]. \tag{7}$$

For K coupling layers, the log of the Jacobian determinant is given by the sum of the log determinants for each layer. It is recommended to shuffle the order of dimensions after each coupling layer to ensure that each dimension is subject to an affine transformation. For further information on the theoretical background and the application of normalizing flows, we refer the reader to [13, 14, 15, 16].

### 3 Section S3 Technical implementation of TimeFlow

TimeFlow consists of two modules: the density estimation, written with PyTorch [17], and the pseudotime computation module, written with Python 3.11.3. In the density estimation module, we followed the educational paradigm from [18] to implement the Real NVP architecture and infer the log probability density function of the input data. The architecture included ten coupling layers implemented as feed-forward neural networks. For the shift operation, the networks used four fully connected hidden layers, followed by the Leaky ReLU function. For the scaling operation, the networks used three fully connected hidden layers, followed by Leaky ReLU and Tanh functions (this combination has been found to perform best [19]). The data were split into a 70% training set and a 30% validation set for density estimation on a specific dataset with early stopping. To compare different architectures, we recommend the user to include a test set. Our models were trained for a maximum of 500 epochs, and early stopping with a 15-epoch patience was applied if there was no improvement on the validation set after 15 consecutive epochs. The order of input dimensions was reversed after each coupling layer. All experiments were run using CPUs, but GPUs may also be used to accelerate the training. In the pseudotime module, we constructed a k-Nearest Neighbour Graph in the original space using the pairwise Euclidean distances between the cells. We initially set the value of k to five to preserve the local geometry information and increased it by one if there were disconnected components in the graph to avoid non-existing shortest paths (TimeFlow targets connected differentiation processes). We weighted the graph based on the absolute difference of the log probability density at the endpoints of each edge and computed the density-driven shortest paths using the Python version of the igraph package [20]. Following the example of other TI methods [21, 22], the root cell of each trajectory was manually chosen based on high CD34 expression, which is characteristic to hematopoietic stem and progenitor cells (HSPCs). Then, we accumulated the Euclidean distances between the nodes that comprise each path. The sum of a cell's path corresponds to its pseudotime and describes the evolution of its marker expression from the onset of differentiation. To model the evolution of the markers for a specific lineage, we selected the Generalized Additive Model (GAM) with a cubic spline from the mgcv R package [23] or PyGAM [24]. We examined the sensitivity of TimeFlow and found it robust the root selection (results not presented here). As a good practice, we recommend pseudotime estimation with multiple roots and pseudotime averaging for more reliable results.

### 4 Section S4 Datasets

#### 4.1 S4.1 In-house hematopoietic datasets

The panel of CD markers for the in-house flow cytometry datasets, includes the following 20 markers: CD200, CD45, CD45RA, CD64, CD3, CD15, CD133, CD117, CD56, HLA.DR, CD19, CD33, CD34, CD371, CD7, CD16, CD123, CD36, CD38. Supplementary Table S1 presents the cell counts for each population.

#### 4.2 S4.2 Levine-13 dataset

The Levine-13 mass cytometry dataset [25] measures the expression of 13 CD markers on 167,044 cells, out of which 81,747 have been manually gated for 24 immune cell populations. The remaining 85,297 cells are labelled as unassigned. The 13 surface markers are: CD45, CD45RA, CD19, CD11b, CD4, CD8, CD34, CD20, CD33, CD123, CD38, CD90, and CD3. Table S2 presents the cell counts for each population. To evaluate the Pearson Correlation between pseudotime and markers, we selected CD45, CD45RA, CD20 and CD19 for the B-cells lineage and CD11b for the monocytic lineage.

#### 4.3 S4.3 Levine-32 dataset

The Levine-32 mass cytometry dataset [25] measures 265,627 cells from the bone marrow of two healthy patients and consists of 32 CD markers: CD45RA, CD133, CD19, CD22, CD11b, CD4, CD8, CD34, Flt3, CD20, CXCR4, CD235ab, CD45, CD123, CD321, CD14, CD33, CD47, CD11c, CD7, CD15, CD16, CD44, CD38, CD13, CD3, CD61, CD117, CD49d, HLA-DR, CD64, CD41. In total 104,184 cells have been manually gated for 14 cell populations and 161,443 cells remain unassigned. Table S3 presents the cell counts for each population. To evaluate the Pearson Correlation between pseudotime and markers, we selected CD45, CD45RA, CD20 and CD19 for the B-cells lineage and CD11b for the monocytic lineage.

##### 4.4 Wishbone dataset

The Wishbone [22] mass cytometry dataset measures 25,000 cells for 13 surface-protein markers (CD27, CD4, CD5, CD127, CD44, CD69, CD117, CD62L, CD24, CD3, CD8, CD25 and TCRb). Table S4 shows the cell counts for each population. To evaluate the Pearson Correlation between pseudotime and markers, we selected CD8 and CD4 for the $CD8^+$ SP and $CD4^+$ SP lineages, respectively.

#### 5 Section S5 Data pre-processing

Our new in-house hematopoietic datasets were pre-processed with the R packages PeacoQC [26] and flowCore [27] for standard steps such as removal of marginal events, compensation, logicle transformation, removal of doublets and debris cells. The public mass cytometry datasets of Levine-13, Levine-32 and Wishbone were arcsinh-transformed with cofactor of 5. Clean data were also scaled to [0,1] with min-max scaling for numerical stability during density estimation with Real NVP.

#### 6 Section S6 Trajectory Inference methods used for comparisons

We compared the pseudotemporal orderings of TimeFlow to eleven other methods for pseudotime and trajectory inference. We provide a brief description of the selected TI methods.

1. **Wanderlust**

   Wanderlust [21] is designed for flow and mass cytometry datasets and models linear trajectories. It represents the cells with nodes on a k-nearest neighbour graph (k-NNG) in the high-dimensional space defined by the markers. To address short circuits—spurious edges between cells that are close in the marker's space but distant in maturation—Wanderlust introduces stochasticity. Specifically, it randomly discards a few connections from each node in the original graph to generate a set of sub-graphs. For each generated sub-graph, Wanderlust computes the shortest path distance from the root cell to every other cell by summing the edge weights (Euclidean distances). To further mitigate the problem of noisy paths, Wanderlust also introduces waypoints, which are randomly sampled cells. It computes the shortest path distance of each cell from all waypoints and calculates a weighted average of these distances. Waypoints closer to the cell are assigned with higher weights to influence more the overall distance. Finally, Wanderlust averages the shortest path distances obtained from each sub-graph to derive a robust pseudotime estimate for each cell.

2. **SCORPIUS**

   SCORPIUS [28] is designed for RNA-seq data and models only linear trajectories. It finds a low-dimensional representation of the data using multi-dimensional scaling and groups the cells into clusters using k-means. Then, it connects the cluster centroids with a minimum spanning tree (MST) that serves as the initial backbone for the linear trajectory. SCORPIUS further refines the trajectory by iteratively applying the principal curve algorithm. After each iteration, cells are orthogonally projected onto the trajectory, and a new curve is constructed by locally averaging the cells. The cell pseudotime is determined based on the cell's geodesic distance from the root cell.

3. **Component 1**

   Component1 [29] models only linear trajectories. It applies PCA on the input data and returns the first component as a linear trajectory.

4. **scShaper**

   scShaper [30] is designed for RNA-seq data and models only linear trajectories. scShaper proposes an ensemble approach that applies k-means clustering multiple times and combines the cell clusters into one consensus solution. To find a linear trajectory through the cluster centroids, scShaper solves the shortest Hamiltonian path problem. Finally, it assigns the cell clusters with discrete pseudotime values and the individual cells with a continuous estimate.

5. **Diffusion Pseudotime (DPT)**

   DPT [31] constructs a k-NN graph whose edge weights are given by Gaussian kernel convolutions centered at nearby cells. These weights correspond to cell transition probabilities and are stored in a transition matrix, whose eigenvectors are known as diffusion components, and may be used for dimensionality reduction. However, DPT uses the full-rank transition matrix. By raising this matrix to different powers, DPT computes random walks of any length between the root cell and any other cell. DPT finds branching points using a heuristic based on the Kendall's Tau correlation between DPT distances of cells. Branching points are detected when anti-correlated distances start showing increasing correlation.

6. **Partition-based Graph Abstraction (PAGA)**

   PAGA [32] targets RNA-seq data with complex trajectories and creates an abstracted graph representation of cell pathways based on a connectivity statistical hypothesis test.

PAGA initially constructs a k-NN graph and uses the Louvain community detection algorithm to find cell partitions (groups). These partitions represent the nodes of the abstracted graph and can be computed at different resolutions. To determine the connectivity between cells partitions, PAGA uses a statistical test based on the actual number of inter-edges between cell partitions on the k-NNG. These inter-edges are compared to the inter-edges expected under a random graph model. PAGA weighs the edges based on a confidence score, which serves as a threshold to discard edges when lower resolution for abstractness is desired. Pseudotime is computed at single cell level based on the random walk diffusion distance described earlier for DPT.

7. **Palantir**

   Palantir [33] is designed for RNA-seq data and models cell differentiation as a stochastic process on a graph. It finds branching trajectories and provides a solution for automated lineage detection. Initially, Palantir embeds the data into a low-dimensional space using diffusion maps and constructs an undirected k-NN graph. Then, it calculates cell pseudotime using the shortest path distances from the root cell, and adopts Wanderlust's waypoints concept to refine the cell positions. Furthermore, it discards several graph edges pointing from "later" to "earlier" cells and gives directionality to the graph. This allows Palantir to simulate random walks on the directed graph and compute a Markov chain transition matrix. Palantir uses this matrix to identify cells where random walks converge and marks them as terminal states (fully differentiated cells–extrema of the diffusion components). Finally, it removes all the outgoing edges from the terminal cells and computes the probability (differentiation potential) of each cell to reach every terminal state.

8. **ElPiGraph and ElPiLinear**

   ElpiGraph [34] is a principal graph algorithm for tree-shaped or disconnected trajectories from RNA-seq data. It begins with a random graph and modifies it using graph grammar rules such as edge bisection or node addition to generate a set of candidate graphs. ElPiGraph solves iteratively an optimization problem to converge to the optimal graph structure that represents the differentiation trajectory. The objective is to minimize the sum of the elastic principal graph energy and the mean squared distances between the data (cells) and the injected nodes. The elastic energy term consists of the total length of the edges and the graph harmonicity. This objective function allows ElPiGraph to select the graph structure that best balances the reconstruction error and the graph complexity. Once ElPiGraph converges to the final structure, cells are projected onto their closest nodes or edges. ElPiLinear is a variant of ElPiGraph tailored for linear trajectories.

9. **TinGa**

   TinGa [35] is designed for RNA-seq data with complex trajectories. TinGa reduces the data dimensionality and uses the Growing Neural Gas method to grow a graph and fit the data. The initial graph structure is iteratively updated to homogeneously cover the data (cells). After convergence to the final graph structure, TinGa fits an MST to remove triangles from the graph and refines the tree with further heuristics and post-processing steps.

10. **CytoTree**

    CytoTree [36] takes as input flow or mass cytometry datasets and infers linear or tree-shaped trajectories. CytoTree first runs a workflow that includes dimensionality reduction, cell clustering, and data downsampling. Then, it fits an MST on the cluster centroids to represent the trajectory backbone. Using the Louvain algorithm, it detects clusters into the MST branches and identifies differentially expressed markers in each branch. CytoTree further constructs a k-nearest neighbours (k-NN) graph to calculate the shortest path distances from the root cell to all other cells. These distances are converted into pseudotime estimates.

The well-known methods of Slingshot [37] and Monocle3 [38] for branching RNA-seq trajectories showed prohibitive time and memory costs on the large cytometry datasets and were not included in this study. To derive the pseudotemporal orderings of Section 3.4 in the main text for all the above TI methods, we run each method at its default parameters. We applied Wanderlust, ElPiLinear, Component1, SCORPIUS, scShaper, ElPiGraph and TinGa using the R packages dynmethods and dyno from the dynverse collection: https://dynverse.org/ [29]. Results for Palantir, CytoTree, DPT and PAGA were obtained based on the following tutorials or documentations: `https://github.com/dpeerlab/Palantir/tree/master/notebooks`, `https://ytdai.github.io/CytoTree`, `https://scanpy.readthedocs.io/en/stable/generated/scanpy.tl.dpt.html`, and `https://scanpy-tutorials.readthedocs.io/en/latest/paga-paul15.html`.

### 7 Section S7 More results on branching and linear trajectories from in-house datasets

We discuss the results of TimeFlow, ElPiGraph, PAGA, Palantir and TinGa on the P1/2/3-BM datasets, which are illustrated in Supplementary Figures S6-S11 and reported in Supplementary Table S6. As verified in Supplementary Table S6 TimeFlow, PAGA, ElPiGraph and Palantir achieved high mean scores for all evaluation metrics in the P1-BM monocytic trajectory, and as seen in Supplementary Figure S6 (first and second rows) these methods made evident the stage transitions along pseudotime. All methods except for Palantir resulted in correct stage ordering for P1-BM Neu (Supplementary Figure S6). We note that PAGA made barely visible the transition between the final three maturation stages due to multiple duplicate pseudotime assignments (Supplementary Figure S6). Regarding P1-BM Erythrocytes, Palantir, PAGA and TimeFlow spread the cells in line with their maturation stage, while ElPiGraph and TinGa failed to position most mature erythrocytes after intermediate II cells (Supplementary Figure S7). Similarly, as seen in Supplementary Figure S7 (third and fourth rows), ElPiGraph did not clearly resolve the transitions between intermediate and mature B-cells, and TinGa assigned disproportionally many mature cells with early pseudotime values. While all methods agreed in the stage ordering for P2-BM Mono (Supplementary Figure S8, Supplementary Table S6), we noticed that mature monocytes tended to be concentrated within a narrow range of pseudotime values for ElPiGraph, PAGA, Palantir and TinGa, as implied also by their low entropy scores in Supplementary Table S9. The same phenomenon was observed for the P2-BM Neu trajectory for PAGA and Palantir (Supplementary Figure S8). PAGA resulted in less errors between intermediate and mature erythrocytes in P2-BM Ery trajectory compared to the other methods. Palantir, TinGa and ElPiGraph clearly assigned more mature erythrocytes before intermediate II erythrocytes compared to TimeFlow (Supplementary Figure S9, first and second rows). ElPiGraph and TinGa were the only methods that did not distinguish between the intermediate and mature B-cell transitions (Supplementary Figure S9). All methods ordered monocytes in agreement with their maturation stages in the P3-BM Mono trajectory (Supplementary Figure S10, first and second rows). TinGa, ElPiGraph, TimeFlow and Palantir respected the neutrophilic maturation stages in P3-BM Neu (Supplementary Figure S10, third and fourth rows), but Palantir concentrated most mature neutrophils within a narrow pseudotime range, as also indicated by its low entropy score (Supplementary Table S9). PAGA assigned mature neutrophils before intermediate I and II neutrophils (Supplementary Figure S10). TimeFlow clearly distinguished between intermediate II and mature erythrocytes of the P3-BM Ery. PAGA assigned several mature erythrocytes with early pseudotime values but on average respected the known stage transitions, and Palantir distinguished accurately the stage transitions (Supplementary Figure S11, first and second rows). ElPiGraph and TinGa failed in positioning mature erythrocytes later than intermediate II cells (Supplementary Figure S11, first and second rows). PAGA wrongly shifted mature P3-BM B-cells ahead of intermediate B-cells, and TinGa positioned all mature B-cells in the beginning of the pseudotime axis (Supplementary Figure S11, third and fourth rows). TimeFlow, ElPiGraph and Palantir captured more accurately the B-cell progression along pseudotime.

Supplementary Figures S12-S23 illustrate the results of different methods on the linear trajectories of P1/2/3-Mono/Ery/B-cells/Neu datasets and Supplementary Tables S7-S8 provided the analytic scores. Supplementary Figures S12-S14 show the monocytic trajectories. TimeFlow captured the smooth increase in the CD14 expression for all three patients. Moreover, Comp1, DPT, ElPiLinear, and CytoTree had CSC=1 for each monocytic trajectory. Wanderlust struggled with positioning late-stage monocytes after mid-stage monocytes, and scShaper yielded variable scores for different random seeds. Supplementary Figures S15-S17 present the results on the P1/2/3-B-cells datasets. TimeFlow finely resolved the cell transitions and captured the increase in CD45 as cells progressed from immature to mature stages. Despite the overlap between immature and intermediate B-cells, Comp1 also captured the CD45 increase. DPT consistently yielded valid transitions but produced unexpected gaps in the trajectory. Wanderlust and SCORPIUS had a mean CSC score of 1 for the P1/2-B-cells datasets, but performed poorly for P2 B-cells. ElPiLinear produced accurate and highly resolved B-cell trajectories. CytoTree respected the stage transitions but estimated discrete pseudotime values. scShaper tended to produce accurate orderings (except for P3 B-cells) but was found prone to the random seed selection. Supplementary Figures S18-S20 show the results on the erythrocytic differentiation. TimeFlow found the expected CD36 non-linear pattern for each trajectory and was not affected by the overlap between intermediate II and mature erythrocytes of P2. The linear approach of Comp1 (PCA) was not sufficient to resolve the transitions between erythrocytes, particularly towards the end of the trajectory. DPT performed very well on all three erythrocytic trajectories. SCORPIUS, scShaper and CytoTree failed in capturing the characteristic CD36 pattern. ElPiLinear performed very well on P1/3-Ery, but suffered from substantial overlap on P2-Ery, missing the expected CD36 pattern. Supplementary Figures S21-S23 show results on P1/2/3-Neu for TimeFlow, Comp1, DPT and ElPiLinear. TimeFlow ordered the cells in agreement with their known stage and reflected the smooth changes in the CD16 expession curves. However, it only positioned slightly more mature than intermediate II cells near the end of the trajectory. Comp1 tended to merge early neutrophils, but accurately distributed intermediate II and mature neutrophils along pseudotime. ElPiLinear consistently had CSC=1 for linear neutrophilic trajectories. DPT respected the stage order for P1 and P3 but positioned late cells within a narrow range of pseudotime values, contradicting the continuum of transitions, as verified by its low entropy scores (Supplementary Table S9). It had low evaluation scores for P2-Neu (Supplementary Table S7-S8) because it dispersed mature and intermediate neutrophils towards the beginning of the pseudotime axis.

### 8 Section S8 More results on branching trajectories from public datasets

We discuss the results on the Levine-13, Levine-32 and Wishbone public mass cytometry datasets, which describe branching trajectories. In Levine-13 monocytic lineage, all methods resulted in substantial overlaps between cells from the early hematopoietic stages (HSC, MPP, CMP cells) (Supplementary Figure S25). As shown in the violin plots, the median pseudotime of the GMPs and monocytic populations increased progressively along the TimeFlow pseudotime. ElPiGraph distributed immature HSCs and MPPs widely across the entire pseudotime axis, and concentrated CMP cells at earlier pseudotime values. Furthermore, it positioned CD11b mid and high monocytes within narrow pseudotime ranges, leading to multiple cells with outlier pseudotime values. PAGA assigned CMP cells with earlier pseudotime values than HSCs or MPPs and spread along the entire pseudotime axis the GMP cells. It ordered more accurately the mature monocytic populations, distinguishing between their transitions. Palantir unexpectedly shifted mature monocytes from all three stages toward the beginning of the pseudotime axis and performed poorly across all evaluation metrics. TinGa accumulated most CD11b mid and high monocytes within narrow pseudotime ranges, suggesting coarse cell transitions. Supplementary Figure S27 presents a side-by-side comparison on the Levine-32 monocytic lineage. As shown in the violin plots in the first row, pseudotime generally increased on average as cells progressed along the monocytic lineage for all methods, except for ElPiGraph, where HSCPs tended to have lower pseudotime values than HSCs. PAGA clearly reflected the expected ordering. As shown in the stacked bar plots, Palantir placed mainly HSPCs before HSCs and did not clearly resolve the transitions between monocytes. TinGa wrongly assigned with early pseudotime multiple cells from the monocytic stage. Supplementary Figure S28 presents side-by-side the markers of CD4, CD8, CD3, and CD25 along the inferred pseudotime. These are well-studied markers for T-cell lymphopoiesis. All methods ordered most cells in accordance with their known stage and the expected marker patterns. However, we observed that ElPiGraph, PAGA, and Palantir produced discontinuous trajectories and tended to group many lymphoid progenitors within a narrow range of pseudotime values.

Finally, we discuss the running times of the methods used for the largest datasets (P1/2/3-BM, P1/2/3-Neu, Levine-32, Levine-13) and two randomly chosen smaller (P3-Mono, P1-Bcells). As shown in Supplementary Figure S29, we found that the runtime of TimeFlow scaled approximately linearly with the number of cells. The total runtime was less than 1 hour for the largest dataset used in this study. TinGa was significantly faster but less reliable in terms of cell ordering metrics. Both PAGA and Palantir also produced comparable, fast results. Although, Supplementary Figure S29 accounts for both the density and pseudotime module of TimeFlow, its runtime is dominated by the Real NVP module. We did not use GPUs to obtain these results, but their use can significantly accelerate the running time. In addition, TimeFlow yields robust pseudotime estimates by transferring weights across patients for the same biological process, without re-training the Real NVP model. This is important for applications where the number of patients increases drastically and computational time matters. A subset of patients may be selected for training and the learnt weights can be reused to directly evaluate the probability density function of the other patients, eliminating the need for training multiple times the Real NVP.

### 9 Section S9 Supplementary Tables

| Patient | Cell Population | Immature | Intermediate I | Intermediate II | Mature | Total Count |
|---|---|---|---|---|---|---|
| P1 | Mono | 7,703 | 12,629 | - | 18,924 | 39,256 |
| | Neu | 6,705 | 92,575 | 180,567 | 99,858 | 379,705 |
| | Ery | 6,012 | 1,801 | 24,204 | 30,377 | 62,394 |
| | B cells | 883 | 1,444 | - | 15,862 | 18,189 |
| | BM | - | - | - | - | 499,544 |
| P2 | Mono | 9,368 | 16,280 | - | 33,081 | 58,729 |
| | Neu | 5,320 | 48,227 | 242,539 | 80,014 | 376,100 |
| | Ery | 1,720 | 4,603 | 61,061 | 1,720 | 72,264 |
| | B cells | 451 | 970 | - | 3,694 | 5,115 |
| | BM | - | - | - | - | 512,208 |
| P3 | Mono | 11,425 | 31,655 | - | 24,347 | 67,427 |
| | Neu | 7,687 | 129,203 | 224,483 | 113,752 | 475,125 |
| | Ery | 4,384 | 3,049 | 11,344 | 9,060 | 27,837 |
| | B cells | 737 | 3,593 | - | 22,894 | 27,224 |
| | BM | - | - | - | - | 597,613 |

Table S1: Distribution of cell populations in our new hematopoietic single-cell flow cytometry datasets.

| Cell Population | Cell Counts |
|---|---|
| Mature CD4+ T cells | 13,964 |
| Erythroblast cells | 12,030 |
| Naive CD8+ T cells | 9,564 |
| Mature CD8+ T cells | 7,821 |
| Mature CD38lo B cells | 7,796 |
| Naive CD4+ T cells | 6,987 |
| CD11bhi Monocyte cells | 6,779 |
| NK cells | 3,864 |
| Megakaryocyte cells | 3,684 |
| Myelocyte cells | 3,025 |
| CD11bmid Monocyte cells | 1,278 |
| Pre-B II cells | 994 |
| CD11b- Monocyte cells | 912 |
| Mature CD38mid B cells | 608 |
| Immature B cells | 502 |
| unassigned | 85,297 |
| GMP cells | 73 |
| Plasmacytoid DC cells | 293 |
| HSC cells | 261 |
| Platelet cells | 5 |
| Pre-B I cells | 240 |
| CMP cells | 253 |
| MEP cells | 194 |
| MPP cells | 152 |
| Plasma cells | 468 |
| Total | 167,044 |

Table S2: Distribution of cell populations in Levine-13 hematopoietic dataset.

| Cell Population | Cell Counts |
|---|---|
| CD16- NK cells | 3,905 |
| CD34+CD38lo HSCs | 916 |
| CD34+CD38-CD123- HSPCs | 3,295 |
| CD34+CD38+CD123+ | 304 |
| CD16+ NK cells | 2,248 |
| Pro-B cells | 513 |
| Pre-B cells | 6,135 |
| Mature B cells | 16,520 |
| Plasma B cells | 330 |
| Monocytes | 21,099 |
| pDCs | 1,238 |
| Basophils | 1,207 |
| CD4 T cells | 26,366 |
| CD8 T cells | 20,108 |
| unassigned | 161,443 |
| Total | 265,627 |

Table S3: Distribution of cell populations in Levine-32 hematopoietic dataset.

| Cell Population | Cell Counts |
|---|---|
| Lymphoid progenitors | 20,835 |
| CD4 SP cells | 2,963 |
| CD8 SP cells | 1,202 |
| Total | 25,000 |

Table S4: Distribution of cell populations in the Wishbone T cell development dataset.

| Dataset | Modality | Process | Total Cells | CD Markers | Trajectory & Lineages | Trajectory Methods Used |
|---|---|---|---|---|---|---|
| P1 Mono | flow | human monopoiesis | 39,256 | 20 | linear (1) | TimeFlow, SCORPIUS, scShaper, Wanderlust, CytoTree |
| P1 Neu | flow | human neutropoiesis | 379,705 | 20 | linear (1) | TimeFlow, Comp1, ElpiLinear, DPT |
| P1 Ery | flow | human erythropoiesis | 62,394 | 20 | linear (1) | TimeFlow, SCORPIUS, scShaper, Wanderlust, CytoTree |
| P1 B-cells | flow | human B-cell lymphopoiesis | 18,189 | 20 | linear (1) | TimeFlow, SCORPIUS, scShaper, Wanderlust, CytoTree |
| P1 BM | flow | human hematopoiesis | 499,544 | 20 | branching (4) | TimeFlow, Palantir, TinGa, PAGA, ElPiGraph |
| P2 Mono | flow | human monopoiesis | 58,729 | 20 | linear (1) | TimeFlow, SCORPIUS, scShaper, Wanderlust, CytoTree |
| P2 Neu | flow | human neutropoiesis | 376,100 | 20 | linear (1) | TimeFlow, Comp1, ElpiLinear, DPT |
| P2 Ery | flow | human erythropoiesis | 72,264 | 20 | linear (1) | TimeFlow, SCORPIUS, scShaper, Wanderlust, CytoTree |
| P2 B-cells | flow | human B-cell lymphopoiesis | 5,115 | 20 | linear (1) | TimeFlow, SCORPIUS, scShaper, Wanderlust, CytoTree |
| P2 BM | flow | human hematopoiesis | 512,208 | 20 | branching (4) | TimeFlow, Palantir, TinGa, PAGA, ElPiGraph |
| P3 Mono | flow | human monopoiesis | 67,427 | 20 | linear (1) | TimeFlow, SCORPIUS, scShaper, Wanderlust, CytoTree |
| P3 Neu | flow | human neutropoiesis | 475,125 | 20 | linear (1) | TimeFlow, Comp1, ElpiLinear, DPT |
| P3 Ery | flow | human erythropoiesis | 27,837 | 20 | linear (1) | TimeFlow, SCORPIUS, scShaper, Wanderlust, CytoTree |
| P3 B-cells | flow | human B-cell lymphopoiesis | 27,224 | 20 | linear (1) | TimeFlow, SCORPIUS, scShaper, Wanderlust, CytoTree |
| P3 BM | flow | human hematopoiesis | 597,613 | 20 | branching (4) | TimeFlow, Palantir, TinGa, PAGA, ElPiGraph |
| Levine-13 | mass | human hematopoiesis | 167,044 | 13 | branching (2) | TimeFlow, Palantir, TinGa, PAGA, ElPiGraph |
| Levine-32 | mass | human hematopoiesis | 265,627 | 32 | branching (2) | TimeFlow, Palantir, TinGa, PAGA, ElPiGraph |
| Wishbone T-cell | mass | mouse hematopoiesis | 25,000 | 13 | branching (2) | TimeFlow, Palantir, TinGa, PAGA, ElPiGraph |

Table S5: Summary of datasets and methods applied in this study for comparisons.

| Dataset | Cell Populations | Kendall's Tau Correlation | | | | | Spearman's Rank Correlation | | | | | Cell Accuracy | | | | | Cell Stage Consistency | | | | |
|---|---|---|---|---|---|---|---|---|---|---|---|---|---|---|---|---|---|---|---|---|---|
| | | TimeFlow | PAGA | Palantir | TinGa | ElpIGraph | TimeFlow | PAGA | Palantir | TinGa | ElpIGraph | TimeFlow | PAGA | Palantir | TinGa | ElpIGraph | TimeFlow | PAGA | Palantir | TinGa | ElpIGraph |
| P1 BM | Mono | 0.68 | 0.73 | 0.74 | 0.52 | 0.7 | 0.81 | 0.86 | 0.87 | 0.6 | 0.86 | 0.8 | 0.89 | 0.91 | 0.68 | 0.86 | 1 | 1 | 1 | 0.89 | 1 |
| | Neu | 0.7 | 0.72 | 0.34 | 0.77 | 0.77 | 0.83 | 0.85 | 0.49 | 0.89 | 0.89 | 0.76 | 0.79 | 0.5 | 0.89 | 0.87 | 1 | 1 | 0.66 | 0.99 | 1 |
| | Ery | 0.71 | 0.71 | 0.76 | 0.1 | -0.2 | 0.84 | 0.84 | 0.89 | 0.15 | -0.17 | 0.86 | 0.91 | 0.97 | 0.39 | 0.23 | 1 | 1 | 1 | 0.69 | 0.66 |
| | B-cells | 0.45 | 0.47 | 0.47 | 0.38 | 0.58 | 0.54 | 0.57 | 0.57 | 0.42 | 0.63 | 0.94 | 0.99 | 0.99 | 0.86 | 0.91 | 1 | 1 | 1 | 0.45 | 0.91 |
| P2-BM | Mono | 0.63 | 0.74 | 0.69 | 0.55 | 0.67 | 0.76 | 0.86 | 0.81 | 0.63 | 0.76 | 0.79 | 0.93 | 0.91 | 0.75 | 0.88 | 1 | 1 | 1 | 0.97 | 1 |
| | Neu | 0.6 | 0.6 | 0.71 | 0.66 | 0.69 | 0.72 | 0.73 | 0.84 | 0.79 | 0.81 | 0.78 | 0.81 | 0.95 | 0.89 | 0.93 | 1 | 1 | 1 | 0.98 | 1 |
| | Ery | 0.45 | 0.48 | 0.36 | 0.43 | 0.49 | 0.55 | 0.58 | 0.45 | 0.48 | 0.57 | 0.95 | 0.97 | 0.93 | 0.9 | 0.92 | 0.91 | 1 | 0.91 | 0.78 | 0.91 |
| | B-cells | 0.64 | 0.63 | 0.65 | 0.52 | 0.67 | 0.76 | 0.76 | 0.77 | 0.62 | 0.69 | 0.95 | 0.95 | 0.99 | 0.46 | 0.74 | 1 | 1 | 1 | 0.57 | 0.81 |
| P3-BM | Mono | 0.72 | 0.76 | 0.75 | 0.5 | 0.75 | 0.85 | 0.87 | 0.88 | 0.59 | 0.85 | 0.85 | 0.92 | 0.91 | 0.61 | 0.85 | 1 | 1 | 1 | 0.83 | 1 |
| | Neu | 0.54 | -0.58 | 0.78 | 0.7 | 0.73 | 0.66 | -0.69 | 0.9 | 0.82 | 0.85 | 0.62 | 0.35 | 0.93 | 0.78 | 0.77 | 1 | 0.1 | 1 | 0.94 | 1 |
| | Ery | 0.8 | 0.73 | 0.75 | 0.15 | 0.19 | 0.92 | 0.58 | 0.87 | 0.24 | 0.3 | 0.89 | 0.88 | 0.9 | 0.33 | 0.33 | 1 | 0.99 | 1 | 0.36 | 0.66 |
| | B-cells | 0.49 | -0.09 | 0.51 | 0.2 | 0.59 | 0.61 | -0.07 | 0.62 | 0.22 | 0.65 | 0.96 | 0.74 | 0.97 | 0.77 | 0.97 | 1 | 0.83 | 1 | 0.14 | 1 |
| Levine-13 | B-cells | 0.49 | 0.57 | 0.6 | 0.57 | 0.5 | 0.6 | 0.71 | 0.73 | 0.64 | 0.6 | 0.71 | 0.79 | 0.82 | 0.76 | 0.77 | 0.87 | 0.51 | 0.7 | 0.56 | 0.8 |
| | Mono | 0.36 | 0.55 | -0.63 | 0.1 | 0.5 | 0.45 | 0.66 | -0.75 | 0.11 | 0.61 | 0.64 | 0.69 | 0.39 | 0.59 | 0.69 | 0.7 | 0.55 | 0 | 0.45 | 0.74 |
| Levine-32 | B-cells | 0.64 | 0.74 | 0.49 | 0.78 | 0.61 | 0.77 | 0.86 | 0.72 | 0.88 | 0.73 | 0.75 | 0.92 | 0.62 | 0.87 | 0.68 | 0.85 | 0.96 | 0.08 | 0.82 | 0.5 |
| | Mono | 0.51 | 0.53 | 0.51 | 0.54 | 0.52 | 0.63 | 0.64 | 0.63 | 0.62 | 0.65 | 0.89 | 0.95 | 0.91 | 0.85 | 0.9 | 0.81 | 0.91 | 0.81 | 0.54 | 0.81 |
| Wishbone | CD4 T-cells | 0.42 | 0.46 | 0.46 | 0.33 | 0.46 | 0.52 | 0.56 | 0.57 | 0.4 | 0.56 | 0.94 | 0.99 | 0.99 | 0.93 | 0.98 | 1 | 1 | 1 | 0.85 | 1 |
| | CD8 T-cells | 0.29 | 0.31 | 0.32 | 0.2 | 0.32 | 0.35 | 0.38 | 0.39 | 0.24 | 0.39 | 0.97 | 0.99 | 0.99 | 0.95 | 0.98 | 1 | 1 | 1 | 0.75 | 1 |
| **Average scores across all lineages** | - | **0.55** | **0.5** | **0.51** | **0.44** | **0.53** | **0.66** | **0.58** | **0.62** | **0.51** | **0.62** | **0.83** | **0.85** | **0.93** | **0.73** | **0.79** | **0.95** | **0.88** | **0.84** | **0.69** | **0.87** |
| **Average scores (aggregated per dataset)** | - | **0.52** | **0.5** | **0.45** | **0.43** | **0.51** | **0.63** | **0.59** | **0.56** | **0.50** | **0.61** | **0.82** | **0.86** | **0.84** | **0.75** | **0.8** | **0.93** | **0.86** | **0.77** | **0.68** | **0.85** |

Table S6: Evaluation scores for all datasets with branching trajectories. Average scores are provided based on 1) the mean calculated across all cell lineages and 2) across datasets, with the dataset-specific means computed first.

| Dataset | Cell Populations | Kendall's Tau Correlation | | | | | | | | Spearman's Rank Correlation | | | | | | | |
|---|---|---|---|---|---|---|---|---|---|---|---|---|---|---|---|---|---|
| | | TimeFlow | Wanderlust | SCORPIUS | CytoTree | scShaper | Comp1 | ElpiLinear | DPT | TimeFlow | Wanderlust | SCORPIUS | CytoTree | scShaper | Comp1 | ElpiLinear | DPT |
| P1 | Mono | 0.72 | 0.19 | 0.71 | 0.72 | 0.7 | 0.73 | 0.72 | 0.75 | 0.85 | 0.31 | 0.84 | 0.79 | 0.84 | 0.86 | 0.86 | 0.88 |
| | Neu | 0.66 | - | - | - | - | 0.77 | 0.76 | 0.69 | 0.81 | - | - | - | - | 0.9 | 0.89 | 0.82 |
| | Ery | 0.68 | 0.67 | -0.21 | 0.36 | 0.63 | -0.04 | 0.76 | 0.77 | 0.8 | 0.79 | -0.2 | 0.4 | 0.77 | -0.006 | 0.89 | 0.9 |
| | B-cells | 0.45 | 0.46 | 0.45 | 0.54 | 0.35 | 0.4 | 0.47 | 0.47 | 0.55 | 0.56 | 0.55 | 0.59 | 0.44 | 0.5 | 0.57 | 0.57 |
| P2 | Mono | 0.71 | -0.06 | 0.72 | 0.79 | 0.67 | 0.75 | 0.73 | 0.71 | 0.84 | -0.03 | 0.86 | 0.86 | 0.83 | 0.88 | 0.87 | 0.83 |
| | Neu | 0.7 | - | - | - | - | 0.71 | 0.68 | 0.06 | 0.62 | - | - | - | - | 0.83 | 0.81 | 0.01 |
| | Ery | 0.5 | 0.15 | 0.42 | 0.44 | 0.47 | 0.39 | 0.46 | 0.5 | 0.6 | 0.2 | 0.52 | 0.47 | 0.72 | 0.76 | 0.56 | 0.6 |
| | B-cells | 0.62 | 0.64 | 0.62 | 0.71 | 0.6 | 0.63 | 0.65 | 0.65 | 0.74 | 0.77 | 0.74 | 0.77 | 0.72 | 0.76 | 0.77 | 0.77 |
| P3 | Mono | 0.7 | 0.01 | 0.68 | 0.7 | 0.3 | 0.71 | 0.76 | 0.75 | 0.83 | 0.09 | 0.8 | 0.78 | 0.35 | 0.84 | 0.89 | 0.88 |
| | Neu | 0.65 | - | - | - | - | 0.77 | 0.73 | 0.64 | 0.78 | - | - | - | - | 0.9 | 0.85 | 0.77 |
| | Ery | 0.73 | 0.29 | 0.3 | 0.46 | 0.5 | 0.56 | 0.81 | 0.82 | 0.87 | 0.4 | 0.46 | 0.55 | 0.58 | 0.68 | 0.93 | 0.94 |
| | B-cells | 0.5 | 0.35 | 0.38 | 0.56 | 0.5 | 0.51 | 0.51 | 0.51 | 0.6 | 0.42 | 0.46 | 0.62 | 0.61 | 0.62 | 0.61 | 0.62 |
| **Average Scores (P1/2/3-Mono/Ery/B-cells)** | - | **0.62** | **0.38** | **0.58** | **0.58** | **0.45** | - | - | - | **0.74** | **0.38** | **0.55** | **0.64** | **0.63** | - | - | - |
| **Average Scores (P1/2/3-Mono/Neu/Ery/B-cells)** | - | **0.61** | - | - | - | - | **0.57** | **0.66** | **0.6** | **0.73** | - | - | - | - | **0.68** | **0.79** | **0.71** |

Table S7: Kendall's Tau and Spearman's correlation for all datasets with linear trajectories. Average scores are provided based on 1) the mean calculated across cell lineages for P1/2/3-Mono/Ery/B-cells and 2) the mean calculated across cell lineages for P1/2/3-Mono/Neu/Ery/B-cells.

| Dataset | Cell Populations | Cell Accuracy | | | | | | | | Cell Stage Consistency | | | | | | | |
|---|---|---|---|---|---|---|---|---|---|---|---|---|---|---|---|---|---|
| | | TimeFlow | Wanderlust | SCORPIUS | CytoTree | scShaper | Comp1 | ElpiLinear | DPT | TimeFlow | Wanderlust | SCORPIUS | CytoTree | scShaper | Comp1 | ElpiLinear | DPT |
| P1 | Mono | 0.84 | 0.45 | 0.89 | 0.76 | 0.83 | 0.85 | 0.88 | 0.89 | 1 | 0.33 | 0.99 | 1 | 0.95 | 1 | 1 | 1 |
| | Neu | 0.73 | - | - | - | - | 0.87 | 0.89 | 0.75 | 1 | - | - | - | - | 0.91 | 1 | 1 |
| | Ery | 0.83 | 0.9 | 0.18 | 0.54 | 0.85 | 0.33 | 0.98 | 0.98 | 1 | 1 | 0.52 | 0.77 | 0.79 | 0.66 | 1 | 1 |
| | B-cells | 0.93 | 0.97 | 0.96 | 0.94 | 0.87 | 0.87 | 0.98 | 0.99 | 1 | 1 | 1 | 1 | 0.78 | 0.81 | 1 | 1 |
| P2 | Mono | 0.87 | 0.43 | 0.91 | 0.88 | 0.8 | 0.94 | 0.9 | 0.93 | 1 | 0.81 | 1 | 1 | 0.86 | 1 | 1 | 1 |
| | Neu | 0.68 | - | - | - | - | 0.92 | 0.94 | 0.59 | 1 | - | - | - | - | 0.91 | 1 | 0.18 |
| | Ery | 0.93 | 0.86 | 0.91 | 0.9 | 0.96 | 0.92 | 0.93 | 0.98 | 0.91 | 0.18 | 0.91 | 0.66 | 0.96 | 0.91 | 0.91 | 1 |
| | B-cells | 0.9 | 0.98 | 0.95 | 0.91 | 0.96 | 0.92 | 0.99 | 0.99 | 1 | 1 | 1 | 1 | 1 | 1 | 1 | 1 |
| P3 | Mono | 0.8 | 0.31 | 0.84 | 0.75 | 0.56 | 0.81 | 0.93 | 0.92 | 1 | 0.33 | 0.92 | 1 | 0.47 | 1 | 1 | 1 |
| | Neu | 0.69 | - | - | - | - | 0.88 | 0.78 | 0.69 | 1 | - | - | - | - | 0.91 | 1 | 1 |
| | Ery | 0.76 | 0.58 | 0.41 | 0.48 | 0.66 | 0.65 | 0.95 | 0.97 | 1 | 0.91 | 0.54 | 0.66 | 0.62 | 1 | 1 | 1 |
| | B-cells | 0.92 | 0.92 | 0.87 | 0.9 | 0.96 | 0.96 | 0.98 | 0.99 | 1 | 0.33 | 0.85 | 0.81 | 0.91 | 1 | 1 | 1 |
| **Average Scores (P1/2/3-Mono/Ery/B-cells)** | - | **0.86** | **0.71** | **0.76** | **0.78** | **0.82** | - | - | - | **1** | **0.65** | **0.86** | **0.88** | **0.82** | - | - | - |
| **Average Scores (P1/2/3-Mono/Neu/Ery/B-cells)** | - | **0.82** | - | - | - | - | **0.82** | **0.92** | **0.88** | **1** | - | - | - | - | **0.92** | **1** | **0.93** |

Table S8: Cell Accuracy and Cell Stage Consistency for all datasets with linear trajectories. Average scores are provided based on 1) the mean calculated across cell lineages for P1/2/3-Mono/Ery/B-cells and 2) the mean calculated across cell lineages for P1/2/3-Mono/Neu/Ery/B-cells.

| Methods | Datasets | | | | | | | | | | | | | | | | | | |
|---|---|---|---|---|---|---|---|---|---|---|---|---|---|---|---|---|---|---|---|
| | P1 Mono | P1 Ery | P1 Neu | P1 B-cells | P2 Mono | P2 Ery | P2 Neu | P2 B-cells | P3 Mono | P3 Ery | P3 Neu | P3 B-cells | Levine-13 B-cells | Levine-13 Mono | Levine-32 B-cells | Levine-32 Mono | Wishbone CD4 | Wishbone CD8 | **Average** |
| TimeFlow (linear) | 4.26 | 3.89 | 3.72 | 3.92 | 4.28 | 3.73 | 3.75 | 4.28 | 4.3 | 3.95 | 3.77 | 3.96 | - | - | - | - | - | - | **3.98** |
| ElPiLinear | 4.23 | 4.25 | 4.38 | 4.14 | 4.21 | 4.28 | 4.35 | 4.42 | 4.44 | 4.15 | 4.36 | 4.44 | - | - | - | - | - | - | **4.3** |
| SCORPIUS | 4.31 | 4.17 | - | 4.3 | 4.39 | 4.2 | - | 4.41 | 4.35 | 4.14 | - | 4.45 | - | - | - | - | - | - | **4.3** |
| Wanderlust | 3.61 | 3.46 | - | 4 | 2.98 | 3.7 | - | 4.26 | 3.46 | 4.12 | - | 4.1 | - | - | - | - | - | - | **3.74** |
| DPT | 3.74 | 2.55 | 0.62 | 2.07 | 3.51 | 2.06 | 1.37 | 2.87 | 3.7 | 3.44 | 0.38 | 1.74 | - | - | - | - | - | - | **2.34** |
| sc-Shaper | 3.91 | 3.71 | - | 3.57 | 4.03 | 4.12 | - | 3.83 | 3.91 | 3.77 | - | 3.56 | - | - | - | - | - | - | **3.82** |
| Comp1 | 4.27 | 3.43 | 4.31 | 4.22 | 4.23 | 3.35 | 4.25 | 4.34 | 4.25 | 3.44 | 4.36 | 4.14 | - | - | - | - | - | - | **4.05** |
| CytoTree | 1.86 | 1.3 | - | 1.53 | 1.87 | 1.36 | - | 1.66 | 1.89 | 1.64 | - | 1.55 | - | - | - | - | - | - | **1.63** |
| TimeFlow (branching) | 4.06 | 3.84 | 3.85 | 3.83 | 4.13 | 3.74 | 3.68 | 4.33 | 4.19 | 4.14 | 3.79 | 3.86 | 3.96 | 3.86 | 4 | 4.02 | 3.83 | 3.7 | **3.93** |
| ElPiGraph | 3.15 | 2.18 | 3.64 | 1.17 | 2.85 | 2.16 | 3.4 | 0.80 | 3.21 | 2.04 | 3.31 | 1.62 | 2.5 | 2.68 | 3.14 | 3.5 | 3.37 | 3.25 | **2.67** |
| Palantir | 3.42 | 2.35 | 2.61 | 1.87 | 2.85 | 1.8 | 1.52 | 2.72 | 3.02 | 2.91 | 1.58 | 1.96 | 3.42 | 3.39 | 3.14 | 2.73 | 1.8 | 1.3 | **2.47** |
| PAGA | 3.1 | 2.59 | 1.27 | 3.04 | 2.89 | 2.35 | 1.17 | 3.87 | 3.98 | 3.95 | 0.8 | 2.38 | 3.88 | 3.52 | 3.91 | 3.29 | 1.14 | 0.087 | **2.62** |
| TinGa | 2.55 | 2.29 | 3.63 | 2.25 | 2.65 | 1.52 | 3.72 | 2.9 | 2.89 | 2.69 | 3.53 | 1.01 | 1.79 | 2.45 | 2.77 | 2.6 | 3.76 | 3.68 | **2.7** |

Table S9: Shannon Entropy scores of pseudotime for all methods across all trajectories. Average scores are provided based on the number of trajectories to which a method was applied.

| Experiment | BM Mono | BM Neu | BM Ery | BM B-cells | Mean Cell Stage Consistency (↑) |
|---|---|---|---|---|---|
| P1 trained on P1 | 1.00 | 1.00 | 1.00 | 1.00 | **1.00 ± 0.00** |
| P1 trained on P2 | 1.00 | 0.91 | 1.00 | 1.00 | 0.98 ± 0.04 |
| P1 trained on P3 | 1.00 | 1.00 | 1.00 | 1.00 | 1.00 ± 0.00 |
| P2 trained on P2 | 1.00 | 1.00 | 0.91 | 1.00 | **0.98 ± 0.04** |
| P2 trained on P1 | 1.00 | 0.91 | 0.91 | 1.00 | 0.96 ± 0.04 |
| P2 trained on P3 | 1.00 | 1.00 | 1.00 | 1.00 | 1.00 ± 0.00 |
| P3 trained on P3 | 1.00 | 1.00 | 1.00 | 1.00 | **1.00 ± 0.00** |
| P3 trained on P1 | 1.00 | 1.00 | 0.66 | 1.00 | 0.92 ± 0.15 |
| P3 trained on P2 | 1.00 | 1.00 | 1.00 | 1.00 | 1.00 ± 0.00 |

Table S10: TimeFlow pseudotime analysis with weight transferring across patients for the branching trajectories of the P1/2/3-BM Mono, Neu, Ery, B-cells datasets. TimeFlow Cell Stage Consistency scores for each cell population and overall mean for all populations. In bold the original scores where density and pseudotime modules were applied on the same patient.

| Experiment | Linear Mono | Linear Neu | Linear Ery | Linear B-cells | Mean Cell Stage Consistency (↑) |
|---|---|---|---|---|---|
| P1 trained on P1 | 1.00 | 1.00 | 1.00 | 1.00 | **1.00 ± 0.00** |
| P1 trained on P2 | 1.00 | 0.91 | 1.00 | 1.00 | 0.98 ± 0.04 |
| P1 trained on P3 | 1.00 | 0.91 | 0.91 | 1.00 | 0.96 ± 0.04 |
| P2 trained on P2 | 1.00 | 1.00 | 0.91 | 1.00 | **0.98 ± 0.04** |
| P2 trained on P1 | 1.00 | 1.00 | 0.91 | 1.00 | 0.98 ± 0.04 |
| P2 trained on P3 | 1.00 | 0.91 | 0.91 | 1.00 | 0.96 ± 0.04 |
| P3 trained on P3 | 1.00 | 1.00 | 1.00 | 1.00 | **1.00 ± 0.00** |
| P3 trained on P1 | 1.00 | 1.00 | 1.00 | 0.81 | 0.95 ± 0.08 |
| P3 trained on P2 | 1.00 | 1.00 | 1.00 | 0.81 | 0.95 ± 0.08 |

Table S11: TimeFlow pseudotime analysis with weight transferring across patients for the linear trajectories of the P1/2/3- Mono, Neu, Ery, B cells datasets. TimeFlow Cell Stage Consistency scores for each cell population and overall mean for all populations. In bold the original scores where density and pseudotime modules were applied on the same patient.

| Patient | Trained w/o Immat Mono | Trained w/o Interm Mono | Trained w/o Mat Mono |
|---|---|---|---|
| P1 | 1.00 | 1.00 | 1.00 |
| P2 | 1.00 | 1.00 | 1.00 |
| P3 | 1.00 | 1.00 | 1.00 |

Table S12: TimeFlow pseudotime analysis with weight transferring from discontinuous to full linear trajectories for the P1/2/3-Mono datasets. TimeFlow Cell Stage Consistency scores refer to full monocytic trajectories. Each column specifies which cell population was absent during density fitting.

| Patient | Trained w/o Immat Neu | Trained w/o Interm I Neu | Trained w/o Interm II Neu | Trained w/o Mat Neu |
|---|---|---|---|---|
| P1 | 0.91 | 1.00 | 0.91 | 1.00 |
| P2 | 1.00 | 1.00 | 0.91 | 1.00 |
| P3 | 1.00 | 0.91 | 1.00 | 1.00 |

Table S13: TimeFlow pseudotime analysis with weight transferring from discontinuous to full linear trajectories for the P1/2/3-Neu datasets. TimeFlow Cell Stage Consistency scores refer to full neutrophilic trajectories. Each column specifies which cell population was absent during density fitting.

| Patient | Trained w/o Immat Ery | Trained w/o Interm I Ery | Trained w/o Interm II Ery | Trained w/o Mat Ery |
|---|---|---|---|---|
| P1 | 1.00 | 0.91 | 1.00 | 1.00 |
| P2 | 1.00 | 0.91 | 0.91 | 1.00 |
| P3 | 1.00 | 1.00 | 1.00 | 0.91 |

Table S14: TimeFlow pseudotime analysis with weight transferring from discontinuous to full linear trajectories for the P1/2/3-Ery datasets. TimeFlow Cell Stage Consistency scores refer to full erythroid trajectories. Each column specifies which cell population was absent during density fitting.

| Patient | Trained w/o Immat B cells | Trained w/o Interm B cells | Trained w/o Mat B cells |
|---|---|---|---|
| P1 | 1.00 | 1.00 | 1.00 |
| P2 | 0.81 | 1.00 | 1.00 |
| P3 | 0.81 | 0.81 | 1.00 |

Table S15: TimeFlow pseudotime analysis with weight transferring from discontinuous to full linear trajectories for the P1/2/3-B-cells datasets. TimeFlow Cell Stage Consistency scores refer to full B cell trajectories. Each column specifies which cell population was absent during density fitting.

| Real NVP Configuration | Coupling Layers | Hidden Neurons | Batch Size |
|---|---|---|---|
| 1 | 8 | 128 | 512 |
| 2 | 8 | 256 | 512 |
| 3 | 10 | 128 | 512 |
| 4 | 10 | 256 | 512 |

Table S16: Hyper-parameters for different Real NVP architectures tested on the branching trajectories of P1/2/3-BM datasets.

| Real NVP Configuration | Coupling Layers | Hidden Neurons | Batch Size |
|---|---|---|---|
| 1 | 8 | 128 | 256 |
| 2 | 8 | 256 | 256 |
| 3 | 10 | 128 | 256 |
| 4 | 10 | 256 | 256 |
| 5 | 8 | 128 | 512 |
| 6 | 8 | 256 | 512 |
| 7 | 10 | 128 | 512 |
| 8 | 10 | 256 | 512 |

Table S17: Hyper-parameters for different Real NVP architectures tested on the linear trajectories of P1/2/3-Mono/Neu/Ery/B-cells datasets.

| Real NVP Configuration | BM Mono | BM Neu | BM Ery | BM B-cells | Mean Cell Stage Consistency (↑) |
|---|---|---|---|---|---|
| 1 | 1.00 | 0.97 | 0.94 | 1.00 | 0.98 ± 0.00 |
| 2 | 1.00 | 0.90 | 0.94 | 1.00 | 0.98 ± 0.00 |
| 3 | 1.00 | 0.90 | 0.83 | 1.00 | 0.93 ± 0.08 |
| 4 | 1.00 | 0.97 | 0.85 | 1.00 | 0.95 ± 0.01 |

Table S18: Comparisons of different Real NVP configurations for the branching trajectories of the P1/2/3-BM datasets. TimeFlow Cell Stage Consistency scores of each BM population are averaged across the three patients. An overall mean score for each configuration across all cell populations is also provided.

| Real NVP Configuration | Linear Mono | Linear Ery | Linear B-cells | Mean Cell Stage Consistency (↑) |
|---|---|---|---|---|
| 1 | 1.00 | 0.97 | 0.94 | 0.97 ± 0.00 |
| 2 | 1.00 | 1.00 | 0.94 | 0.98 ± 0.00 |
| 3 | 1.00 | 0.97 | 1.00 | 0.99 ± 0.00 |
| 4 | 1.00 | 0.97 | 0.94 | 0.97 ± 0.00 |
| 5 | 1.00 | 0.97 | 0.94 | 0.97 ± 0.00 |
| 6 | 1.00 | 0.97 | 0.94 | 0.97 ± 0.00 |
| 7 | 1.00 | 1.00 | 0.94 | 0.98 ± 0.00 |
| 8 | 1.00 | 0.97 | 0.94 | 0.97 ± 0.00 |

Table S19: Comparisons for different Real NVP configurations for the linear trajectories of the P1/2/3-Mono/Neu/Ery/B-cells datasets. TimeFlow Cell Stage Consistency scores of each population are averaged across the three patients. An overall mean score for each configuration across all cell populations is also provided.

| k-NN | P1/2/3-BM Mono | P1/2/3-BM Neu | P1/2/3-BM Ery | P1/2/3-BM B-cells | Mean Cell Stage Consistency (↑) |
|---|---|---|---|---|---|
| 3 | 1.00 | 0.97 | 1.00 | 1.00 | 0.99 ± 0.01 |
| 4 | 1.00 | 1.00 | 1.00 | 1.00 | 1.00 ± 0.00 |
| 5 | 1.00 | 1.00 | 0.97 | 1.00 | 0.99 ± 0.01 |
| 6 | 1.00 | 1.00 | 0.97 | 1.00 | 0.99 ± 0.01 |
| 7 | 1.00 | 1.00 | 0.97 | 1.00 | 0.99 ± 0.01 |
| 8 | 1.00 | 1.00 | 0.97 | 1.00 | 0.99 ± 0.01 |
| 9 | 1.00 | 1.00 | 0.97 | 1.00 | 0.99 ± 0.01 |
| 10 | 1.00 | 1.00 | 0.97 | 1.00 | 0.99 ± 0.01 |
| 11 | 1.00 | 1.00 | 0.86 | 1.00 | 0.96 ± 0.06 |
| 12 | 1.00 | 1.00 | 0.97 | 0.88 | 0.96 ± 0.05 |
| 13 | 1.00 | 1.00 | 0.97 | 0.94 | 0.97 ± 0.02 |
| 14 | 1.00 | 1.00 | 0.97 | 1.00 | 0.99 ± 0.01 |
| 15 | 1.00 | 1.00 | 0.97 | 1.00 | 0.99 ± 0.01 |
| 16 | 1.00 | 1.00 | 1.00 | 1.00 | 1.00 ± 0.00 |
| 17 | 1.00 | 1.00 | 0.97 | 1.00 | 0.99 ± 0.01 |
| 18 | 1.00 | 1.00 | 1.00 | 1.00 | 1.00 ± 0.00 |
| 19 | 1.00 | 1.00 | 0.94 | 1.00 | 0.98 ± 0.03 |
| 20 | 1.00 | 1.00 | 1.00 | 1.00 | 1.00 ± 0.00 |

Table S20: Comparisons for different values of k Nearest Neighbours for the branching trajectories of the P1/2/3-BM datasets. TimeFlow Cell Stage Consistency scores of each population are averaged across the three patients. An overall mean score for each value of k across all cell populations is also provided.

| k-NN | P1/2/3-BM Mono | P1/2/3-BM Neu | P1/2/3-BM Ery | P1/2/3-BM B-cells | Mean Cell Stage Consistency (↑) |
|---|---|---|---|---|---|
| 3 | 1.00 | 0.97 | 1.00 | 1.00 | 0.99 ± 0.01 |
| 4 | 1.00 | 1.00 | 0.97 | 1.00 | 0.99 ± 0.01 |
| 5 | 1.00 | 1.00 | 0.97 | 1.00 | 0.99 ± 0.01 |
| 6 | 1.00 | 1.00 | 0.97 | 0.94 | 0.97 ± 0.02 |
| 7 | 1.00 | 0.97 | 0.97 | 0.94 | 0.97 ± 0.02 |
| 8 | 1.00 | 1.00 | 1.00 | 0.94 | 0.98 ± 0.03 |
| 9 | 1.00 | 0.97 | 0.97 | 0.94 | 0.97 ± 0.02 |
| 10 | 1.00 | 0.97 | 0.97 | 0.94 | 0.97 ± 0.02 |
| 11 | 1.00 | 0.90 | 0.97 | 0.94 | 0.95 ± 0.04 |
| 12 | 1.00 | 0.97 | 0.97 | 0.94 | 0.97 ± 0.02 |
| 13 | 1.00 | 0.97 | 0.90 | 0.94 | 0.95 ± 0.04 |
| 14 | 1.00 | 1.00 | 0.97 | 1.00 | 0.99 ± 0.01 |
| 15 | 1.00 | 1.00 | 0.90 | 0.94 | 0.96 ± 0.04 |
| 16 | 1.00 | 1.00 | 0.97 | 0.88 | 0.96 ± 0.05 |
| 17 | 1.00 | 1.00 | 0.90 | 0.88 | 0.94 ± 0.06 |
| 18 | 1.00 | 1.00 | 0.97 | 0.88 | 0.96 ± 0.05 |
| 19 | 1.00 | 1.00 | 0.97 | 0.88 | 0.96 ± 0.05 |
| 20 | 1.00 | 1.00 | 0.97 | 0.88 | 0.96 ± 0.05 |

Table S21: Comparisons for different values of k Nearest Neighbours for the linear trajectories of the P1/2/3-Mono/Neu/Ery/B-cells datasets. TimeFlow Cell Stage Consistency scores of each population are averaged across the three patients. An overall mean score for each value of k across all cell populations is also provided.
