## Supplementary Figures for "TimeFlow: a density-driven pseudotime method for flow cytometry data analysis"

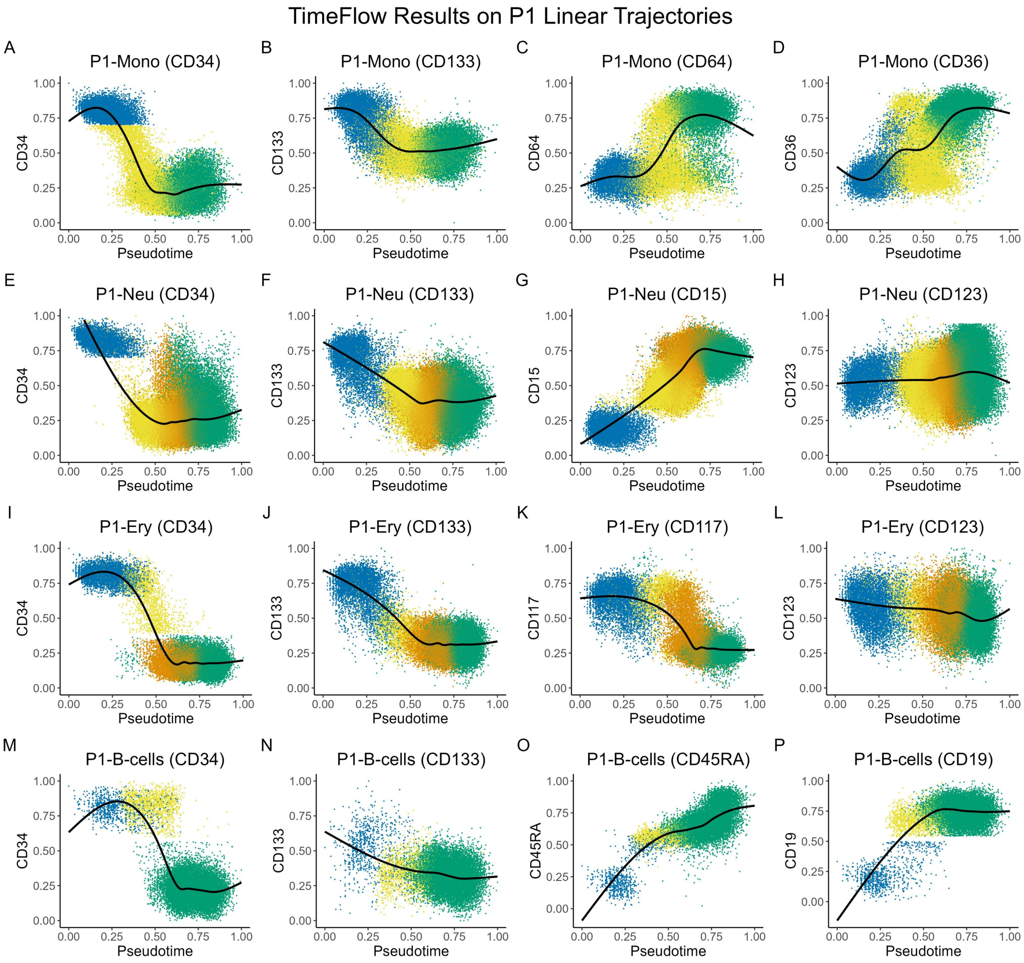


Figure S1: Early hematopoietic markers (CD34 and CD133) and other markers along pseudotime inferred by TimeFlow. The datasets used are: P1-Mono, P1-Neu, P1-Ery, P1-Bcells. Each scatterplot dot represents a cell, coloured by its maturation stage. GAM models were fitted without use of labels. Stacked bar plots show the cell distribution (%) in each of their known maturation stages, following pseudotime ordering. Gaps in the marker expression (y-axis) between consecutive cell populations are a consequence of using rectangular gates during dataset preparation. (A-D) Marker dynamics of CD34, CD133, CD14 and CD64 for P1-Mono. (E-H) Marker dynamics of CD34, CD133, CD15 and CD123 for P1-Neu. (I-L) Marker dynamics of CD34, CD133, CD117 and CD123 for P1-Ery. (M-P) Marker dynamics of CD34, CD133, CD45RA and CD19.

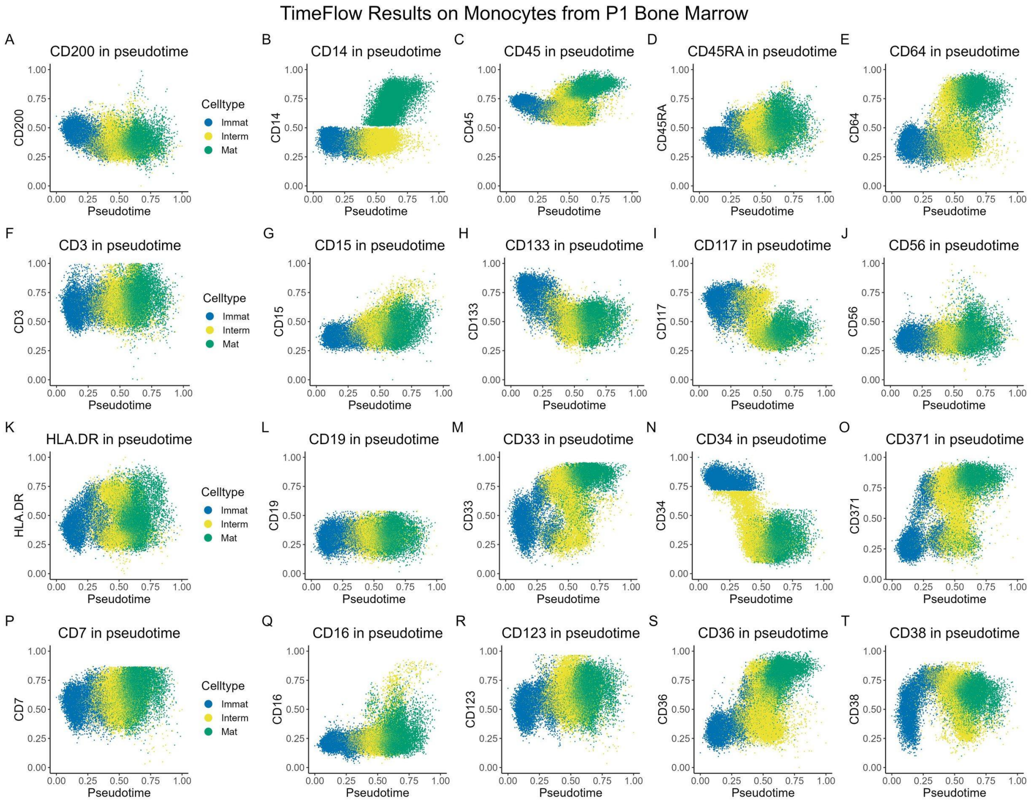


Figure S2: TimeFlow pseudotemporal orderings of Monocytes using the P1-BM dataset. (A-T) Scatterplots showing the expression levels of each marker along pseudotime. Each scatterplot dot represents a cell, coloured by its maturation stage. Gaps in the marker expression (y-axis) between consecutive cell populations are a consequence of using rectangular gates during dataset preparation.

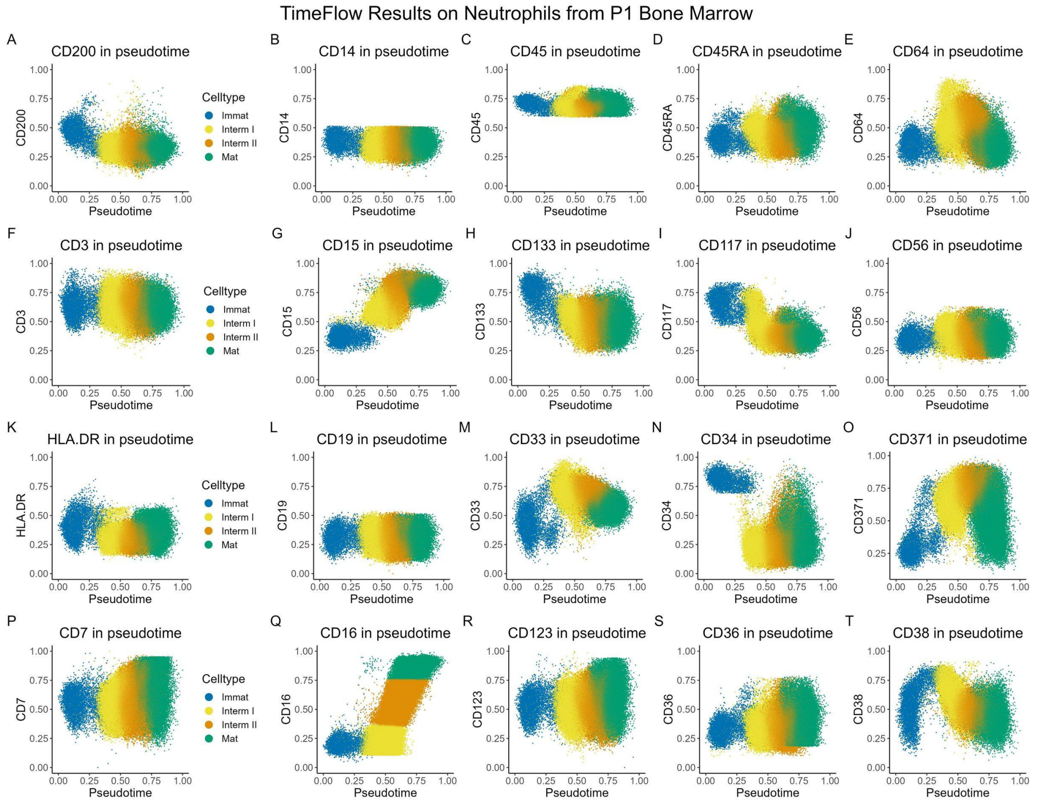


Figure S3: TimeFlow pseudotemporal orderings of Neutrophils using the P1-BM dataset. (A-T) Scatterplots showing the expression levels of each marker along pseudotime. Each scatterplot dot represents a cell, coloured by its maturation stage. Gaps in the marker expression (y-axis) between consecutive cell populations are a consequence of using rectangular gates during dataset preparation.

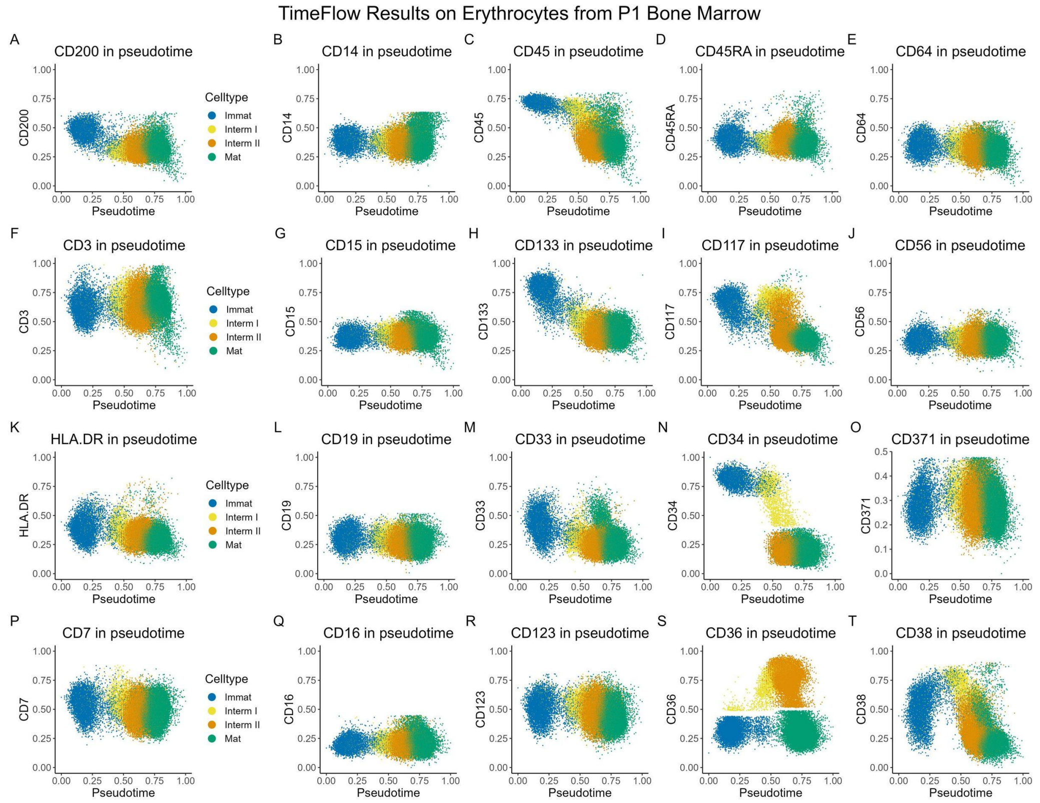


Figure S4: TimeFlow pseudotemporal orderings of Erythrocytes using the P1-BM dataset. (A-T) Scatterplots showing the expression levels of each marker along pseudotime. Each scatterplot dot represents a cell, coloured by its maturation stage. Gaps in the marker expression (y-axis) between consecutive cell populations are a consequence of using rectangular gates during dataset preparation.

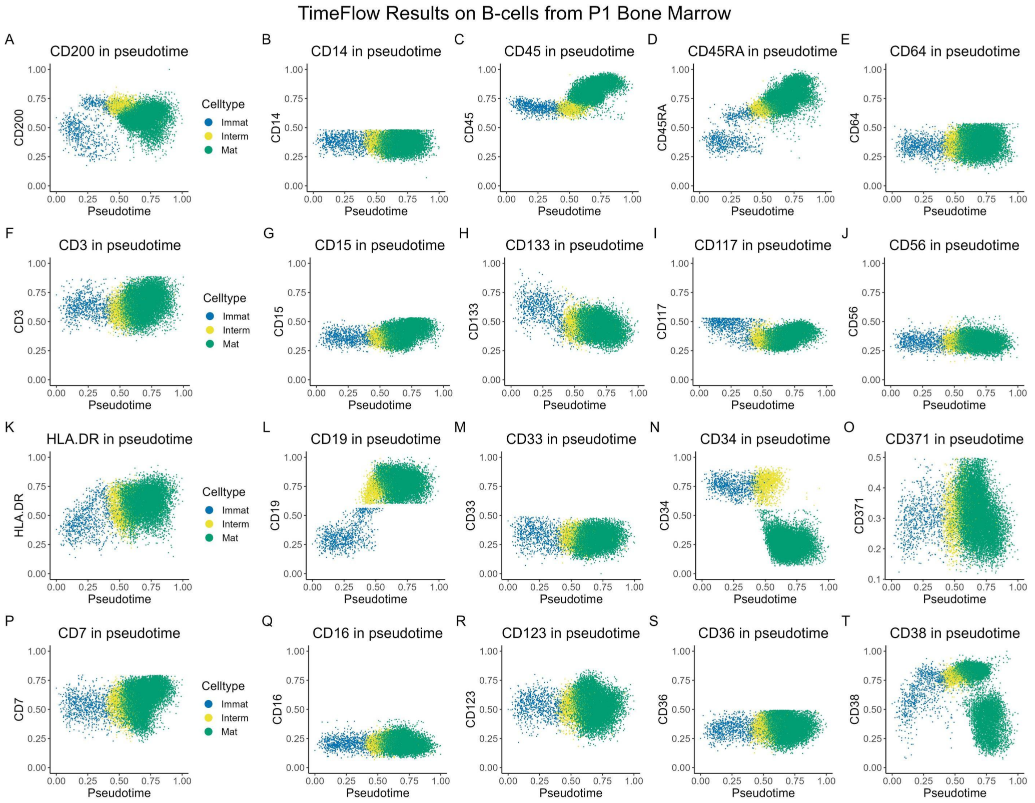


Figure S5: TimeFlow pseudotemporal orderings of B-cells using the P1-BM dataset. (A-T) Scatterplots showing the expression levels of each marker along pseudotime. Each scatterplot dot represents a cell, coloured by its maturation stage. Gaps in the marker expression (y-axis) between consecutive cell populations are a consequence of using rectangular gates during dataset preparation.

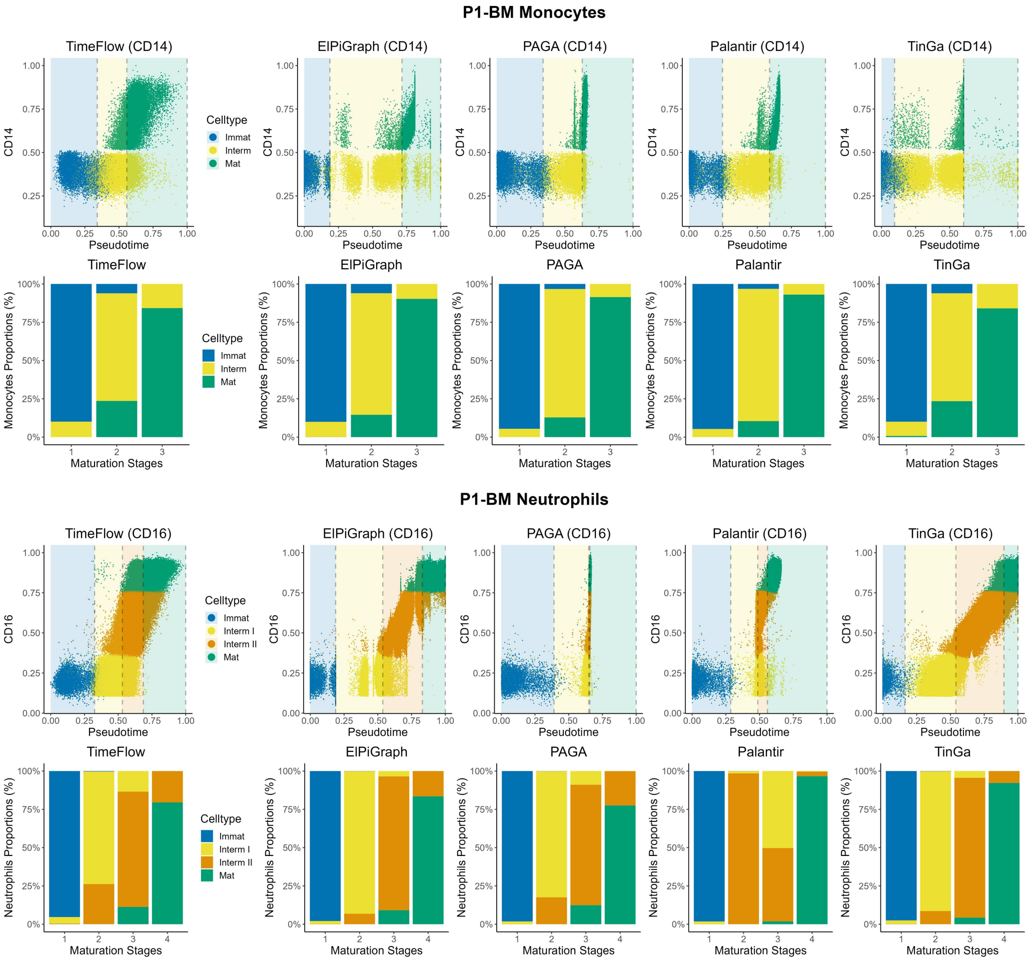


Figure S6: Methods comparisons on P1-BM Mono and P1-BM Neu trajectories. Each scatterplot dot represents a cell, coloured by its maturation stage. CD14 and CD16 were scaled in [0,1]. The vertical dashed grey lines indicate the boundaries between the cell stages. For each cell stage, the cut-off is defined at the pseudotime value corresponding to the last cell in that stage's ordered sequence (cumulative cell counts, see Section 2.3). Boundaries might overlap if a method does not separate between distinct stages. Regions correspond to different cell stages and are colored such that the transitions between them are highlighted. Stacked bar plots show the cell distribution (%) in each of their known maturation stages, following pseudotime ordering. The stacked barplots cut-offs are the same with the cut-offs on the pseudotime axis. The first two rows show the results for P1-BM Mono, while the last two rows present the results for P1-BM Neu based on TimeFlow, ElPiGraph, PAGA, Palantir, and TinGa. Gaps in the marker expression (y-axis) between consecutive cell populations are a consequence of using rectangular gates during dataset preparation.

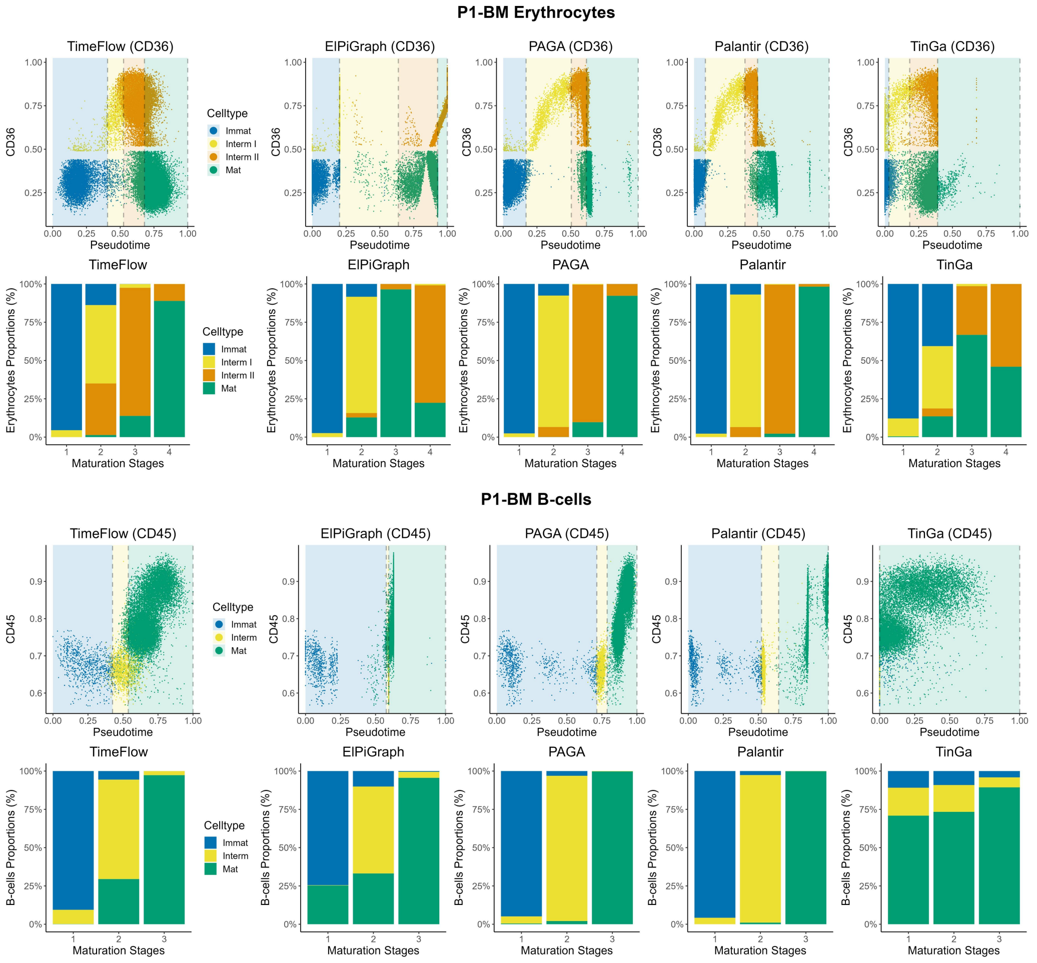


Figure S7: Methods comparisons on P1-BM Ery and P1-BM B-cells trajectories. Each scatterplot dot represents a cell, coloured by its maturation stage. CD36 and CD45 were scaled in [0,1]. The vertical dashed grey lines indicate the boundaries between the cell stages. For each cell stage, the cut-off is defined at the pseudotime value corresponding to the last cell in that stage's ordered sequence (cumulative cell counts, see Section 2.3). Boundaries might overlap if a method does not separate between distinct stages. Regions correspond to different cell stages and are colored such that the transitions between them are highlighted. Stacked bar plots show the cell distribution (%) in each of their known maturation stages, following pseudotime ordering. The stacked barplots cut-offs are the same with the cut-offs on the pseudotime axis. The first two rows show the results for P1-BM Ery, while the last two rows present the results for P1-BM B-cells based on TimeFlow, ElPiGraph, PAGA, Palantir, and TinGa. Gaps in the marker expression (y-axis) between consecutive cell populations are a consequence of using rectangular gates during dataset preparation.

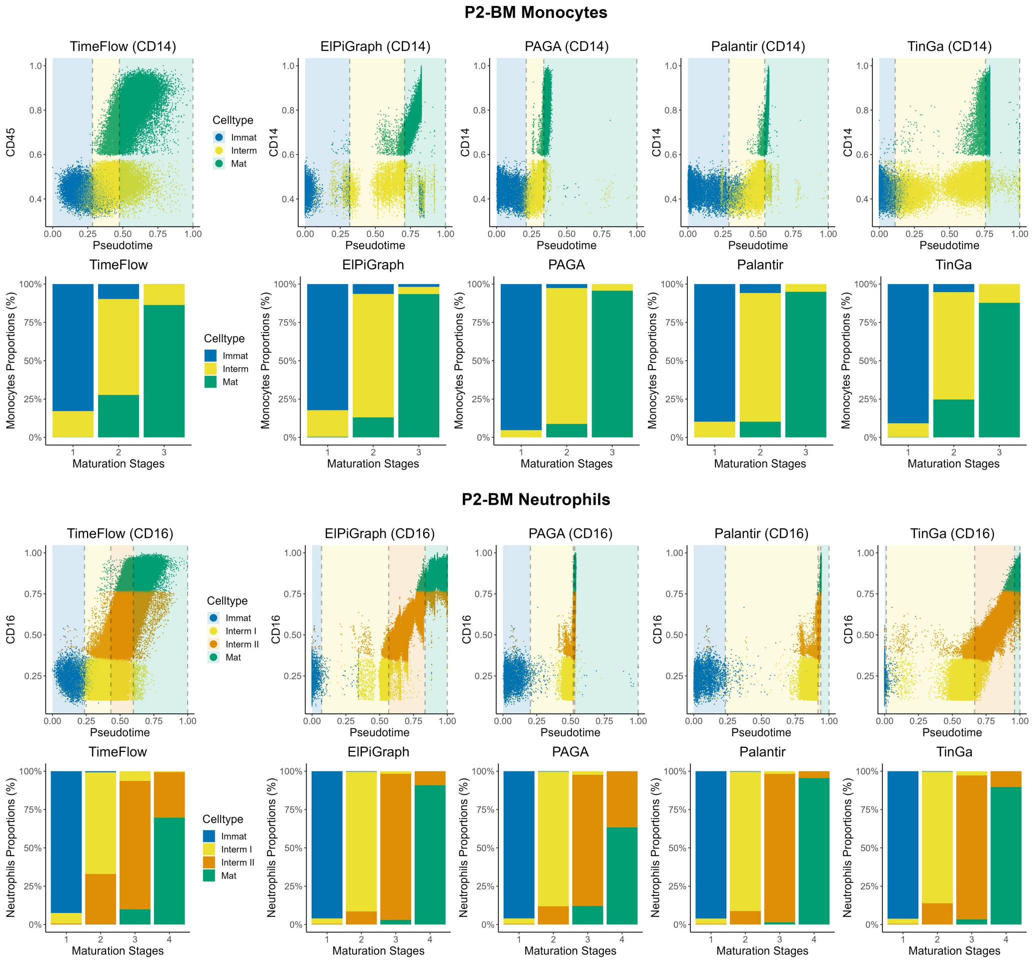


Figure S8: Methods comparisons on P2-BM Mono and P2-BM Neu trajectories. Each scatterplot dot represents a cell, coloured by its maturation stage. CD14 and CD16 were scaled in [0,1]. The vertical dashed grey lines indicate the boundaries between the cell stages. For each cell stage, the cut-off is defined at the pseudotime value corresponding to the last cell in that stage's ordered sequence (cumulative cell counts, see Section 2.3). Boundaries might overlap if a method does not separate between distinct stages. Regions correspond to different cell stages and are colored such that the transitions between them are highlighted. Stacked bar plots show the cell distribution (%) in each of their known maturation stages, following pseudotime ordering. The stacked barplots cut-offs are the same with the cut-offs on the pseudotime axis. The first two rows show the results for P2-BM Mono, while the last two rows present the results for P2-BM Neu based on TimeFlow, ElPiGraph, PAGA, Palantir, and TinGa. Gaps in the marker expression (y-axis) between consecutive cell populations are a consequence of using rectangular gates during dataset preparation.

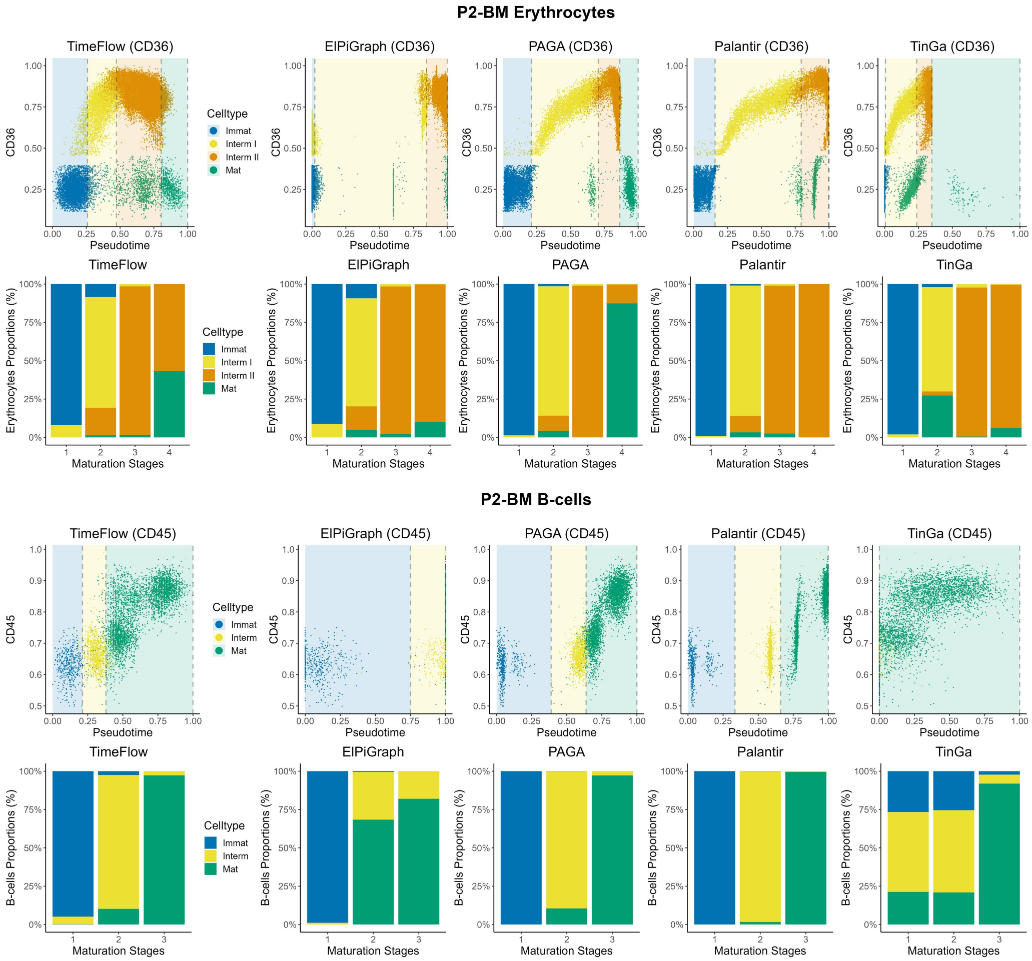


Figure S9: Methods comparisons on P2-BM Ery and P2-BM B-cells trajectories. Each scatterplot dot represents a cell, coloured by its maturation stage. CD36 and CD45 were scaled in [0,1]. The vertical dashed grey lines indicate the boundaries between the cell stages. For each cell stage, the cut-off is defined at the pseudotime value corresponding to the last cell in that stage's ordered sequence (cumulative cell counts, see Section 2.3). Boundaries might overlap if a method does not separate between distinct stages. Regions correspond to different cell stages and are colored such that the transitions between them are highlighted. Stacked bar plots show the cell distribution (%) in each of their known maturation stages, following pseudotime ordering. The stacked barplots cut-offs are the same with the cut-offs on the pseudotime axis. The first two rows show the results for P2-BM Ery, while the last two rows present the results for P2-BM B-cells based on TimeFlow, ElPiGraph, PAGA, Palantir, and TinGa. Gaps in the marker expression (y-axis) between consecutive cell populations are a consequence of using rectangular gates during dataset preparation.

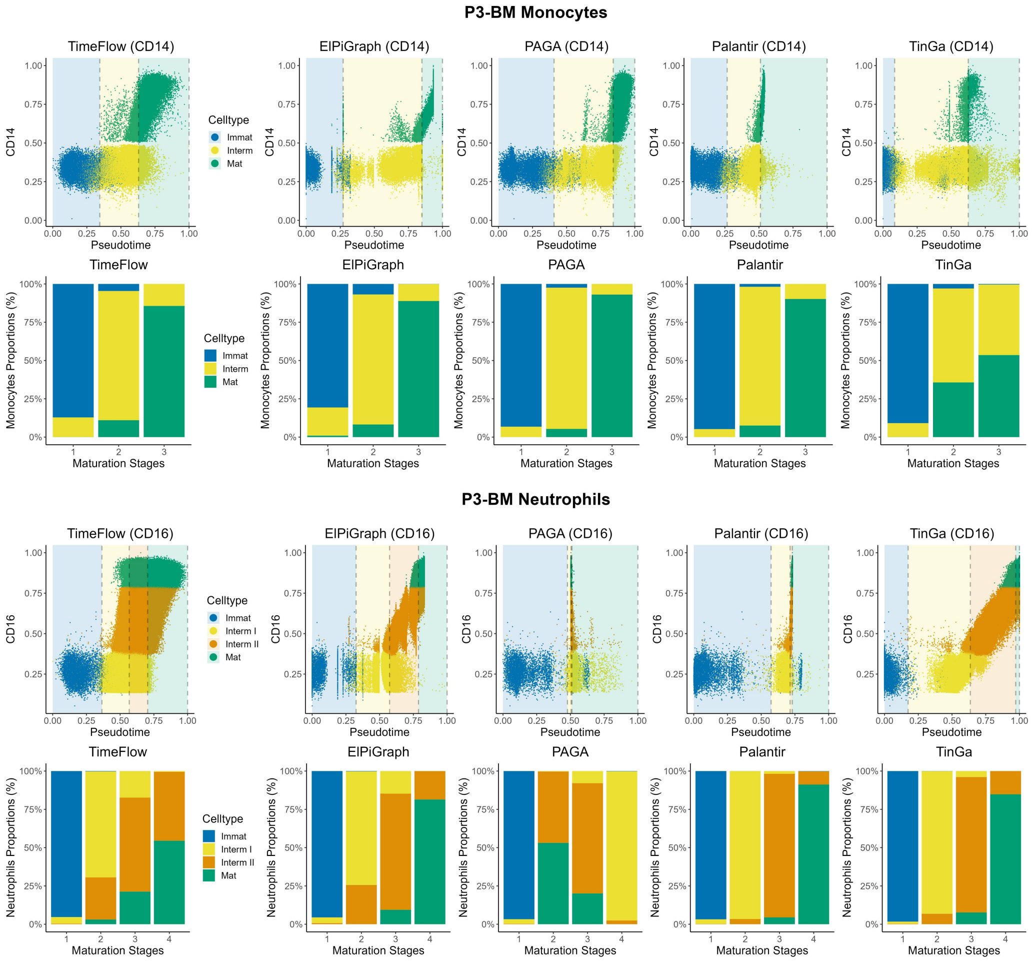


Figure S10: Methods comparisons on P3-BM Mono and P3-BM Neu trajectories. Each scatterplot dot represents a cell, coloured by its maturation stage. CD14 and CD16 were scaled in [0,1]. The vertical dashed grey lines indicate the boundaries between the cell stages. For each cell stage, the cut-off is defined at the pseudotime value corresponding to the last cell in that stage's ordered sequence (cumulative cell counts, see Section 2.3). Boundaries might overlap if a method does not separate between distinct stages. Regions correspond to different cell stages and are colored such that the transitions between them are highlighted. Stacked bar plots show the cell distribution (%) in each of their known maturation stages, following pseudotime ordering. The stacked barplots cut-offs are the same with the cut-offs on the pseudotime axis. The first two rows show the results for P3-BM Mono, while the last two rows present the results for P3-BM Neu based on TimeFlow, ElPiGraph, PAGA, Palantir, and TinGa. Gaps in the marker expression (y-axis) between consecutive cell populations are a consequence of using rectangular gates during dataset preparation.

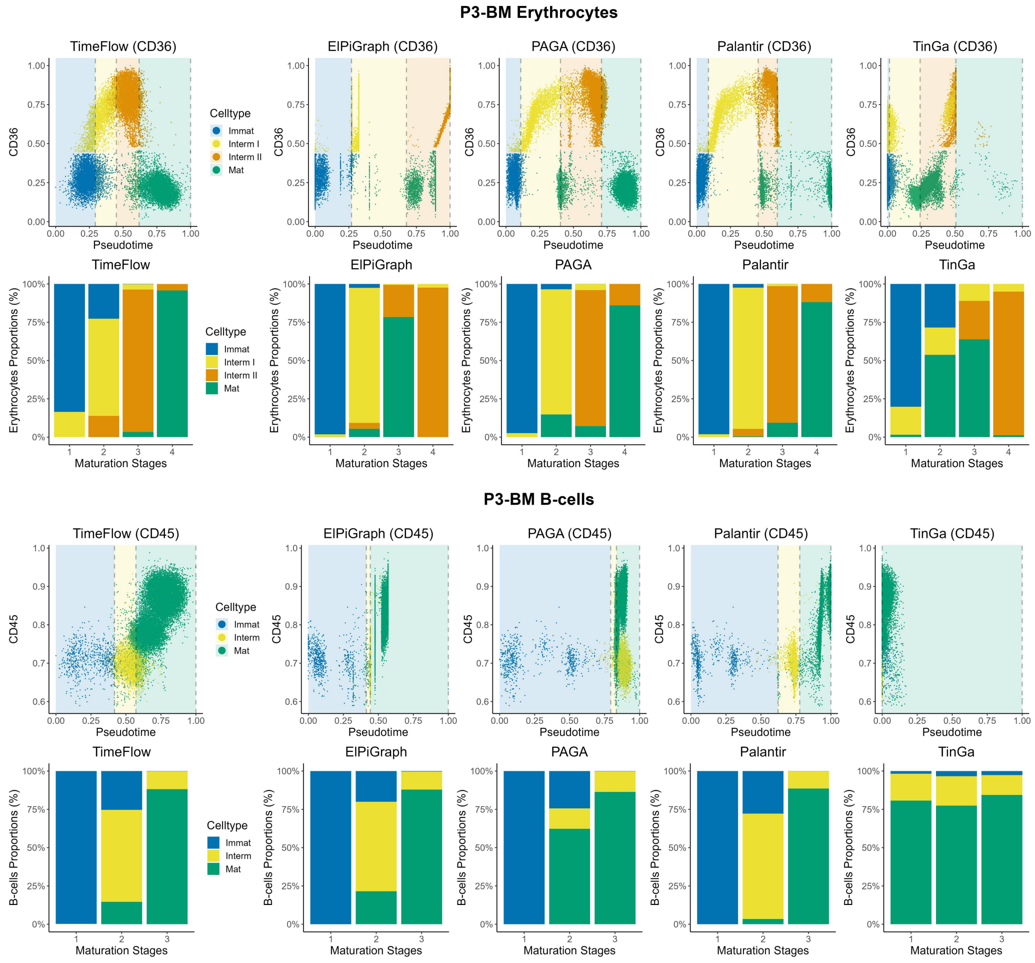


Figure S11: Methods comparisons on P3-BM Ery and P3-BM B-cells trajectories. Each scatterplot dot represents a cell, coloured by its maturation stage. CD36 and CD45 were scaled in [0,1]. The vertical dashed grey lines indicate the boundaries between the cell stages. For each cell stage, the cut-off is defined at the pseudotime value corresponding to the last cell in that stage's ordered sequence (cumulative cell counts, see Section 2.3). Boundaries might overlap if a method does not separate between distinct stages. Regions correspond to different cell stages and are colored such that the transitions between them are highlighted. Stacked bar plots show the cell distribution (%) in each of their known maturation stages, following pseudotime ordering. The stacked barplots cut-offs are the same with the cut-offs on the pseudotime axis. The first two rows show the results for P3-BM Ery, while the last two rows present the results for P3-BM B-cells based on TimeFlow, ElPiGraph, PAGA, Palantir, and TinGa. Gaps in the marker expression (y-axis) between consecutive cell populations are a consequence of using rectangular gates during dataset preparation.

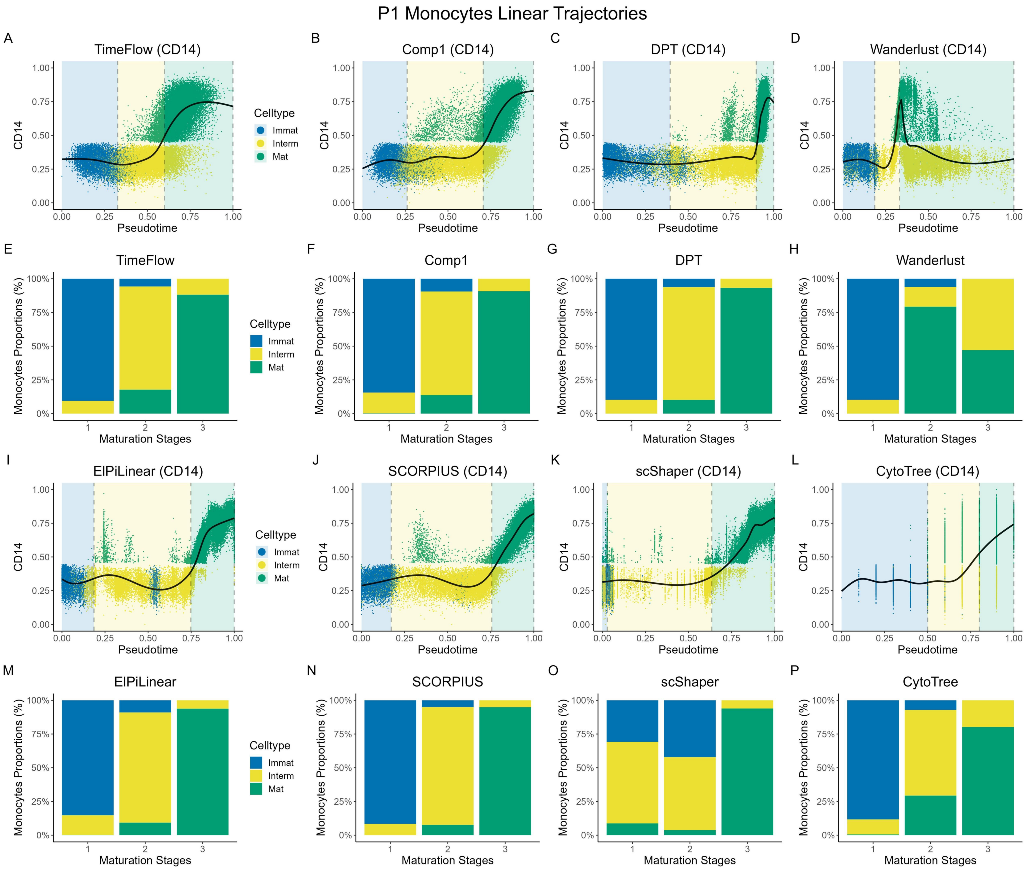


Figure S12: Methods comparisons on linear P1 Mono trajectory. Each scatterplot dot represents a cell, coloured by its maturation stage. CD14 was scaled in [0,1]. A GAM model (solid black curve) was fitted without use of labels. The vertical dashed grey lines indicate the boundaries between the cell stages. For each cell stage, the cut-off is defined at the pseudotime value corresponding to the last cell in that stage's ordered sequence (cumulative cell counts, see Section 2.3). Boundaries might overlap if a method does not separate between distinct stages. Regions correspond to different cell stages and are colored such that the transitions between them are highlighted. Stacked bar plots show the cell distribution (%) in each of their known maturation stages, following pseudotime ordering. The stacked barplots cut-offs are the same with the cut-offs on the pseudotime axis. (A-P) Results for TimeFlow, Comp1, DPT, Wanderlust, ElPiLinear, SCORPIUS, scShaper, and CytoTree. Gaps in the marker expression (y-axis) between consecutive cell populations are a consequence of using rectangular gates during dataset preparation.

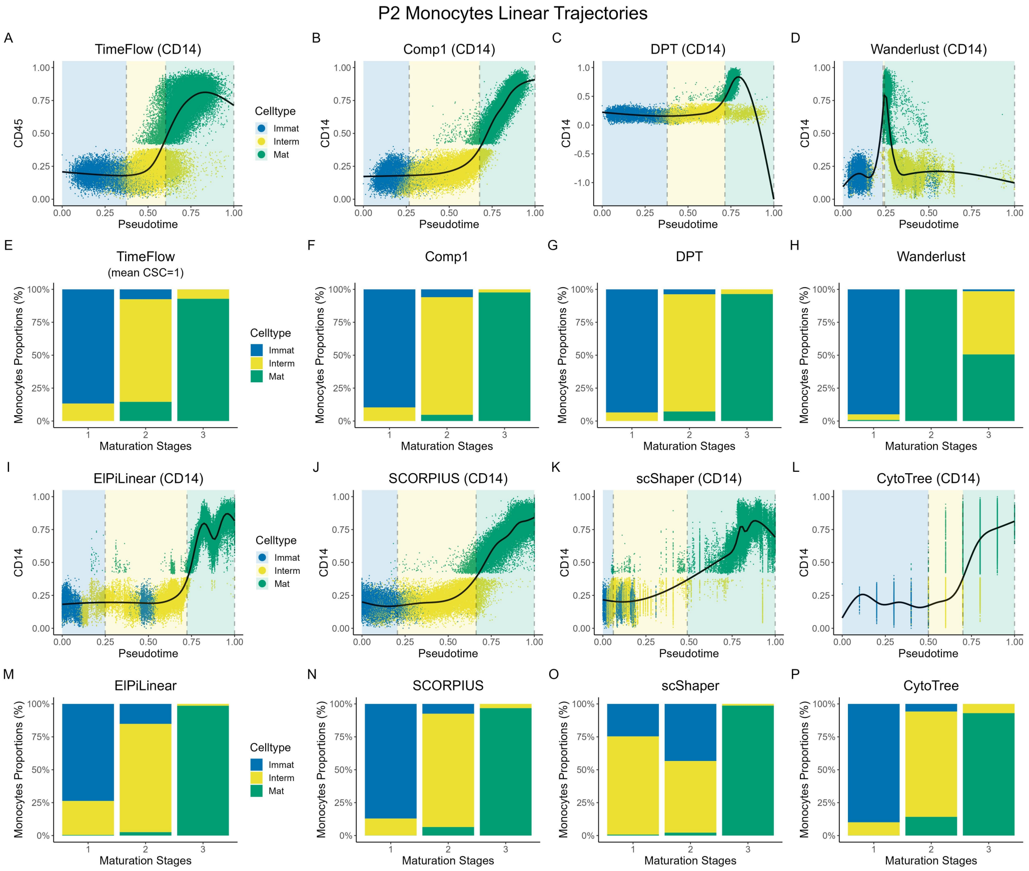


Figure S13: Methods comparisons on linear P2 Mono trajectory. Each scatterplot dot represents a cell, coloured by its maturation stage. CD14 was scaled in [0,1]. A GAM model (solid black curve) was fitted without use of labels. The vertical dashed grey lines indicate the boundaries between the cell stages. For each cell stage, the cut-off is defined at the pseudotime value corresponding to the last cell in that stage's ordered sequence (cumulative cell counts, see Section 2.3). Boundaries might overlap if a method does not separate between distinct stages. Regions correspond to different cell stages and are colored such that the transitions between them are highlighted. Stacked bar plots show the cell distribution (%) in each of their known maturation stages, following pseudotime ordering. The stacked barplots cut-offs are the same with the cut-offs on the pseudotime axis. (A-P) Results for TimeFlow, Comp1, DPT, Wanderlust, ElPiLinear, SCORPIUS, scShaper, and CytoTree. Gaps in the marker expression (y-axis) between consecutive cell populations are a consequence of using rectangular gates during dataset preparation.

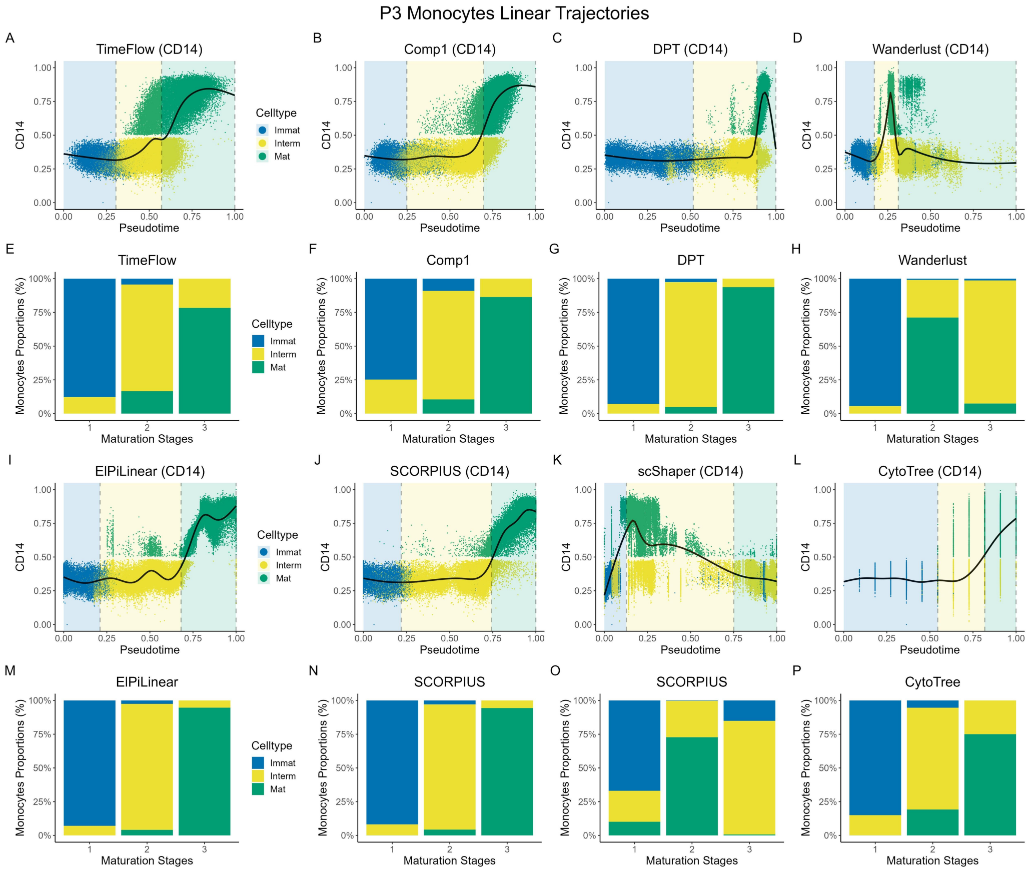


Figure S14: Methods comparisons on linear P3 Mono trajectory. Each scatterplot dot represents a cell, coloured by its maturation stage. CD14 was scaled in [0,1]. A GAM model (solid black curve) was fitted without use of labels. The vertical dashed grey lines indicate the boundaries between the cell stages. For each cell stage, the cut-off is defined at the pseudotime value corresponding to the last cell in that stage's ordered sequence (cumulative cell counts, see Section 2.3). Boundaries might overlap if a method does not separate between distinct stages. Regions correspond to different cell stages and are colored such that the transitions between them are highlighted. Stacked bar plots show the cell distribution (%) in each of their known maturation stages, following pseudotime ordering. The stacked barplots cut-offs are the same with the cut-offs on the pseudotime axis. (A-P) Results for TimeFlow, Comp1, DPT, Wanderlust, ElPiLinear, SCORPIUS, scShaper, and CytoTree. Gaps in the marker expression (y-axis) between consecutive cell populations are a consequence of using rectangular gates during dataset preparation.

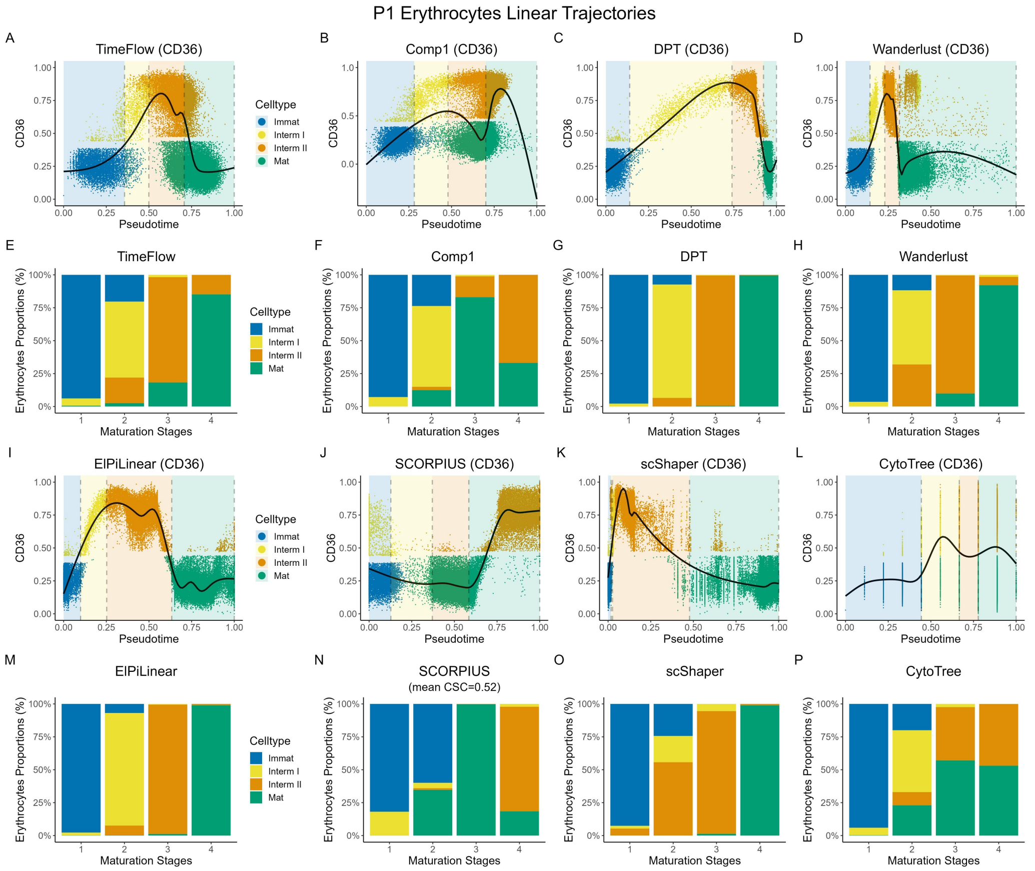


Figure S15: Methods comparisons on linear P1 Ery trajectory. Each scatterplot dot represents a cell, coloured by its maturation stage. CD36 was scaled in [0,1]. A GAM model (solid black curve) was fitted without use of labels. The vertical dashed grey lines indicate the boundaries between the cell stages. For each cell stage, the cut-off is defined at the pseudotime value corresponding to the last cell in that stage's ordered sequence (cumulative cell counts, see Section 2.3). Boundaries might overlap if a method does not separate between distinct stages. Regions correspond to different cell stages and are colored such that the transitions between them are highlighted. Stacked bar plots show the cell distribution (%) in each of their known maturation stages, following pseudotime ordering. The stacked barplots cut-offs are the same with the cut-offs on the pseudotime axis. (A-P) Results for TimeFlow, Comp1, DPT, Wanderlust, ElPiLinear, SCORPIUS, scShaper, and CytoTree. Gaps in the marker expression (y-axis) between consecutive cell populations are a consequence of using rectangular gates during dataset preparation.

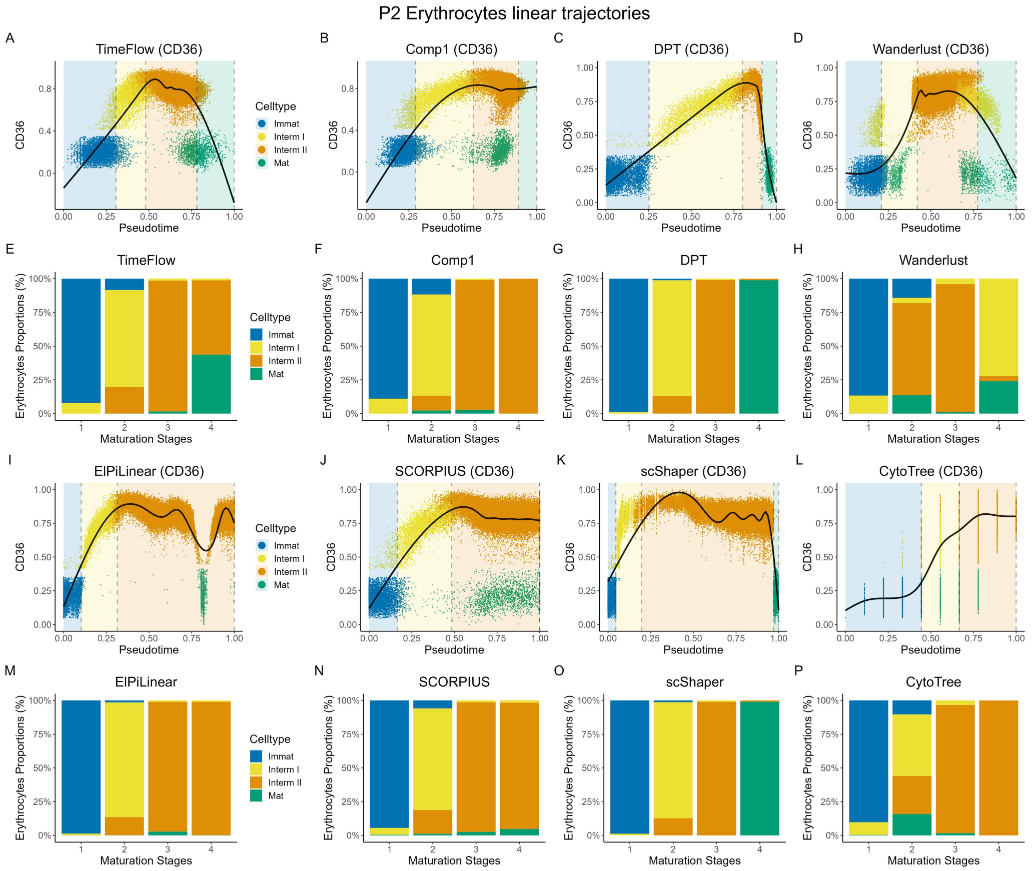


Figure S16: Methods comparisons on linear P2 Ery trajectory. Each scatterplot dot represents a cell, coloured by its maturation stage. CD36 was scaled in [0,1]. A GAM model (solid black curve) was fitted without use of labels. The vertical dashed grey lines indicate the boundaries between the cell stages. For each cell stage, the cut-off is defined at the pseudotime value corresponding to the last cell in that stage's ordered sequence (cumulative cell counts, see Section 2.3). Boundaries might overlap if a method does not separate between distinct stages. Regions correspond to different cell stages and are colored such that the transitions between them are highlighted. Stacked bar plots show the cell distribution (%) in each of their known maturation stages, following pseudotime ordering. The stacked barplots cut-offs are the same with the cut-offs on the pseudotime axis. (A-P) Results for TimeFlow, Comp1, DPT, Wanderlust, ElPiLinear, SCORPIUS, scShaper, and CytoTree. Gaps in the marker expression (y-axis) between consecutive cell populations are a consequence of using rectangular gates during dataset preparation.

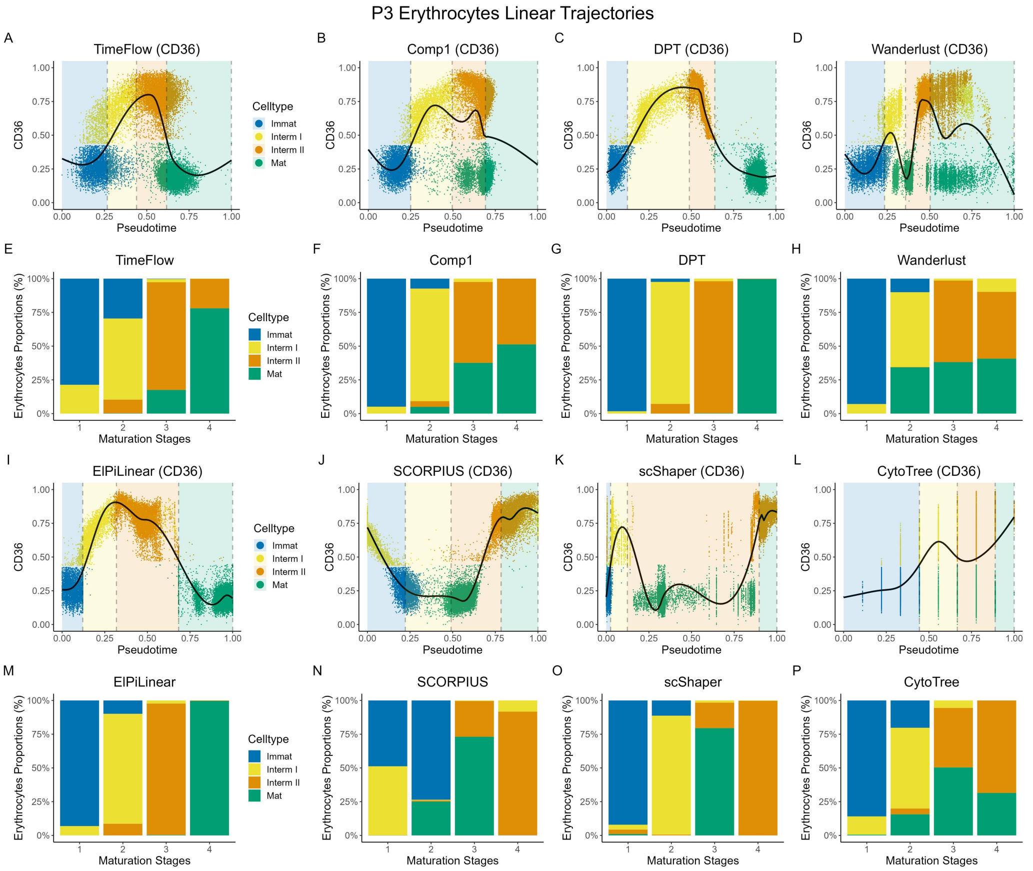


Figure S17: Methods comparisons on linear P3 Ery trajectory. Each scatterplot dot represents a cell, coloured by its maturation stage. CD36 was scaled in [0,1]. A GAM model (solid black curve) was fitted without use of labels. The vertical dashed grey lines indicate the boundaries between the cell stages. For each cell stage, the cut-off is defined at the pseudotime value corresponding to the last cell in that stage's ordered sequence (cumulative cell counts, see Section 2.3). Boundaries might overlap if a method does not separate between distinct stages. Regions correspond to different cell stages and are colored such that the transitions between them are highlighted. Stacked bar plots show the cell distribution (%) in each of their known maturation stages, following pseudotime ordering. The stacked barplots cut-offs are the same with the cut-offs on the pseudotime axis. (A-P) Results for TimeFlow, Comp1, DPT, Wanderlust, ElPiLinear, SCORPIUS, scShaper, and CytoTree. Gaps in the marker expression (y-axis) between consecutive cell populations are a consequence of using rectangular gates during dataset preparation.

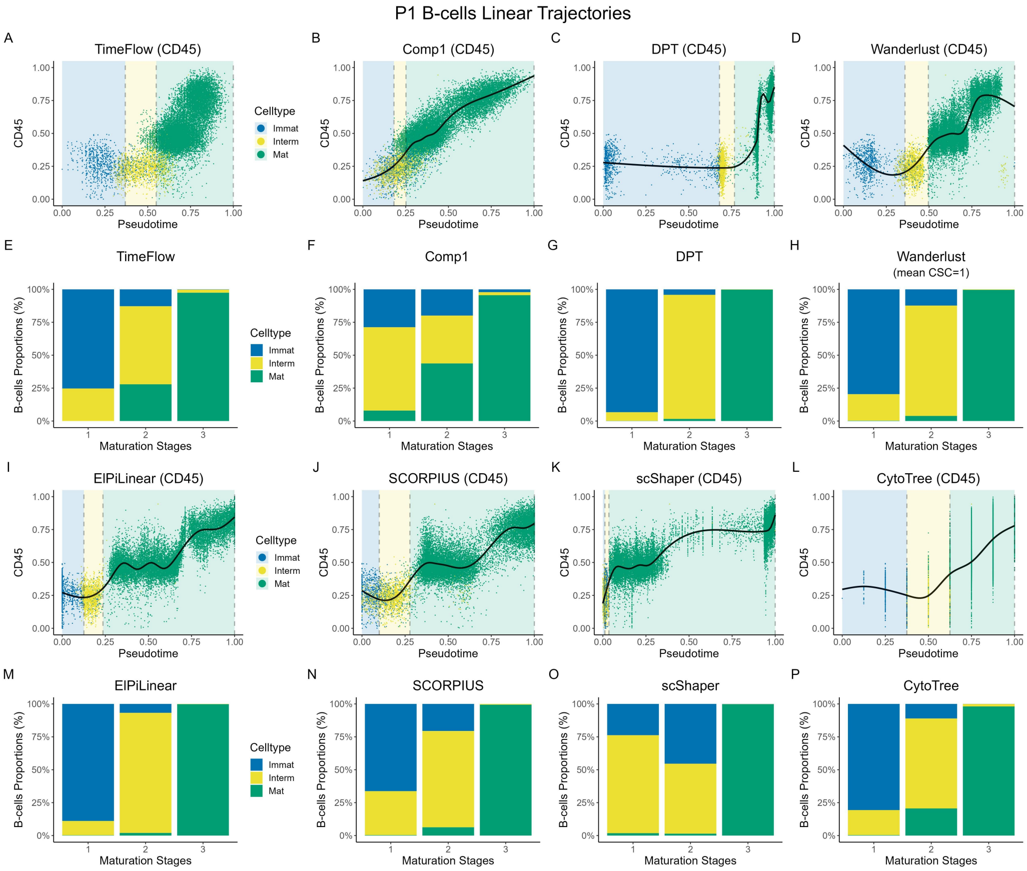


Figure S18: Methods comparisons on linear P1 B-cells trajectory. Each scatterplot dot represents a cell, coloured by its maturation stage. CD45 was scaled in [0,1]. A GAM model (solid black curve) was fitted without use of labels. The vertical dashed grey lines indicate the boundaries between the cell stages. For each cell stage, the cut-off is defined at the pseudotime value corresponding to the last cell in that stage's ordered sequence (cumulative cell counts, see Section 2.3). Boundaries might overlap if a method does not separate between distinct stages. Regions correspond to different cell stages and are colored such that the transitions between them are highlighted. Stacked bar plots show the cell distribution (%) in each of their known maturation stages, following pseudotime ordering. The stacked barplots cut-offs are the same with the cut-offs on the pseudotime axis. (A-P) Results for TimeFlow, Comp1, DPT, Wanderlust, ElPiLinear, SCORPIUS, scShaper, and CytoTree. Gaps in the marker expression (y-axis) between consecutive cell populations are a consequence of using rectangular gates during dataset preparation.

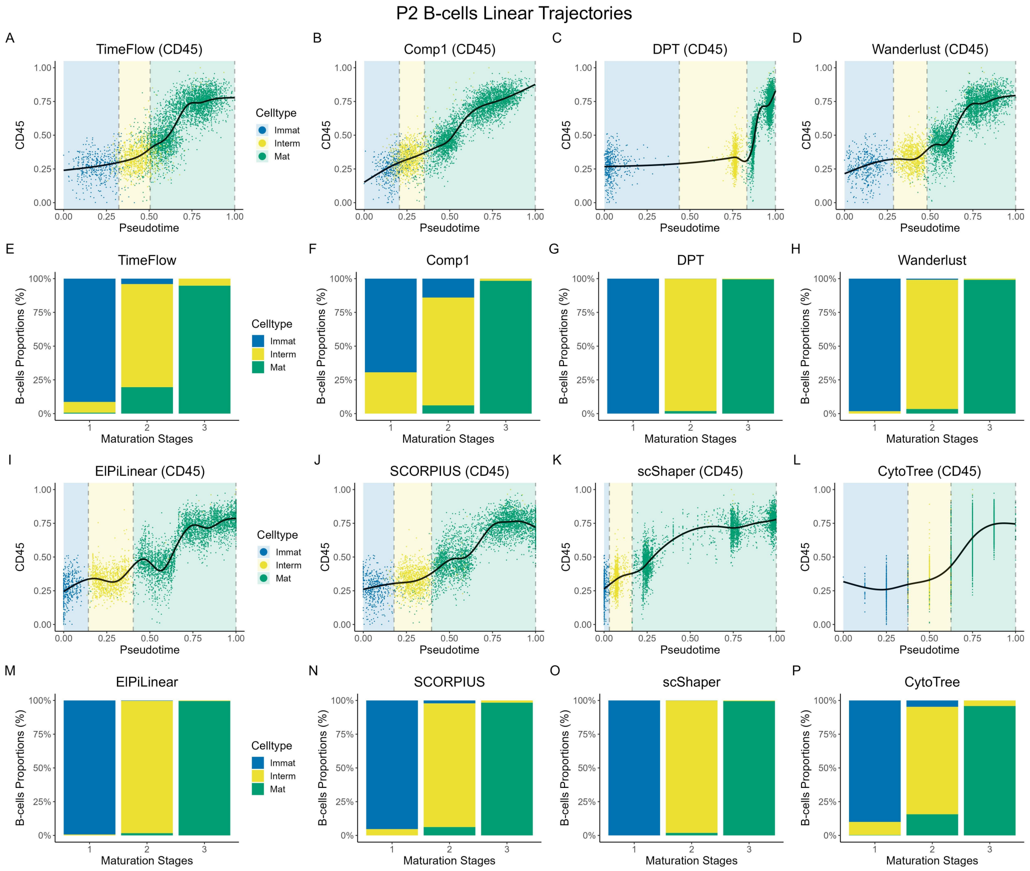


Figure S19: Methods comparisons on linear P2 B-cells trajectory. Each scatterplot dot represents a cell, coloured by its maturation stage. CD45 was scaled in [0,1]. A GAM model (solid black curve) was fitted without use of labels. The vertical dashed grey lines indicate the boundaries between the cell stages. For each cell stage, the cut-off is defined at the pseudotime value corresponding to the last cell in that stage's ordered sequence (cumulative cell counts, see Section 2.3). Boundaries might overlap if a method does not separate between distinct stages. Regions correspond to different cell stages and are colored such that the transitions between them are highlighted. Stacked bar plots show the cell distribution (%) in each of their known maturation stages, following pseudotime ordering. The stacked barplots cut-offs are the same with the cut-offs on the pseudotime axis. (A-P) Results for TimeFlow, Comp1, DPT, Wanderlust, ElPiLinear, SCORPIUS, scShaper, and CytoTree. Gaps in the marker expression (y-axis) between consecutive cell populations are a consequence of using rectangular gates during dataset preparation.

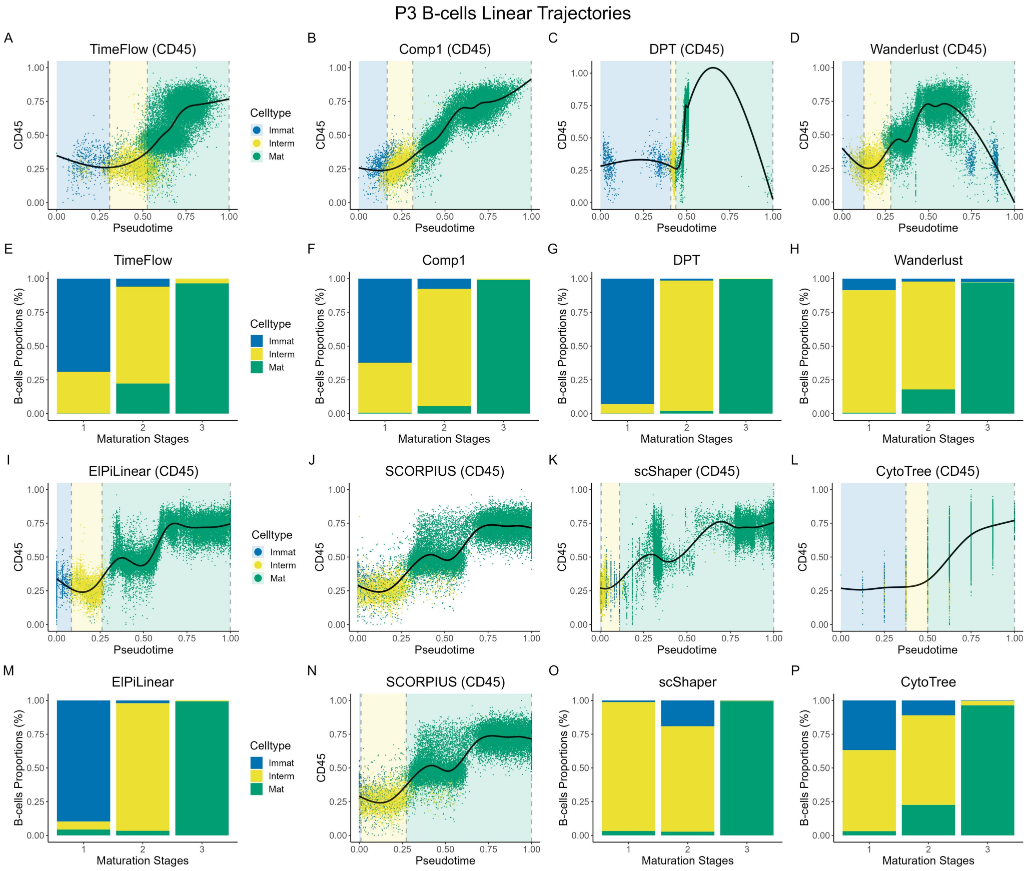


Figure S20: Methods comparisons on linear P3 B-cells trajectory. Each scatterplot dot represents a cell, coloured by its maturation stage. CD45 was scaled in [0,1]. A GAM model (solid black curve) was fitted without use of labels. The vertical dashed grey lines indicate the boundaries between the cell stages. For each cell stage, the cut-off is defined at the pseudotime value corresponding to the last cell in that stage's ordered sequence (cumulative cell counts, see Section 2.3). Boundaries might overlap if a method does not separate between distinct stages. Regions correspond to different cell stages and are colored such that the transitions between them are highlighted. Stacked bar plots show the cell distribution (%) in each of their known maturation stages, following pseudotime ordering. The stacked barplots cut-offs are the same with the cut-offs on the pseudotime axis. (A-P) Results for TimeFlow, Comp1, DPT, Wanderlust, ElPiLinear, SCORPIUS, scShaper, and CytoTree. Gaps in the marker expression (y-axis) between consecutive cell populations are a consequence of using rectangular gates during dataset preparation.

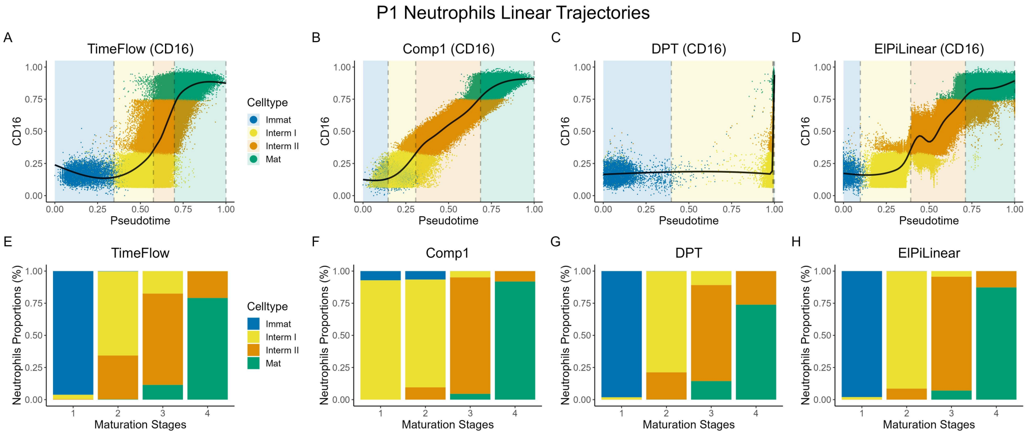


Figure S21: Methods comparisons on linear P1 Neutrophils trajectory. Each scatterplot dot represents a cell, coloured by its maturation stage. CD16 was scaled in [0,1]. A GAM model (solid black curve) was fitted without use of labels. The vertical dashed grey lines indicate the boundaries between the cell stages. For each cell stage, the cut-off is defined at the pseudotime value corresponding to the last cell in that stage's ordered sequence (cumulative cell counts, see Section 2.3). Boundaries might overlap if a method does not separate between distinct stages. Regions correspond to different cell stages and are colored such that the transitions between them are highlighted. Stacked bar plots show the cell distribution (%) in each of their known maturation stages, following pseudotime ordering. The stacked barplots cut-offs are the same with the cut-offs on the pseudotime axis. (A-H) Results for TimeFlow, Comp1, DPT, and ElPiLinear. Gaps in the marker expression (y-axis) between consecutive cell populations are a consequence of using rectangular gates during dataset preparation.

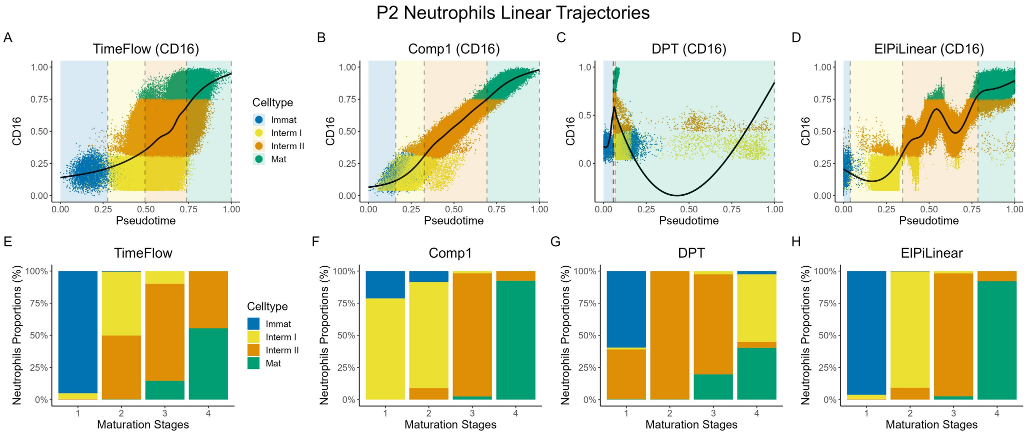


Figure S22: Methods comparisons on linear P2 Neutrophils trajectory. Each scatterplot dot represents a cell, coloured by its maturation stage. CD16 was scaled in [0,1]. A GAM model (solid black curve) was fitted without use of labels. The vertical dashed grey lines indicate the boundaries between the cell stages. For each cell stage, the cut-off is defined at the pseudotime value corresponding to the last cell in that stage's ordered sequence (cumulative cell counts, see Section 2.3). Boundaries might overlap if a method does not separate between distinct stages. Regions correspond to different cell stages and are colored such that the transitions between them are highlighted. Stacked bar plots show the cell distribution (%) in each of their known maturation stages, following pseudotime ordering. The stacked barplots cut-offs are the same with the cut-offs on the pseudotime axis. (A-H) Results for TimeFlow, Comp1, DPT, and ElPiLinear. Gaps in the marker expression (y-axis) between consecutive cell populations are a consequence of using rectangular gates during dataset preparation.

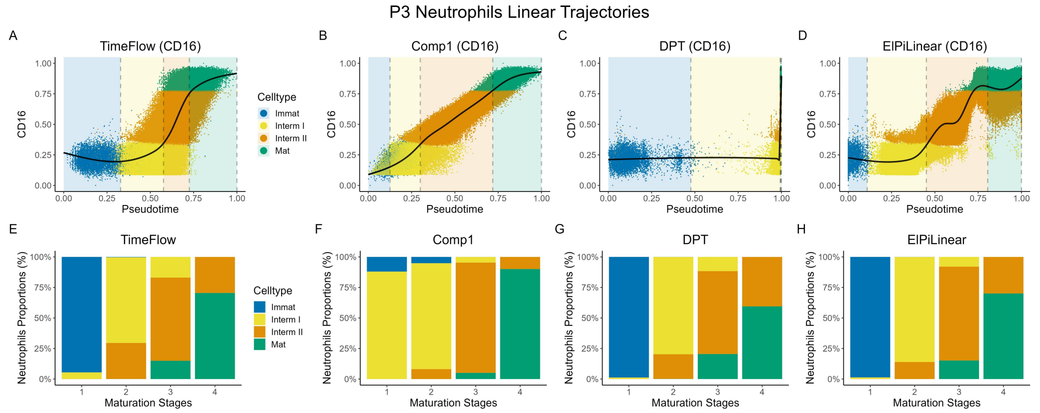


Figure S23: Methods comparisons on linear P3 Neutrophils trajectory. Each scatterplot dot represents a cell, coloured by its maturation stage. CD16 was scaled in [0,1]. A GAM model (solid black curve) was fitted without use of labels. The vertical dashed grey lines indicate the boundaries between the cell stages. For each cell stage, the cut-off is defined at the pseudotime value corresponding to the last cell in that stage's ordered sequence (cumulative cell counts, see Section 2.3). Boundaries might overlap if a method does not separate between distinct stages. Regions correspond to different cell stages and are colored such that the transitions between them are highlighted. Stacked bar plots show the cell distribution (%) in each of their known maturation stages, following pseudotime ordering. The stacked barplots cut-offs are the same with the cut-offs on the pseudotime axis. (A-H) Results for TimeFlow, Comp1, DPT, and ElPiLinear. Gaps in the marker expression (y-axis) between consecutive cell populations are a consequence of using rectangular gates during dataset preparation.

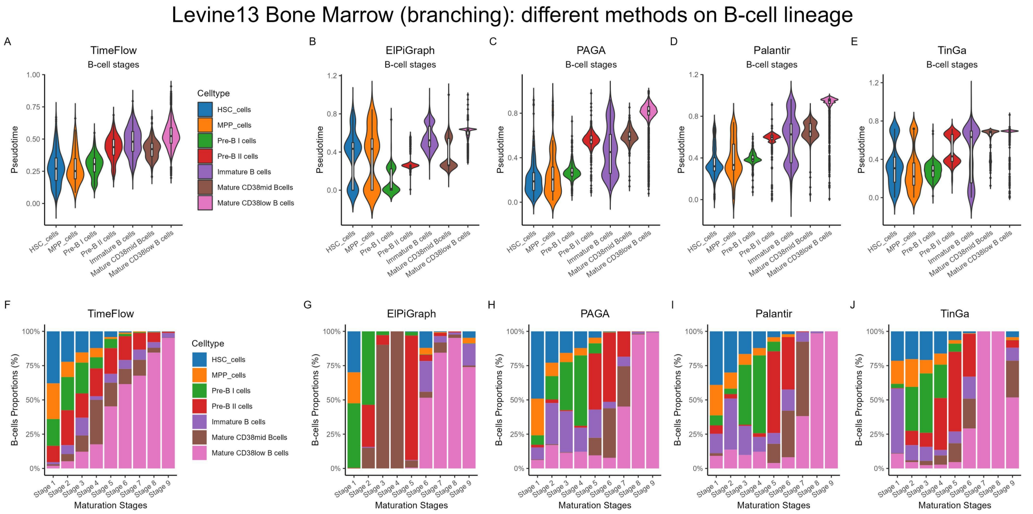


Figure S24: Side-by-side methods comparison on B-cell lineage of the public Levine-13 dataset (branching trajectory). Each scatterplot dot represents a cell, coloured by its maturation stage. Violin plots with boxplots are ordered based on the expected stage progress of B-cell differentiation and show the pseudotime distribution for each cell population. Boxplots show the median, quartiles, minimum, and maximum. (A-E) Violin plots with boxplots show the distribution of pseudotime, inferred by TimeFlow, ElPiGraph, PAGA, Palantir and TinGa, across cell populations (F-J) Stacked bar plots show the cell distribution (%) in each of their known maturation stages, following pseudotime ordering.

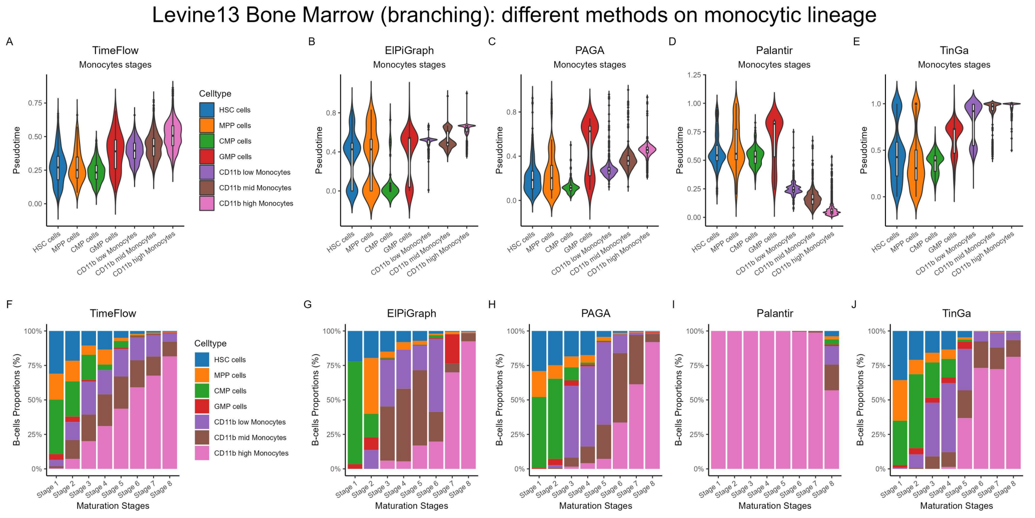


Figure S25: Side-by-side methods comparison on monocytic lineage of the public Levine-13 dataset (branching trajectory). Each scatterplot dot represents a cell, coloured by its maturation stage. Violin plots with boxplots are ordered based on the expected stage progress of monocytic differentiation and show the pseudotime distribution for each cell population. Boxplots show the median, quartiles, minimum, and maximum. (A-E) Violin plots with boxplots show the distribution of pseudotime, inferred by TimeFlow, ElPiGraph, PAGA, Palantir and TinGa, across cell populations (F-J) Stacked bar plots show the cell distribution (%) in each of their known maturation stages, following pseudotime ordering.

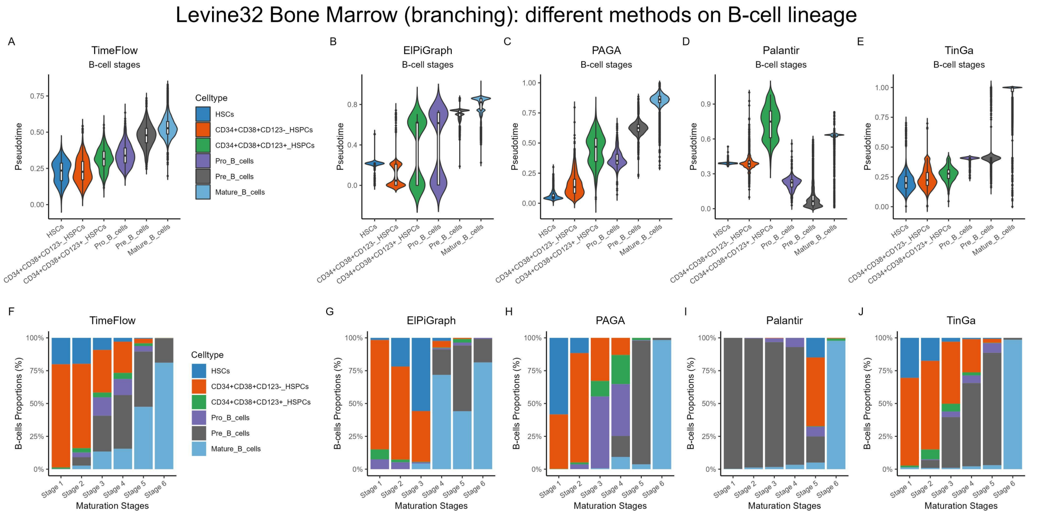


Figure S26: Side-by-side methods comparison on B-cell lineage of the public Levine-32 dataset (branching trajectory). Violin plots with boxplots are ordered based on the expected stage progress of B-cell differentiation and show the pseudotime distribution for each cell population. Boxplots show median, quartiles, minima, maxima. (A-E) Violin plots with boxplots show the distribution of pseudotime, inferred by TimeFlow, ElPiGraph, PAGA, Palantir and TinGa, across cell populations. (F-J) Stacked bar plots show the cell distribution (%) in each of their known maturation stages, following pseudotime ordering.

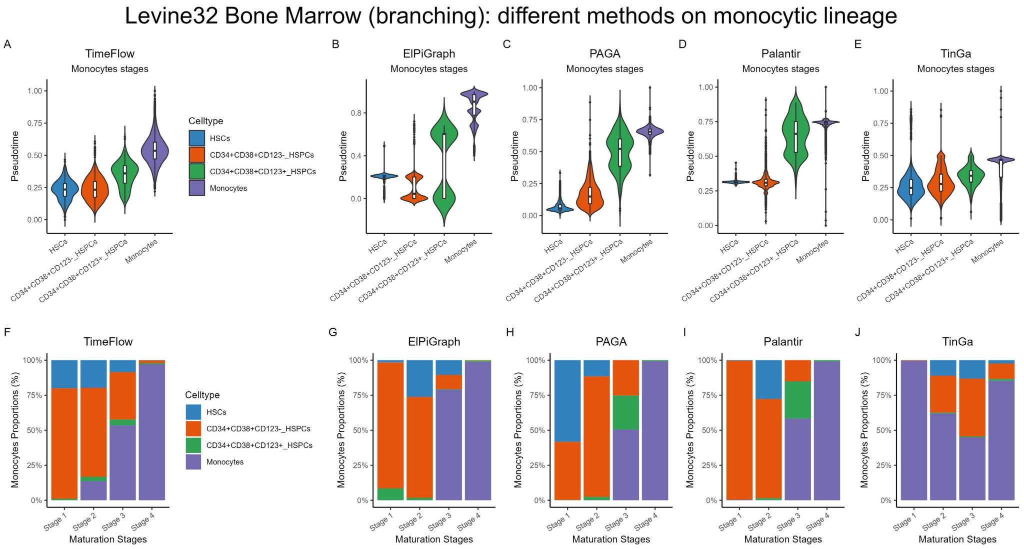


Figure S27: Side-by-side methods comparison on monocytic lineage of the public Levine-32 dataset (branching trajectory). Violin plots with boxplots are ordered based on the expected stage progress of monocytic differentiation and show the pseudotime distribution for each cell population. Boxplots show median, quartiles, minima, maxima. (A-E) Violin plots with boxplots show the distribution of pseudotime, inferred by TimeFlow, ElPiGraph, PAGA, Palantir and TinGa, across cell populations. (F-J) Stacked bar plots show the cell distribution (%) in each of their known maturation stages, following pseudotime ordering.

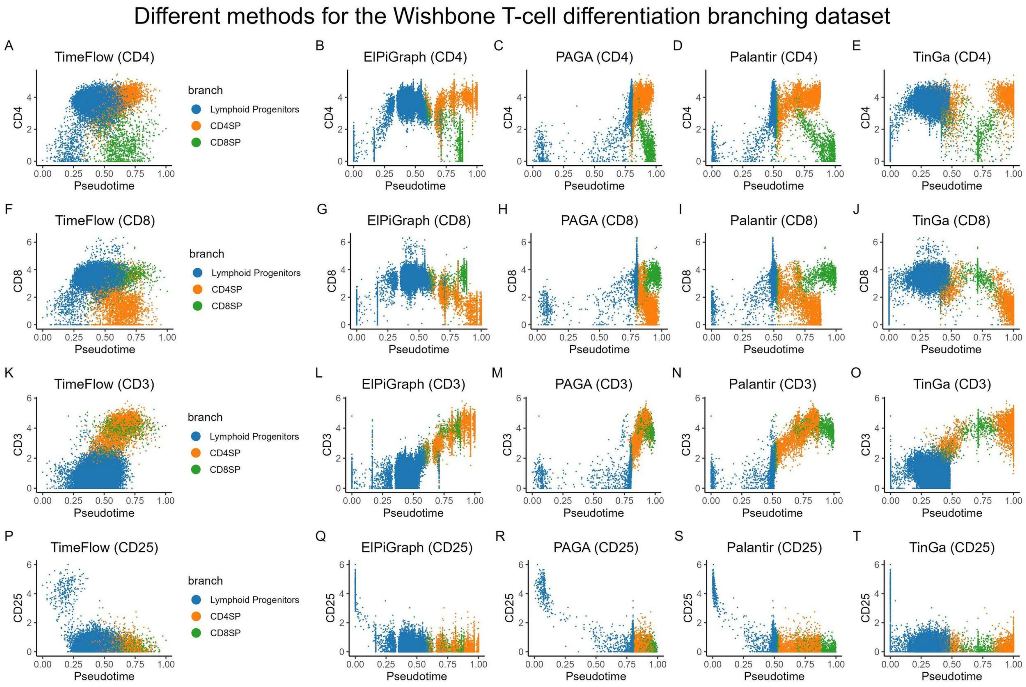


Figure S28: Side-by-side methods comparison on $CD4^+$ SP and $CD8^+$ SP lineages of the public Wishbone dataset (branching trajectory). Each scatterplot dot represents a cell, coloured by its maturation stage. (A-E) CD4 expression (CD4SP-specific lineage marker) for all cells after pseudotime ordering based on TimeFlow, ElPiGraph, PAGA, Palantir and TinGa. (F-J) CD8 expression (CD8SP-specific lineage marker) for all cells after pseudotime ordering. (K-O) CD3 expression for all cells sorted by pseudotime (marker with known increasing expression). (P-T) CD25 expression for all cells sorted by pseudotime (marker with known decreasing expression).

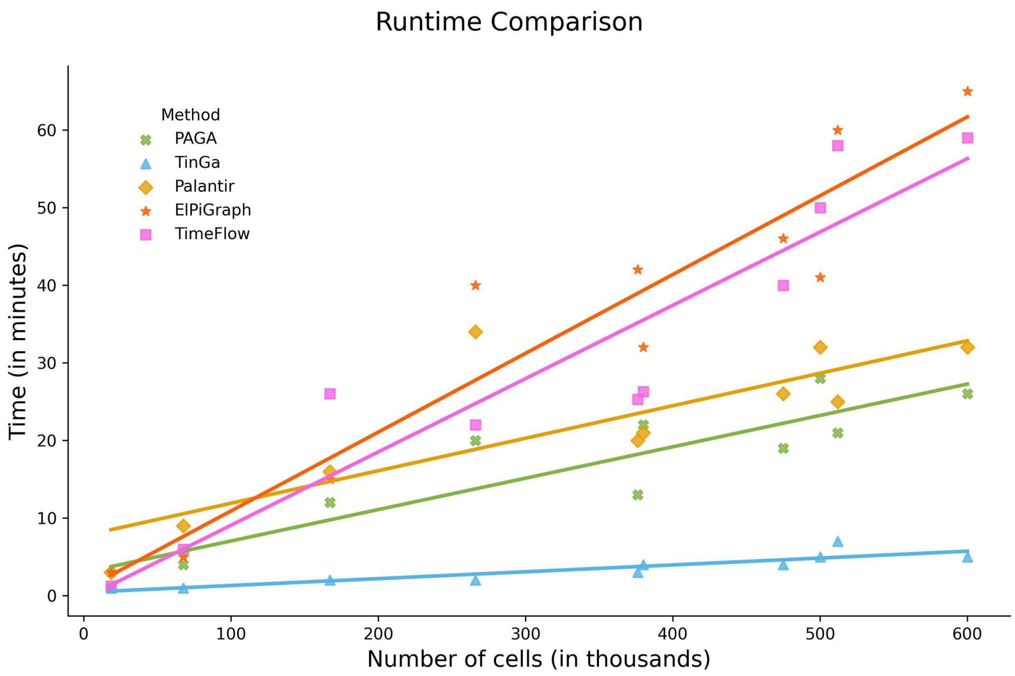


Figure S29: Runtime comparison for TimeFlow, PAGA, Palantir, TinGa and ElPiGraph using the largest datasets of this study (P1/2/3-BM, P1/2/3-Neu, Levine13, Levine32, and two randomly chosen smaller ones—P3 Mono and P1-Bcells). Differences in scalability are shown with regression lines. Time is measured in minutes and number of cells in thousands.
